# Organ-specific electrophile responsivity mapping in live *C. elegans*

**DOI:** 10.1101/2024.09.27.615266

**Authors:** Jinmin Liu, Amogh Kulkarni, Yong-Qi Gao, Daniel A. Urul, Romain Hamelin, Balázs Á. Novotny, Marcus J. C. Long, Yimon Aye

## Abstract

Proximity labeling technologies are limited to indexing localized protein residents. Such data— although valuable—cannot inform on small-molecule responsivity of local residents. We here bridge this gap by demonstrating in live *C. elegans* how electrophile-sensing propensity in specific organs can be quantitatively mapped and ranked. Using this method, >70% of tissue-specific responders exhibit novel responsivity, *independent* of tissue-specific abundance. One responder, cyp-33e1—for which both human and worm orthologs are electrophile responsive—marshals stress-dependent gut functions, despite manifesting uniform abundance across all tissues studied. Cyp-33e1’s localized electrophile responsivity operates site-specifically, triggering multifaceted responses: electrophile sensing through the catalytic-site cysteine results in partitioning between enzyme inhibition and localized production of a critical metabolite that governs global lipid availability, whereas a rapid dual-cysteine site-specific sensing modulates gut homeostasis. Beyond pinpointing chemical actionability within local proteomes, tissue-specific electrophile responsivity mapping illuminates otherwise intractable locale-specific metabolite signaling and stress response programs influencing organ-specific decision-making.

**IN BRIEF:** Context-specific protein reactivity is a cornerstone of biological stress responsivity, and thus has important ramifications for drug discovery; no current method can inform on these parameters. Here we debut a method that cartographs locale-specific actionability to reactive metabolites; Cyp-33e1, a gut-specific responder, emerged to partition between electrophile-driven localized enzymatic turnover that triggers gut-specific metabolite production shaping global lipid storage, and localized electrophile sensing that marshals global stress response.

**HIGHLIGHTS:**

- Localis-REX maps organ-specific electrophile-responsive proteins in worms
- Hits are neither identified by organ-specific Ultra-ID nor biased by localized expression
- Hits are enriched in proteins functionally relevant to stress-related phenotypes
- Localized responsivity of gut-specific hit, cyp-33e1, shapes global lipid availability

---

Single cell RNA-sequencing and proximity mapping technologies have revolutionized precision investigations into locale-specific biological makeups. At the proteome level, pioneering techniques like BioID/TurboID, APEX, and µMap are leading the way to census protein residents at defined biological locales^1–3^, although techniques like BONCAT that profile localized nascent proteomes^4,5^ are also insightful. Nonetheless, we remain limited in mapping locale-specific chemical actionability, i.e., how protein responsivity to small molecules varies as a function of tissue or subcellular contexts. Indeed, mapping the plasticity of chemical reactivity as a function of locale remains outside the remit of all cutting-edge methods. This is because noninvasive strategies to selectively dispatch biologically or drug relevant, yet ephemeral, small molecules to a specific organ/tissue within an intact organism on demand, are scarce. Indeed, to identify chemically actionable protein targets, reactive small-molecule metabolites, or drug-like electrophilic fragments, are often administered in bulk to cells/animals. Such approaches have limited spatiotemporal precision, and poor control of reactive ligand distribution, bioavailability and metabolism. These uncontrolled approaches are also prone to issues associated with reactive electrophile overload, including toxicity and stress.

Unfortunately, signaling modalities of electrophiles are particularly context dependent, with productive signaling occurring upon the correct confluence of situation, association, and redox state/microenvironment of specific proteins^6^. Uncovering such context-specific responsivity to reactive metabolites is biomedically important, because innate electrophile signaling can modulate specific signaling pathways^7–11^. Such data can uncover rapid response circuits, and also elucidate (specific) functions of specific proteins within these pathways, e.g., in specific subcellular compartments^12^ or cell types^13^. As electrophile–protein engagement mimics how covalent drug-protein interactions occur, such discoveries are also relevant to drug design^14,15^.

Here we evaluate the hypothesis that by releasing a transient burst of a specific native electrophile in a specific organ of a live, intact worm, we can fingerprint localized native proteins with privileged functional electrophile responsivity in the organ of interest. We achieved this feat using a method that mimics natural, localized, transient signaling by the native electrophilic metabolite, 4-hydroxynonenal (HNE)^6^. This setup, termed ‘organ specific (OS) Localis-REX’ hereafter, allows the best protein sensors to be HNEylated, prior to interception by bystander proteins (which are frequently labeled during bolus administration). We used enrichment-coupled quantitative mass spectrometry to identify electrophile actionable proteins in specific locales. The resulting maps of local proteome responsivity within distinct organs revealed that >70% of the targets displayed localized electrophile responsivity, unreflective of tissue-specific abundance^16^. Proteins identified in OS-Localis-REX workflow were also not identified by tissue-specific biotinylation using Ultra-ID, a hyperactive variant of BioID^17^. Several of the identified proteins emerged to be involved in tissue-specific homeostatic regulatory events consistent with the locales in which they were identified. More frequently than expected based on literature precedents, knockdown of these genes changed responsivity to electrophilic stress, consistent with OS-Localis-REX hits being involved in electrophilic stress management. We pursued one specific candidate, cyp-33e1; both *C. elegans* and human orthologs emerged to be electrophile responsive. In *C. elegans*, despite similar expression across all 3 organs, cyp-33e1 was an active electrophile responder specifically in the gut, consistent with our data from OS-Localis-REX. This localized electrophile responsivity in turn triggered a gut-specific signaling program crucial for global lipid homeostasis.

These data support that contrary to other methods^18^, OS-Localis-REX directly informs on chemical actionability within specified local proteomes in live animals, independent of locale-specific expression levels. Beyond paving the way toward an arsenal of chemical genetic methods to perturb *local* proteome function in living systems, data so derived *proffer* a means to identify organ-specific responders to localized stress. Such information is not easily accessible by traditional profiling technologies.

## RESULTS

### Organ-specific controlled electrophile generation, OS-Localis-REX, in *C. elegans*

Our experimental design involved repurposing our precision electrophile delivery tools^7^ to achieve light-driven release of a reactive native electrophile within specific organs in *C. elegans*. To anchor our small-molecule photocaged-electrophile probe^19,20^ that interacts specifically with Halo protein in specific tissues in live worms, we generated worms with tissue-specific expression of GFP-Halo. Our chosen promoters to drive GFP-Halo expression were: myo-2p (pharyngeal expression); ges-1p (intestinal expression); and myo-3p [body-wall (BW) muscle expression]. We surmised that following exposure of these live transgenic worms to our probe, Halo-protein would irreversibly anchor our photocaged-electrophile probe, such that following washout of unbound probe, light illumination would drive tissue-specific electrophile release. Worms with non-integrated extrachromosomal arrays were thus created, which were transmissive at ∼70% frequency, and showed expected tissue-specific, albeit, as expected mosaic, GFP expression (**Figure 1A**). There was less than 2-fold difference in Halo expression across these constructs (**Figure S1A**).

**Figure 1.**
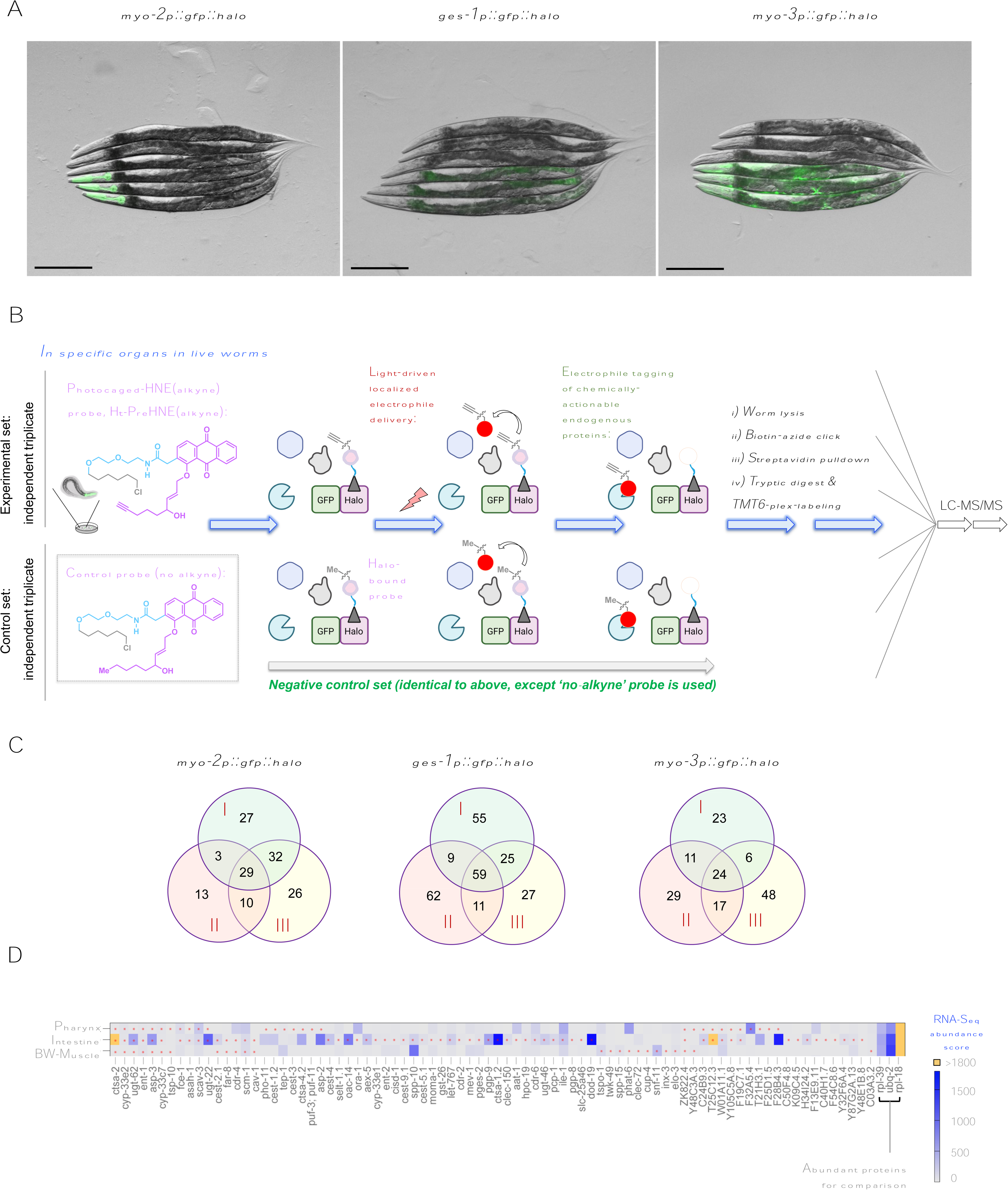
OS-Localis-REX identifies *chemically actionable* tissue-specific targets independent of native tissue preference/abundance. **(A)** Fluorescence imaging confirmed tissue-specific GFP-Halo expression in Day 1 adult worms: myo-2, ges-1, and myo-3 promoters respectively drive GFP-Halo expression in pharynx, intestine, and body-wall muscles. [3 transgenic worms (lower) against 3 wild-type N2 worms (upper)]. Scale bars: 200 µm. See also **Figure S1A-S1D**. ***Note:*** *due to mosaicity, GFP expression patterns were not identical across worms in the same group*. **(B)** OS-Localis-REX: tissue-specific Halo transgenic worms were treated with either photocaged HNE(alkyne) [Ht-PreHNE(alkyne)] (experimental set comprising independent biological triplicates, top) or a non-alkyne-functionalized variant, Ht-PreHNE(no alkyne) (triplicate controls, bottom). Following washout, light (5 mW/cm^2^, 366 nm, 3 min) exposure rapidly (*t*_1/2_ ∼ 0.5 min) liberates the electrophile, HNE(alkyne) or HNE(no alkyne). Alkyne-functionalized HNEylated native protein responders were enriched by biotin-azide click coupling and streptavidin pulldown, whereas HNE(no-alkyne)-tagged proteins were not enriched. Samples were subjected to trypsin digest, isotopic labeling, and LC-MS/MS analysis. See also **Figure S1E-S1H**. **(C)** 29, 59, and 24 proteins were selected as significant electrophile-responders in indicated tissue-specific GFP-Halo transgenic strains, across three independent biological replicates (labeled as I, II, III) against corresponding controls. See also **Figure S1I**; **Data S1-S3**. **(D)** 59 from 82 (∼72%) non-overlapping significant hits across the 3 strains reflect electrophile-actionable protein targets, whose tissue-specific electrophile responsivity manifested in tissues (marked by *) that are not their canonical native/preferred locale(s) (represented by heatmap indicating transcript scores derived from cell-specific RNA-Seq datasets^16^; yellow denotes heat-map scores >1800). ***Note*:** Highly-abundant *C. elegans* proteins—rpl-39, ubq-2, and rpl-18, and their tissue-specific distributions, are included for comparison: these proteins were not scored as electrophile-responders by OS-Localis-REX. See also **Data S2**; **Figure S1J**.

We chose to release the endogenous lipid-derived electrophile, HNE. This choice was spurred by HNE’s prevailing, yet poorly-understood, context-specific pathophysiological roles, including in myriad neurodegenerative disorders^6,21,22^, where *C. elegans* is an established model^23–25^. Furthermore, recent work indicates that HNE houses a natural electrophilic pharmacophore of relevance to precision medicine design^11,14^. We thus used our photocaged HNE(alkyne-functionalized) probe (Ht-PreHNE, hereafter, **Figure 1B** and **S1B**) that undergoes efficient cellular uptake; is stable when bound to Halo; and non-toxic in multiple live models^19,20,26^. Alkyne modification allows efficient HNE detection, orthogonal from a non-alkyne-tagged variant used as a control during OS-Localis-REX profiling. Alkyne-modified HNE is commonly deployed, and is widely accepted to be non-invasive as the alkyne is remote from the site of electrophilic chemistry and innocuously changes HNE’s physiochemical properties.

Using an optimized Ht-PreHNE dosage over 6-h treatment followed by washout of excess unbound probe, the probe selectively labeled the desired tissues in all localized Halo expressing strains (**Figure S1C** and **S1D**). Photouncaging of HNE(alkyne) using a low-power hand-held lamp (5 mW/cm^2^, 366 nm), was rapid, *t*_1/2(photouncaging)_∼0.5 min (**Figure S1E** and **S1F**). The liberated HNE labeled proteins in live worms (**Figure S1G**).

### Mapping organ-specific functional first responders in live *C. elegans* using OS-Localis-REX

We proceeded to identify HNE responders as a function of locale. Tandem mass tag (TMT)-6-plex labeling^27^ was used to quantitatively score enriched targets from each strain, over three independent biological replicates, against a control triplicate set (processed identically, but treated with a non-alkyne-functionalized probe) (**Figure 1B** and **S1G-I**; **Data S1** and **S2**; **Method S1**). For each strain, enriched proteins were strongly correlated (Pearson r>0.95) (**Data S3A**). We thus identified responders in the pharynx (29), intestine (59), and BW-muscles (24) (**Figure 1C**). ∼70% of these hits were HNE responsive only in one tissue, ∼20% were responsive in two tissues (**Data S2**). Several tissue-specific responders are involved in processes relevant to the tissue in which they sensed electrophiles: e.g., aex-5, an intestinal-specific responder, functions in the intestinal regulation of *C. elegans* defecation motor program (DMP)^28^; aex-5 mutant animals have impaired intestinal fat metabolism^29^.

### Tissue-specific electrophile responsivity identified by OS-Localis-REX is not correlated with tissue-specific expression

Comparison of our Localis-REX hits against published *C. elegans* tissue-specific single-cell RNA-sequence (scRNA-Seq) datasets^16^, showed that tissue-specific electrophile responsivity was independent of expression in respective tissues (**Figure 1D**). A similar conclusion was drawn upon comparison of our Localis-REX hits with PAXdb, a protein abundance database derived from published experimental data (**Figure S1J**). Such observations indicate that locale-specific expression does not strongly influence locale-specific electrophile responsivity, consistent with a recent report from our laboratory investigating subcellular responsivity to electrophiles^12^.

We sought further evidence to rule out that our identified tissue-specific HNE responders were biased by protein abundance in the tissues we profiled. We engineered worms expressing Ultra-ID (the smallest and most active promiscuous biotin ligase available to date^17^ for proximity/interactome mapping). Tissue-specific Ultra-ID expression was driven by the same promoters used in OS-Localis-REX. We validated Ultra-ID localization in each strain, by imaging *mCherry* [encoded upstream of *Ultra-ID*, spaced by P2A ‘self-cleaving’ peptide^20^ (**Method S1**; **Figure 2A** and **S2A**)]. In these organ-specific (OS) Ultra-ID strains, total proteome abundance/composition^30^ was largely unaffected (**Data S4**; **Figure S2B**). Streptavidin blot confirmed biotinylation activity of Ultra-ID in each strain (**Figure S2C**).

**Figure 2.**
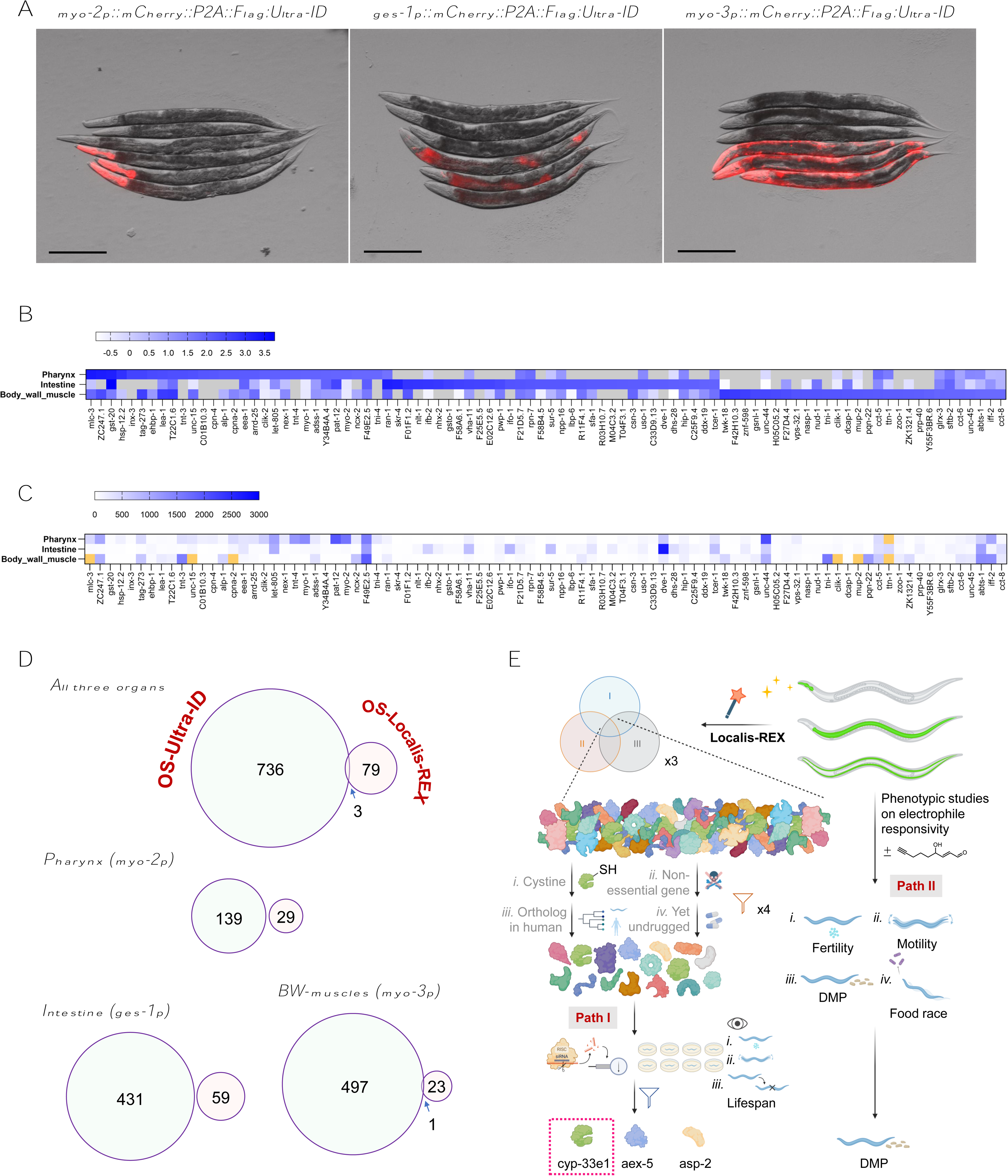
OS-Localis-REX and OS-Ultra-ID exhibit divergent target spectra and functional enrichment. (**A**) Fluorescence imaging of mCherry expression in day 1 Ultra-ID transgenic hermaphrodites. Each image shows 3 tissue-specific mCherry-P2A-UltraID-expressing animals (lower), against 3 wild-type N2 worms (upper, controls). Scale bars: 200 µm. ***Note:*** *due to mosaicity, mCherry expression patterns are not identical across all 3 transgenic worms.* See also **Figure S2A-S2D**. (**B**) Tissue-specific protein abundance mapped by OS-Ultra-ID, featuring top ∼30 hits (from 90 total) ranked based on log_2_FC (fold change of designated transgenic strain against wild type) and FDR for all hits ≤0.05. Gray boxes denote corresponding proteins not identified. See also **Figure S2E-S2G**, **Data S5**. Scale bar above designates Log_2_FC (designated transgenic strain vs. wild type) derived from TMT ratios (see **Data S5a-c**). (**C**) OS-Ultra-ID hits shown in **Figure 2B** were searched for their corresponding tissue-specific abundance scores from cell-specific RNA-Seq datasets, covering the same tissues. Yellow boxes correspond to transcript scores >3000 (thus, out of range of the indicated scale bar). (**D**) *<u>Top:</u>* By applying FDR≤0.05 and Log_2_FC (designated transgenic strain vs. wild type) > 0 to OS-Ultra-ID hits across all 3 tissues, the resulting 739 targets (**Data S5a-c**) featured only 3 proteins that overlapped with chemically-actionable responders mapped by OS-Localis-REX (**Figure 1D**, **Data S1-S3**). *<u>Bottom:</u>* Same analysis as above but within each tissue, revealing no overlap between OS-Ultra-ID (larger circle) and OS-Localis-REX (smaller circle) hits, except in body-wall muscles (one overlapping target, cav-1). Such divergence extends to outputs obtained from functional pathway enrichments. See also **Figure S3**, **Data S1** vs. **S5**. (**E**) Roadmap illustrating OS-Localis-REX hit discovery and subsequent functional and phenotypic validations. Top left: selection criteria (i-iv) applied to 82 first-responders from OS-Localis-REX (see **Figure 1C** and **1D**) manifesting at least one cysteine; human orthologs; and disease relevance. Essential genes (not amenable to KO/KD-studies in whole animals) were further removed; hits yet unknown to be druggable/ligandable were favored, while ensuring final selected list reflected both unique and common hits across three different tissues. The resulting 15 candidates were then subjected to: (**Path I**) RNAi phenotypic screen, against commonly-assayed *C. elegans* phenotypes: (i) egg-laying (fertility); (ii) movements (motility); and (iii) lifespan (survival). ***Note***: tissue-specific RNAi screen, in lieu of/supplementing ubiquitous knockdown, may be undertaken here, as was done for aex-5. A parallel RNAi screen in the presence or absence of acute electrophilic stress, focusing on egg-laying, pinpointed 3 potential electrophile-stress-dependent functionally-relevant candidates: cyp-33e1, aex-5, and asp-2. T-REX cell-based validations singled out cyp-33e1 (and its human ortholog, CYP2A6) as kinetically-privileged responder, and established labeling sites. In **Path II**, DMP, a phenotype reporting intestinal physiology, was first identified as the phenotype most affected by acute electrophilic stress.

To reduce background biotinylation, we used biotin-auxotrophic *E. coli* as food^31^ (**Figure S2C** and **S2D**). We then executed OS-Ultra-ID, mapping locale-specific protein abundance over 3 independent biological replicates, against 2 separate control groups in triplicate (**Data S5**). Briefly, post worm lysis, streptavidin enrichment, on-bead digestion of enriched proteins, and labeling of eluted peptides with isotopomeric TMT labels^27^, peptides were compared by MS (**Figure S2E** and **S2F**). This workflow is similar to the enrichment workflow deployed in OS-Localis-REX, except endogenously biotinylated proteins cannot be removed prior to enrichment, as they are in OS-Localis-REX (**Figure 1B**, **S1I**, **S2E** and **S2F**; **Data S3B**).

Assessment of OS-Ultra-ID datasets unveiled several key points (**Figure 2B-D** and **S2G**). (***i***) Numerous targets were captured exclusively in a single tissue: focusing on enriched hits (with protein FDR≤0.05) bearing 2-fold change against biotin-treated wild-type controls, 32, 107, and 125 (**Figure S2G**: lower-right panel) respectively constitute proteins abundantly expressed in one locale. (***ii***) Taking 30 top-ranked hits (FDR≤0.05) from each tissue as representatives, the resultant tissue-specific expression profiles (**Figure 2B**) were consistent with those derived from scRNA-Seq^16^ (**Figure 2C**), validating the scRNA-seq datasets used above (**Figure 1D**). (***iii***) Despite the fact that both OS-Localis-REX and OS-Ultra-ID deployed similar biotin-based enrichment strategies (**Figure 1B** and **S1I**; Cf. **Figure S2E** and **S2F**), there was minimal overlap between the techniques (**Figure 2D**). Even when all 739 enriched proteins (FDR≤0.05) across all organs from OS-Ultra-ID were considered, only 3 proteins (**Figure 2D**, top row) overlapped between OS-Ultra-ID and OS-Localis-REX: *two of these overlapping hits were identified in different locales*. Based on the parity of scRNA-Seq and OS-Ultra-ID, as well as OS-Ultra-ID and OS-Localis-REX identifying some overlapping proteins but mostly in *different* locales, we conclude that OS-Localis-REX pinpoints bona fide responders whose HNE responsivity is augmented in specific locales, independent of expression. Finally, functional enrichment analyses using two independent programs (Cytoscape and g:*Profiler*)^32^ showed non-overlapping GO subterms and KEGG pathway terms between OS-Localis-REX vs. OS-Ultra-ID in each tissue (**Figure S3**).

### Proteins mapped by OS-Localis-REX influence animal health and physiology

Among the 82 responders mapped by OS-Localis-REX across all strains, 18 proteins are not yet functionally annotated in Wormbase. BLAST analysis showed that 6 of these 18 proteins showed ∼17– 47% sequence identity with human proteins (**Data S2**). Thus, ∼76%, 62 *C. elegans* responders out of 82, represented proteins with human orthologs. ∼67% (55/82) are disease-associated genes, but only 43% (35/82) are chemically actionable based on representation in DrugBank (**Data S2**). With the goal of identifying interesting “undruggable” proteins for in-depth investigations, we studied functional relevance of 15 selected hits having: (**i**) human orthologs of disease relevance (**Figure 2E** and **S4A**); (**ii**) no known electrophile sensitivity/druggability (**Data S2**); and (**iii**) both human and worm orthologs housing at least one cysteine (**Figure 2E**). These 15 candidates further reflect both unique and common hits across the three tissues (**Figure S4A**; **Data S2**).

Initially, we assessed the functional relevance of these proteins using RNAi (**Figure 2E**), focusing on readily measurable phenotypes: (**i**) survival, (**ii**) motility, and (**iii**) reproductive fitness. Animals were fed dsRNA-producing HT115 bacteria containing plasmids from the Ahringer library^33^. Knockdown efficiencies (**Figure S4B; Method S1**) were >90% for most genes. However, efficiencies were closer to 50% for 4 genes (**Figure S4B**), possibly reflecting low efficiency, or expression localization to regions impermeable to RNAi, such as neurons^34^, which are not the focus of our study. RNAi targeting let-767 (a positive control) caused development and growth defects, as previously reported^35–37^.

***(i) Survival management:*** deficiency in aex-5, pho-11, and cyp-33e1 prolonged mean lifespan, whereas puf-3 and ent-1 (beyond let-767, positive control) deficiency reduced lifespan (**Figure 3A**, **S4C** and **S4D**). Our data on aex-5 are consistent with a previous report^38^. pho-11-knockdown reduces survival of otherwise lifespan-extended *daf-2* and *daf-16* mutant animals^39^, although pho-11-deficiency in the wild type background has not previously been assayed. Puf-3 RNAi-fed animals manifested shortened lifespans. In mouse, hyperactivity of a closely-related isoform of puf-3’s human ortholog, PUM2, drives premature aging^40^. Notably, ent-1 RNAi, in our hands, reduced the lifespan, but a previous study reported no effect on lifespan^41^. Nonetheless, overall consistency of the outcomes from our survival-curve analysis, and additional interesting new findings with relatable phenotypes in human orthologs, altogether hint conserved importance of OS-Localis-REX targets in animal survival.
***(ii) Motility:*** Pharynx and BW-muscles are critical for nematode movement. Body bends per minute (BPM) is an indicator of *C. elegans* musculature function. L3/L4 larvae subjected to let-767 RNAi manifested uncoordinated movement as reported^35,37^, validating our workflow. Knockdown of several other hits showed significant motor defects (**Figure 3B**, **S4E**). Applying a stiff cutoff (± 2σ) to Gaussian fit of the resultant measurements across all hits over the three developmental stages assayed revealed that, in addition to let-767 RNAi, asp-2 RNAi caused significant positive deviation from the mean BPM (**Figure S4F**). Asp-2 is orthologous to cathepsin-E, a disease-associated aspartic protease expressed in mammalian immune system and epidermis. Cathepsin-E deficiency decreases motility of dendritic cells^42^ and Schwann cells^43^.
***(iii) Reproductive fitness:*** The effects of RNAi on the total number of fertilized eggs were next assessed for the 15 candidate proteins. let-767 RNAi decreased fertility, as expected (**Figure 3C**). The same result was observed in animals treated with RNAi of mev-1, whose human ortholog corresponds to subunit-C of the mitochondrial tumor suppressor protein, succinate dehydrogenase (SDH, also known as Complex-II), a non-essential component of the electron transport chain and contributor to the TCA cycle^44^. Loss-of-function mutations in SDH subunits increase predisposition to cancers^45^. In *C. elegans*, a homozygous missense mutant of mev-1 with depleted complex-II activity^46^ shortens lifespan^47–49^, although RNAi of mev-1 has no effect on lifespan^50,51^, consistent with our data (**Figure 3A**, **S4C** and **S4D**). mev-1 loss-of-function mutant worms have diminished fecundity^50^, in agreement with our data (**Figure 3C**). ent-1 RNAi-fed animals, beyond reduced survival (**Figure 3A**, **S4C** and **S4D**), displayed diminished fecundity (**Figure 3C**). The human ortholog of ent-1 is SLC29A1, encoding equilibrative nucleoside transporter 1 (ENT1). Polymorphisms in SLC29A1 are linked to multiple diseases and drug resistance. Ent1^-/-^ mice show paralysis among other defects^52,53^. By contrast, depletion of ctsa-4.2 (human ortholog: CTSA, lysosomal Cathepsin-A serine carboxypeptidase) and cup-4 gene, encoding a coelomocyte-specific ligand-gated ion channel (human ortholog: CHRNA9), have increased fertility (**Figure 3C**).

**Figure 3.**
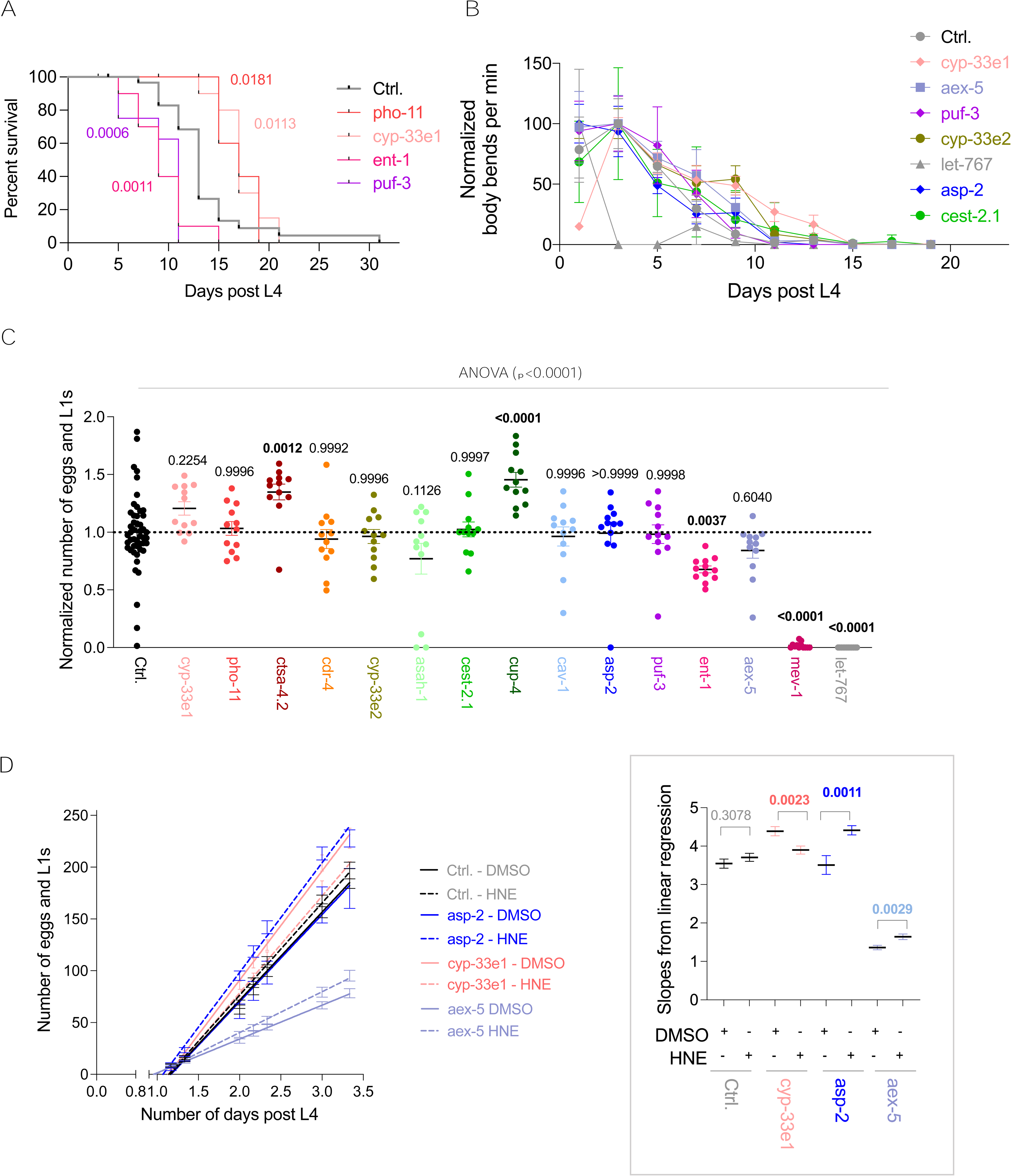
RNAi knockdown of OS-Localis-REX hits influences survival, motility, and reproductive fitness. **(A)** Pho-11, cyp-33e1, ent-1, and puf-3 (see **Figure S4A-S4C**, **Method S1**) were identified as 4 genes (from 15) whose RNAi significantly affected survival, following application of log-rank (Mantel-Cox) test, with indicated P values. Also see **Figure S4D**: Aex-5 was significantly different from control when Gehan-Breslow-Wilcoxon (GBW) test was applied. **(B)** Normalized body bends per min of worms where RNAi of individual indicated genes significantly altered motility with respect to control animals, as judged by Gaussian distribution analysis (see **Figure S4E** and **S4F**). Data were normalized such that the highest body-bending rate corresponds to 100. **(C)** Total number of eggs and L1s from indicated RNAi-fed worms in the absence (Ctrl.) and presence of RNAi of indicated genes. Data were normalized to the mean of control (Ctrl.). P-values were calculated using Dunnett’s multiple comparisons test against control group. Corrected p values < 0.05 are in bold. **(D)** Linear regression analysis of the number of fertile eggs over indicated days where knockdown of indicated genes resulted in differential reproductive fitness, in the presence (HNE) vs. absence (DMSO) of electrophilic stress, compared to knockdown control (Ctrl.). See also **Figure S4G** and **S4H** for data from all 15 genes. ***Inset:*** slope values from linear regression. P values from unpaired two-tailed Students’ t-test. Bonferroni adjustment was applied, p: adjusted, 0.003125. All data present mean±SEM. Sample sizes and decision tree for statistical treatment in **Method S3**.

### OS-Localis-REX hit proteins are implicated in stress-induced changes in animal reproductive fitness

We next investigated how deficiency of our candidate proteins impacts response to HNE stress. For these experiments, we measured reproductive fitness, as this phenotype allows more high-throughput scoring. Furthermore, acute electrophilic stress exerts strong effects on cell-cycle in nematodes^54^; and reproductive fitness is strongly dependent on cell-cycle progression. We thus quantitatively compared fecundity and fertility scores in DMSO- or HNE-treated worms for each RNAi. Since the number of fertile eggs laid proceeds at a consistent rate over several days during young adulthood, the cumulative total of eggs laid can be reliably fit by linear regression. Using this analysis, we assessed to what extent the gradients for DMSO- and HNE-treated worms were different for each condition. There was no significant difference between the twain for control-RNAi treated worms (p, unadjusted, value 0.3078) (**Figure 3D**). Given that we have 15 different RNAi’s and a control, we applied the Bonferroni adjustment to the resulting analysis (null hypothesis rejected if p < 0.003125), which revealed that cyp-33e1, aex-5, and asp-2 play functional roles in reproductive fitness during stress (**Figure 3D**, S4G and S4H). The fact that 20% of our studied proteins were linked to martialing HNE stress is surprising. Although the percentage of HNE responders across proteomes is unknown, several methods have shown that proteins highly sensitive to HNE are scarce: a previous study estimated that ∼98% of cysteines are unresponsive to specific electrophiles^55^; another study, using electrophiles more reactive than HNE, estimated that ∼5% of cysteines are reactive in worms^56^.

In parallel, we also explored preliminarily how phenotypic changes may be functionally linked to specific tissues where these proteins were identified by OS-Localis-REX, harnessing tissue-specific gene knockdown in *C. elegans*. Using aex-5 as an initial test case, the diminished egg laying rate was recapitulated when aex-5 RNAi was targeted to intestinal cells, but not to BW-muscle cells (**Figure S4L**). Interestingly, aex-5 was mapped by OS-Localis-REX as an intestine-specific responder (**Figure 1D** and S4A; Data S1 and S2). Based on a previous report indicating a link between global deficiency of aex-5 and decreased fat accumulation^29^, we further examined the effects of aex-5 deficiency on global lipid content. Using the oil red O (ORO) staining assay, we found aberrant lipid accumulation in the anterior intestine in L4 aex-5-depleted worms (**Figure S4I**) but decreased fat in adult aex-5-depleted worms (**Figure S4J**). Notably, intestine-specific aex-5 knockdown mimicked these effects, but these features were absent in the knockdown control or BW-muscle-specific RNAi animals, in both young and adult hermaphrodites (**Figure S4K**).

### Cyp-33e1, and its human ortholog CYP2A6, are kinetically-privileged first responders to HNE; however HNE only partially inhibits oxidoreductase activity

Based on the outcomes above, we further investigated the three candidate proteins, cyp-33e1, aex-5, and asp-2, specifically in the context of localized electrophile sensing capabilities. First, we used cell-based T-REX quantitative electrophile responsivity ranking assay^19^, to evaluate electrophile sensing propensity of each protein in live cells. T-REX liberates HNE in the vicinity of a protein of interest (POI) and hence can quantitatively rank^7^ HNE responsivities in live cells, by assessing the fraction of HNE that covalently labels a specific POI, prior to HNE irreversibly diffusing away from the POI (**Figure S5A**). T-REX is thus a particularly stringent and quantitative test of electrophile sensitivity. From these experiments, cyp-33e1 emerged to be the most HNE responsive of the three POIs (**Figure 4A**, S5B and S5C). The human ortholog of cyp-33e1, CYP2A6, was also HNE responsive (**Figure S5D**); PCSK1, the human ortholog of aex-5, was not (**Figure S5E**).

**Figure 4.**
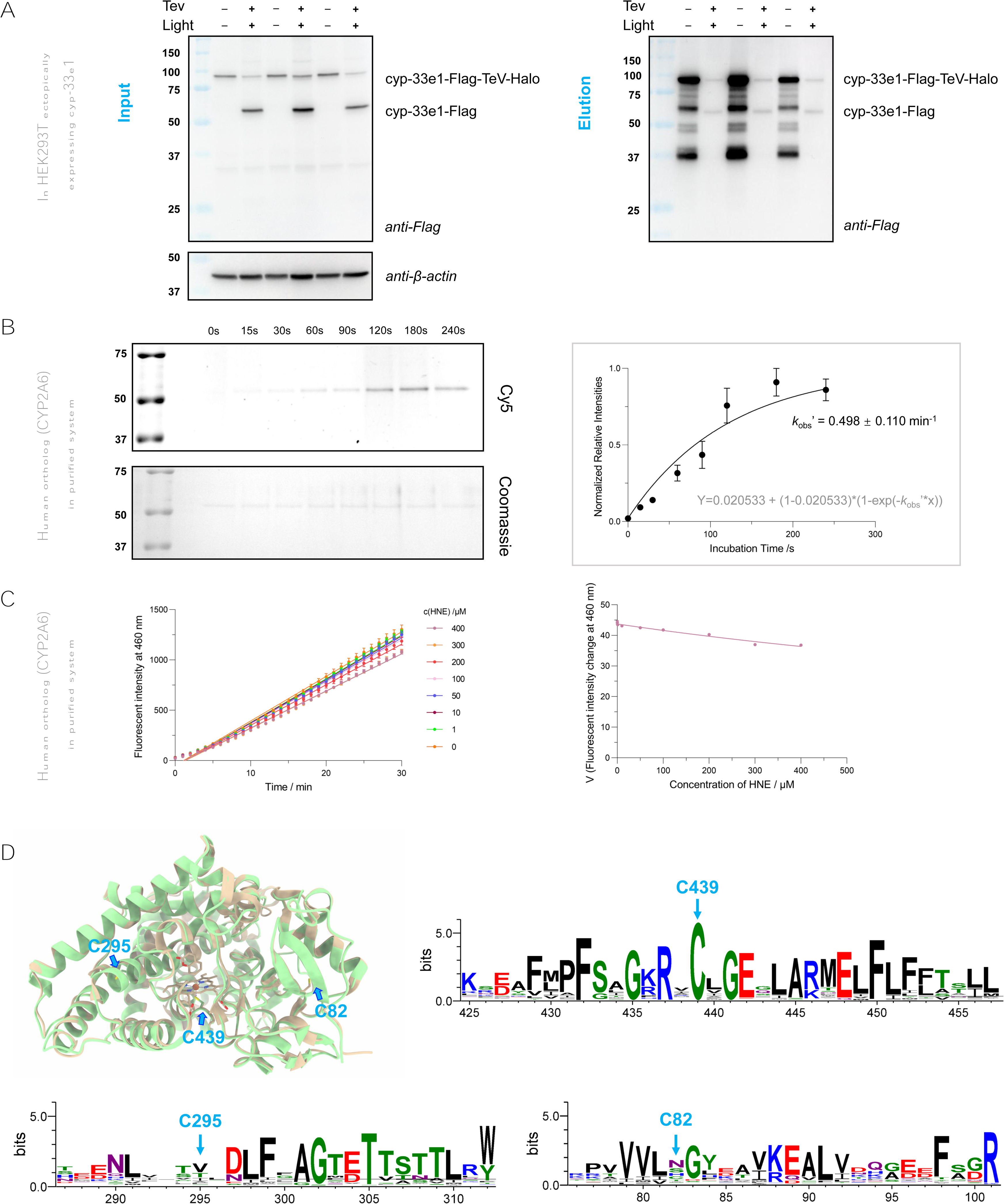
cyp-33e1, and its human ortholog CYP2A6, are kinetically-privileged responders. (**A**) cyp-33e1 was scored as a first responder using T-REX in HEK293T (see **Figure S5A**). Input and elution samples resulting from Click-biotin pulldown of the protein post cell lysis, and subsequent TeV-protease-mediated separation of Halo and cyp-33e1, analyzed by western blot using anti-Flag and anti-β-actin antibodies. MWs: cyp-33e1-Flag-TeV-Halo-His: 95 kDa, cyp-33e1-Flag (post TeV cleavage): 60 kDa. ***Note*:** Incomplete TeV cleavage results in residual full-length fusion protein in ‘+ TeV’ lanes in both Input and Elution. Inadvertent cleavage occurs during pulldown, leading to multiple Flag-positive bands in negative-control samples in Elution. Nonetheless, the HNE(alkyne)-tagged enriched species in experimental samples (+ TeV, + light) in Elution corresponds to full-length cyp-33e1. (**B**) Labeling of recombinant CYP2A6 labeling by HNE(alkyne). ***Inset***: quantification. *k*_obs_’ = 0.50 ± 0.110 min^-1^ was calculated by normalized HNEylation, to CYP2A6 amount deployed, and fitting to: Y=0.021 + (1-0.021) * (1-exp(-*k*_obs_’*x)). For comparison with the previously-reported HNE responders (**Table S9**), the *k*_obs_’ was adjusted to equivalent concentrations as in previous studies, resulting in *k*_obs_ ∼ 4 × 10^8^ M^-2^s^-1^. (**C**) Recombinant CYP2A6 was partially inactivated by HNE. Data (left) were fit by linear regression to derive reaction velocities which were subsequently plotted against [HNE] (right), and fit to Morrison tight-binding equation: v = *V*_o_*(1-(((([E]+(*K*_i_*(1+([S]/*K*_M_))))-((([E]+[I]+(K_i_*(1+([S]/K_M_))))^2)-4*[E]*[I])^0.5))/(2*[E]))). [I]: HNE concentration; [E]: CYP2A6 concentration; [S]: 3-cyano coumarin concentration; for *K*_M_, see **Figure S5F**. The Morrison *K*_i_ : 207.9 ± 35.4 μM. (**D**) Overlay of human CYP2A6 (**gold**) (PDB:1Z10) and *C. elegans* cyp-33e1 (**green**) (homology model) in ribbon presentation, and sequence logos for sequences flanking indicated cysteines, derived from 936 sequences (isoforms) integrating 38 metazoan species (WebLogo). Sequence alignment for 936 sequences was performed using Super5 algorithm in MUSCLE5^73^. C439, catalytically-essential residue, is highly conserved. Almost all Cyps house at least one additional cysteine beyond C439. ***Note***: C14 of CYP2A6 lies within the N-terminal signal peptide sequence and the construct used for crystallization does not bear this sequence. Also see **Figure S5G** and **S5H**. All data present mean±SEM. Sample sizes and decision tree for statistical treatment in **Method S3**.

As cyp-33e1 emerged to be the most avid HNE sensor, and HNE sensing was conserved to its human ortholog, we focused on cyp-33e1. We subjected purified recombinant CYP2A6 to HNE, and detected time-dependent labeling that saturated after ∼150 s (**Figure 4B**). Thus CYP2A6 reacts with HNE faster than several HNE sensors reported from other groups that we have studied in vitro previously^6,57^. However, the time to saturation was slower than some of the electrophile sensors that we have reported (e.g., Ube2V2^58^), although the conditions were admittedly different, rendering direct comparisons difficult.

Several studies implicate either inhibitory, simulatory, or no effects induced by HNE on various mammalian CYP-P450 isozymes^59–63^: CYP2A6 has not previously been examined. Treatment of commercially available CYP2A6 with HNE demonstrated a minor impact on enzymatic activity that occurred at high HNE concentrations, *K*_i_ ∼ >200 µM (**Figure 4C** and **S5F**). Given that an excess of HNE had a minor effect on CYP2A6 activity, and that relatively low HNE concentrations *labeled* the enzyme, we proposed that there are at least two HNE-labeling events within CYP2A6: one occurs rapidly with little impact on activity (such an event would be detected by our profiling strategy, even though under these first-responder conditions, CYP2A6 would remain active); the second event reduced activity. We have reported such scenarios previously^12,13,57^. However, we cannot exclude, at this juncture, the possibility that the second labeling event occurs due to isomerization/aggregation of the first labeled species.

### Cyp-33e1, and its human ortholog CYP2A6, undergo electrophile sensing at two cysteines

Cysteine is the most kinetically-privileged, among nucleophilic protein residues, in terms of reactivity against Michael-acceptor-based electrophiles, including HNE^6^. Cyp-33e1 harbors 2 cysteines, C439, a conserved Heme-iron-binding cysteine within the catalytic site (**Figure 4D, S5G and S5H**), and C295, an off-active-site residue that based on homology modeling, resides in a loop bridging an α-helix and a β-sheet (**Figure 4D**). The human ortholog CYP2A6 houses 3 cysteines: C14 and C82, in addition to the conserved catalytic-site C439. Overlays of the available X-ray crystal structure of human CYP2A6 (PDB: 1Z10) and a homology model of cyp-33e1 indicated that C82 and C295 in human and worm are located in different regions of the protein; C14 is not present in the worm protein (**Figure 4D and S5G**).

To delve deeper into underlying mechanisms and potential site specificity in sensing and inhibition, we created single, double, and where applicable, triple cysteine mutants of both human and nematode constructs and replicated HEK293T cell-based T-REX assays^7,19^. All mutants were expressed, although with variable expression levels (**Figure S6A** and **S6B**). Expression-level differences were subsequently taken into account in calculating ligand occupancy^7^ for each POI, derived from T-REX (**Figure 5A, 5B, S6A** and **S6B**).

**Figure 5.**
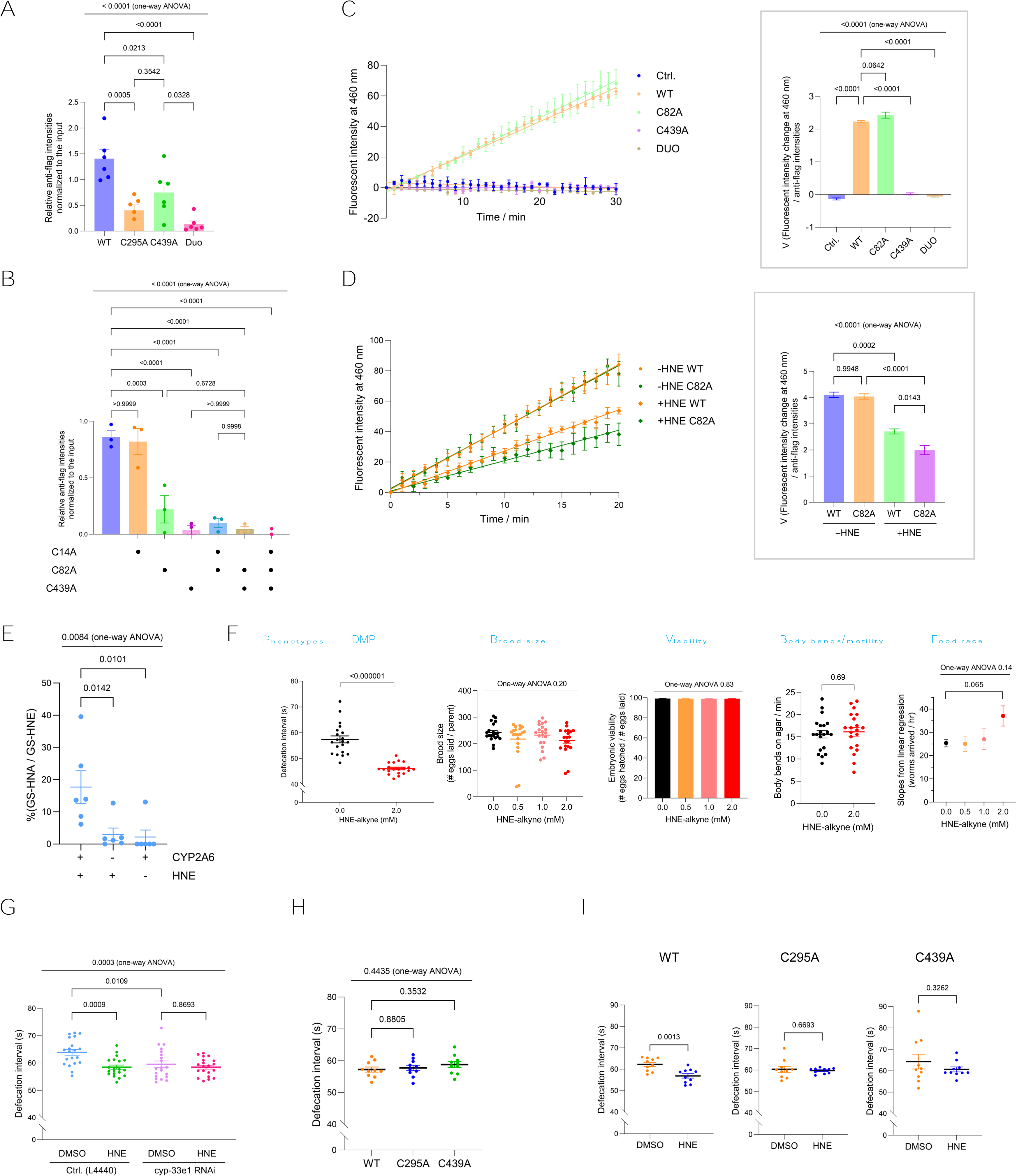
Cyp-33e1/CYP2A6, sense HNE at a conserved catalytic cysteine and an off-active-site cysteine; the former inhibits activity, yet labeling at both sites regulates stress-dependent changes in intestinal physiology. **(A)** Electrophile-sensing ability of cyp-33e1 WT; single; or double (‘duo’) cysteine-to-alanine mutants, was quantitatively compared following T-REX in HEK293T (see **Figure S5A**) and click-biotin pulldown of HNEylated protein, as in Figure 4A. Plot shows quantification of **Figure S6A**. **(B)** Identical to Figure 5A except electrophile sensing ability of CYP2A6 WT; single; double; or triple cysteine-to-alanine mutants, was quantitatively compared. ***Note***: quantification takes into account of differences in WT/mutant expression. See also **Figure S6b; Data S3C** and **S3D**. **(C)** Oxidoreductase activities measured for in-house generated CYP2A6 (WT or indicated single and double cys-to-ala mutants) were fit by linear regression (Ctrl.: y= -0.13*x+ 3.26; WT: y= 2.23*x-1.53; C82A: y= 2.25*x-2.81; C439A: y= 0.021*x-0.284; DUO: y= -0.0578*x-0.777), to derive reaction velocities. Background subtraction and normalization were applied to all datasets. *<u>Inset on right</u>:* quantification of relative slopes. Negative control denotes cells without transfection. See also **Figure S6C** and **S6D**. **(D)** Progress curve analyses of CYP2A6 (WT or C82A) enzymatic turnover, in the presence and absence of HNE (400 μM). Data were fit by linear regression (-HNE WT: y= 4.11*x+ 2.1; -HNE C82A: y= 4.05*x+ 2.6; +HNE WT: y= 2.71*x+ 0.16; +HNE C82A: y= 2.00*x+ 0.95)x. Background subtraction and normalization were applied to all datasets. *<u>Inset on right</u>:* quantification of the relative slopes. See also **Figure S6C** and **S6E**. **(E)** CYP2A6 in the presence of NADPH, and under aerobic conditions, produced measurable amount of acid-metabolite, HNA, detected and quantified as a glutathione (GSH) adduct (termed, GS-HNA), against indicated controls (i.e., in the absence of CYP2A6, or HNE). ***Note***: HNE can undergo inadvertent oxidation in the absence of CYP2A6, oftentimes resulting in aberrant background signals. Quantification here shows combined datasets across 6 independent biological replicates involving substrate concentrations indicated in **Data 3E**. Y-axis represents relative percent of GS-HNA detected over GS-HNE in each replicate. P-values calculated by Dunnett’s multiple comparisons test. **(F)** Indicated phenotypic changes were quantitatively scored for day 1 adult worms following acute HNE stress (2 mM, 1 hour). **(G)** Average defecation motor program (DMP) intervals of day 1 adult worms, in response to electrophilic stress [HNE(alkyne), 2 mM, 1 h] vs. DMSO, in the presence of either cyp-33e1 RNAi or RNAi Control (Ctrl., L 4440). Each data point represents average value of 5 cycles from individual animals. P-values calculated by Tukey’s multiple comparisons test. See also **Figure S6F-S6H**; **Movie 2-7** in **Data S6**. **(H)** Average DMP intervals of L4 worms (48 h after seeding to RNAi plates) (WT or cyp-33e1 mutant knock-in), measured in the absence of DMSO or electrophilic stress. Each data point represents the 6-9 (average 7) cycles from individual animals. P-values calculated by Dunnett’s multiple comparisons test. See also **Method S1**; **Figure S6I**; **Movie 8-10** in **Data S6**. See **Method S2B** and **S2C** for validations of knock-in strains. **(I)** Average DMP intervals of day 1 adult worms (WT or cyp-33e1 mutant knock-in), in response to HNE(alkyne) (2 mM, 1 h). Each datapoint represents 4-7 (average 6) cycles from individual animals. P-values calculated by unpaired Students’ t-test. See also **Method S1**; **Figure S6J**; **Movie 12-16** in **Data S6**. See **Method S2B** and **S2C** for validations of knock-in strains. All data present mean±SEM. Sample sizes and decision tree for statistical treatment in **Method S3**.

In the *C. elegans* protein cyp-33e1, both single mutants intercepted HNE under T-REX (**Figure 5A and S6A**), indicating that C439 and C295 sense HNE. cyp-33e1(C295A, C439A) showed minimal labeling, indicating that these two cysteines account for the majority of HNE sensing. Likewise, C82 and C439 in CYP2A6, but not C14, were kinetically-privileged sensing sites (**Figure 5B** and **S6B**). In parallel, mass-spectrometry-based site-specific modification analysis of CYP2A6, exposed to sub-micromolar HNE, revealed HNE-derived modification at C82 and C439 (Data S3C and S3D). No modification occurred at C14, consistent with T-REX (**Figure 5A**, **5B, S6A** and **S6B**). Thus, there are two HNE-sensing functions within cyp-33e1/CYP2A6: one occurs at a conserved active-site cysteine (C439); the other occurs at a residue that has shifted during evolution (C295^worm^/C82^human^).

We next determined how each HNE-sensing site contributes to partial inhibition of oxidoreductase activity observed above (**Figure 4C** and **S5F**). Preparation of purified cyp-33e1 has not been previously reported. Despite numerous attempts, we were unable to isolate sufficient quantities of cyp-33e1 for enzymatic assays. However, we were able to purify human microsomal CYP2A6 wt and the 3 relevant mutants: C82A, C439A, and (C82A, C439A) double mutant, using HEK293+Hycell expression system (**Figure S6C**). Mutation of the catalytic-site cysteine, C439, rendered CYP2A6(C439A) and CYP2A6(C82A, C439A) inactive (**Figure 5C** and **S6D**). CYP2A6(C82A) exhibited activity similar to CYP2A6(wt) (**Figure 5C** and **S6D**). Under similar HNE dosage and treatment times as deployed for commercial CYP2A6(wt) above (**Figure 4C**), our in-house-expressed CYP2A6(wt) was inhibited (**Figure 5D** and **S6E**), similarly to what we observed for the commercial protein (**Figure 4C**). Under identical conditions, CYP2A6(C82A) was comparably sensitive to HNE (**Figure 5D** and **S6E**). These data affirm that partial suppression of oxidoreductase activity occurs through HNEylation at the conserved catalytic-site cysteine, C439.

HNE is also potentially a CYP2A6 substrate. Indeed, incubation of HNE with the enzyme in the presence of NADPH, in an aerobic environment, produced detectable amounts of HNA. When the enzyme was omitted from the same assay mixture, HNA production was insignificant compared to background control (i.e., in the absence of HNE) (**Figure 5E**; **Data S3E**). Thus, HNE in the active site of CYP2A6 undergoes partitioning between enzymatic oxidation, and labeling the active site.

### Cyp-33e1 is an essential mediator of intestinal physiology in both non-stressed and stressed states

We next used *CRISPR-cas9* technology to generate homozygous cyp-33e1(C295A) and cyp-33e1(C439A) knock-in (KI) *C. elegans*. Following back-crossing into wild-type nematodes, each strain was validated by sequencing using single-worm PCR (**Method S2A-C**). The KI animals grew comparably to wild-type worms. In parallel, we undertook a systematic quantitative assessment of *C. elegans* phenotypes to identify those most prominently affected by acute electrophilic stress (**Figure 2E: Path II**). Between food race, body bends/minute, viability, brood size, and DMP cycles, DMP emerged to be most affected by acute HNE stress (**Figure 5F**). As DMP cycles in *C. elegans* also report on intestinal physiology^64^ (**Figure S6F**; **Movie 1** in **Data S6**) and cyp-33e1 was mapped by OS-Localis-REX as a gut-responsive sensor (**Figure 1D**), we chose DMP to investigate the effects of altering cyp-33e1’s electrophile responsivity on *C elegans*.

Comprising three steps, DMP consists of highly-coordinated periodic rhythms accurately quantifiable from real-time measurements. We first evaluated the effects of cyp-33e1 depletion on DMP. RNAi-control young adults underwent 63.7±1.7 s intervals (**Figure 5G** and **S6G**; **Movie 2-3** in **Data S6**), within the expected range for wild-type *C. elegans* at this age^65^. Cyp-33e1 depletion reduced DMP cycle length (**Figure S6G**; **Movie 3** in **Data S6**). Following whole-animal electrophile stimulation, DMP in control animals was significantly perturbed, reducing the mean cycle length to 58.5±0.8 s (**Figure 5G** and **S6H**; **Movie 4-5** in **Data S6**). Cyp-33e1-deficient animals did not undergo further reduction of cycle length following HNE treatment (**Figure 5G** and **S6H**; **Movie 6-7** in **Data S6**). These results demonstrate: **(i)** that cyp-33e1’s electrophile-sensing activity plays a functional role in natural oscillations of DMP and likely overall gut homeostasis; and **(ii)** that HNE exerts its effect on DMP in a cyp-33e1-specific manner.

We subsequently compared changes in DMP cycles in cyp-33e1(C295A) and cyp-33e1(C439A) KI *C. elegans*, with and without acute HNE exposure. Neither KI strain showed significant decrease in DMP (**Figure 5H** and **S6I**; **Movie 8-10** in **Data S6**). This result is in contrast to the decrease in defecation interval upon cyp-33e1 RNAi [**Figure 5G** (1^st^ vs. 3^rd^ column) and **S6H**; **Movie 2-3** in **Data S6**]. These data imply that changes in defecation interval in cyp-33e1 RNAi nematodes are not ascribable uniquely to loss of cyp-33e1 activity. Moreover, *both* KI strains emerged to be recalcitrant to HNE-induced reduction of DMP-cycle length observed in wild-type animals (**Figure 5I** and **S6J**; **Movie 11-16** in **Data S6**). As cyp-33e1 RNAi and both KI strains are refractory to HNE-induced changes in DMP cycles (**Figure 5G**: 3^rd^ & 4^th^ columns, **Figure 5I**), electrophile sensing by cyp-33e1 at *both* electrophile-sensing sites is important for reduction in defecation intervals during stress.

### Electrophile stress promotes defects in lipid accumulation dependent on cyp-33e1 activity

HNE is a prototypical lipid-derived electrophile upregulated under physiological stress by non-enzymatic and enzymatic lipid peroxidation^6^. Fat metabolism defects are also correlated with DMP defects^29^. We thus next investigated the effect of HNE treatment on lipid storage. Using a lipid-droplet reporter strain^66^ (DHS-3::GFP, **Figure S6K**, Key Resources Table), HNE treatment depleted lipids (**Figure 6A** and **6B**). cyp33e1-RNAi did not induce lipid depletion (**Figure 6B**, 1^st^ vs. 3^rd^ columns). However, cyp-33e1-RNAi worms were refractory to HNE-induced lipid-level changes (**Figure 6B**). ORO staining of control RNAi (wild-type N2) and the corresponding cyp-33e1-RNAi worms, gave similar outcomes (**Figure 6C** and **S6L**). Thus, cyp-33e1-deficient animals marshall lipid-distribution responses to HNE differently from controls. However, cyp-33e1(C439A) KI animals were refractory to HNE-induced lipid depletion (**Figure 6D**, last two columns, and **Figure S6M**), whereas cyp-33e1(C295A) KI strain behaved similarly to wild-type (**Figure 6D**, first four columns, and **Figure S6M**). Indeed, cyp-33e1(C439A) KI strain exhibited reduced global lipid content even in non-stressed states (**Figure 6E**, compare within ‘DMSO’ set, and **Figure S6M**); cyp-33e1(C295A) KI strain exhibited similar lipid levels to wt *C. elegans* (**Figure 6E**, compare within ‘DMSO’ set, and **Figure S6M**). The refractory nature of strains lacking active cyp-33e1 to HNE highlights the importance of cyp-33e1 activity in the maintenance of global lipid storage during stress. Altogether, differences between cyp-33e1(C439A) KI and cyp-33e1 RNAi backgrounds in the absence of exogenous HNE (**Figure 6E**, 1^st^ and 3^rd^ columns, vs. **Figure 6B** and **6C**, 1^st^ and 3^rd^ columns), further demonstrate that there are regulatory modes of cyp-33e1 that are not dependent on activity. However, these modes of regulation are outside the remit of the present study, as they are independent of HNE.

**Figure 6.**
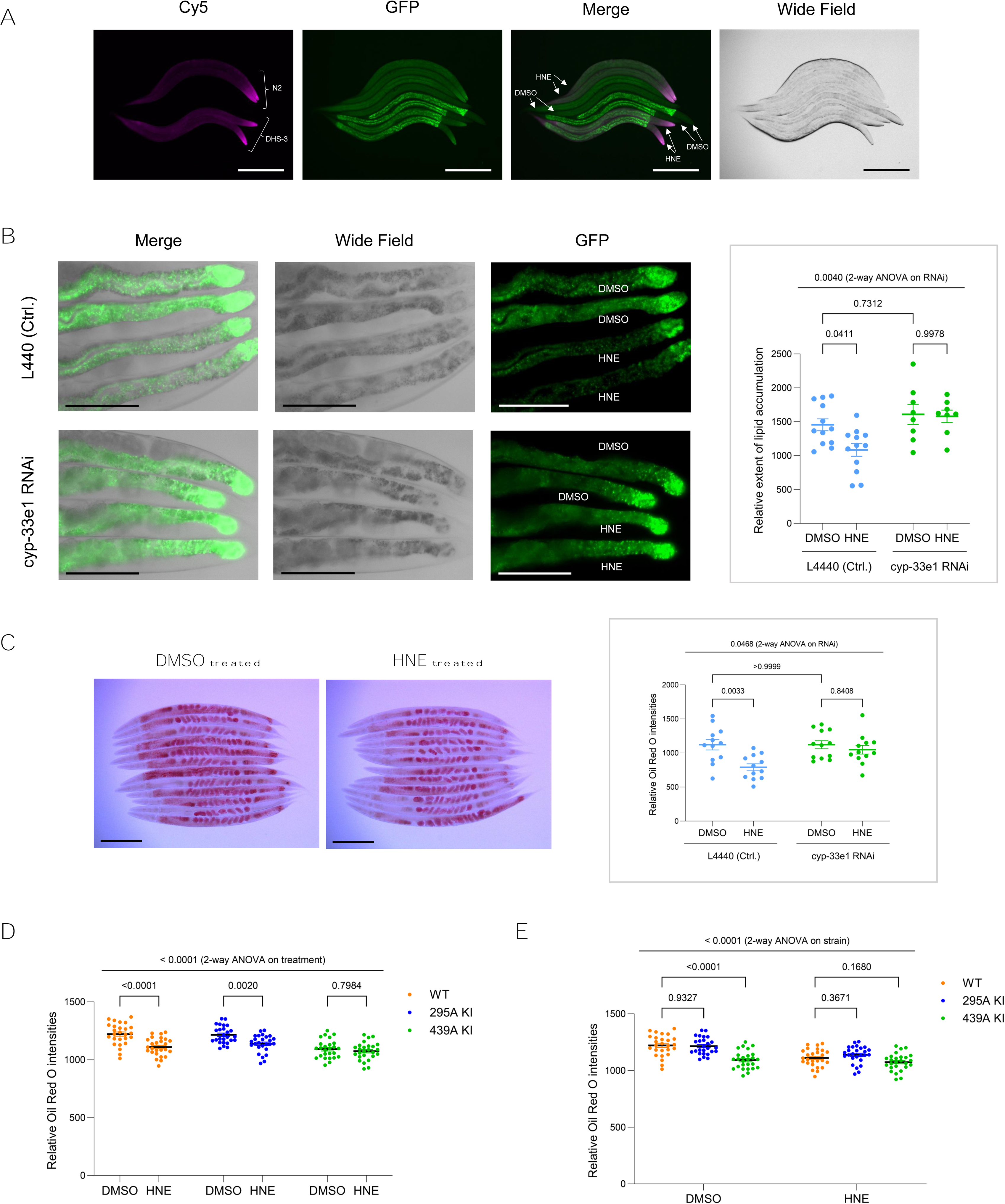
Electrophile stress disrupts lipid homeostasis: these effects are cyp-33e1—specifically, catalytic-cysteine (C439)—dependent. **(A)** Click-Immunofluorescence imaging of paraformaldehyde-fixed animals showed effective uptake of alkyne-functionalized electrophile (following Click coupling with Cy5-azide), subsequent to whole-animal electrophile stimulation [HNE(alkyne) 2 mM, 1 h]. 8 x L4-young adult hermaphrodites—4 x DHS-3::GFP reporter worms (Key Resources Table), 4 x wild-type N2 worms for comparison—are shown. ***Note*:** Elevated HNEylation in the head is likely due to preferential electrophile uptake with food. This issue was circumvented by the use of tissue-specific-Halo-targetable photocaged-HNE, see **Figure S1C** and **S1D**). Scale bar: 200 µm. See also **Figure S6K**. **(B)** Acute electrophile stress [HNE(alkyne), 2 mM, 1 h] impairs lipid accumulation: cyp-33e1 deficiency ablates this phenotype. 4 x day 1 adult DHS-3::GFP hermaphrodites were aligned manually, post levamisole immobilization. Top 2 worms: DMSO treated; bottom 2 worms: electrophile treated. Representative posterior intestinal regions are shown. Scale bar: 100 µm. ***Inset***: quantification derived from total intestinal GFP-signal intensity. P-values calculated by Tukey’s multiple comparisons test. **(C)** Oil Red O staining validated lipid depletion, following acute electrophilic stress [HNE(alkyne), 2 mM, 1 h]: 12 fixed day 1 adult worms were manually aligned. Scale bar: 200 µm. ***Inset***: quantification. Signals were quantified by circling the entire animal except for the pharyngeal region. P-values calculated by Tukey’s multiple comparisons test. See also **Figure S6L**. **(D)** WT and indicated knock-in mutant animals were subjected to the same conditions as Figure 6c. P-values calculated by Šidak’s multiple comparisons test. See also **Figure S6M**. **(E)** Results in Figure 6D presented/analyzed with different groupings. P-values calculated by Dunnett’s multiple comparisons test. All data present mean±SEM. Sample sizes and decision tree for statistical treatment in **Method S3**.

### Cyp-33e1 oxidation of HNE triggers global lipid deficit

Since cyp-33e1 activity emerged to be crucial for HNE-induced lipid storage changes, we hypothesized that either reductive or oxidative metabolism of HNE, mediated by cyp-33e1, initiates lipid deficit. Given the simplicity of HNE, two main cyp-33e1-catalyzed metabolites are possible, HNA (oxidation) and DHN (reduction). Both metabolites were synthesized and their homogeneity was characterized using NMR spectroscopy (**Method S3**). Feeding RNAi-control nematodes with HNA, but not DHN, phenocopied HNE-induced lipid depletion in DHS-3::GFP lipid-reporter worms (**Figure 7A**, left set, and **Figure S7A**). In cyp-33e1-deficient reporter worms, HNE-stressed animals did not undergo changes in lipid accumulation (**Figure 7A**, right set), consistent with our data above (**Figure 6B** and **6C**). In cyp-33e1-RNAi reporter worms, DHN had no effect (**Figure 7A**). However, HNA feeding to cyp-33e1-RNAi fed worms caused lipid depletion (**Figure 7A**, right set, and **Figure S7A**). This observation was further validated by ORO staining (**Figure S7B** and **S7C**). These data are consistent with cyp-33e1 converting HNE to HNA, that directly interferes with lipid storage. Linking these data back to effects on DMP, HNA administration to wild-type N2 nematodes reduced defecation intervals similarly to HNE administration (**Figure 7B**; **Movie 11**, **12**, and **16** in **Data S6**). Thus, cyp-33e1 activity on HNE, produces HNA, that ushers lipid depletion that is likely important for HNE’s effects on DMP activity.

**Figure 7.**
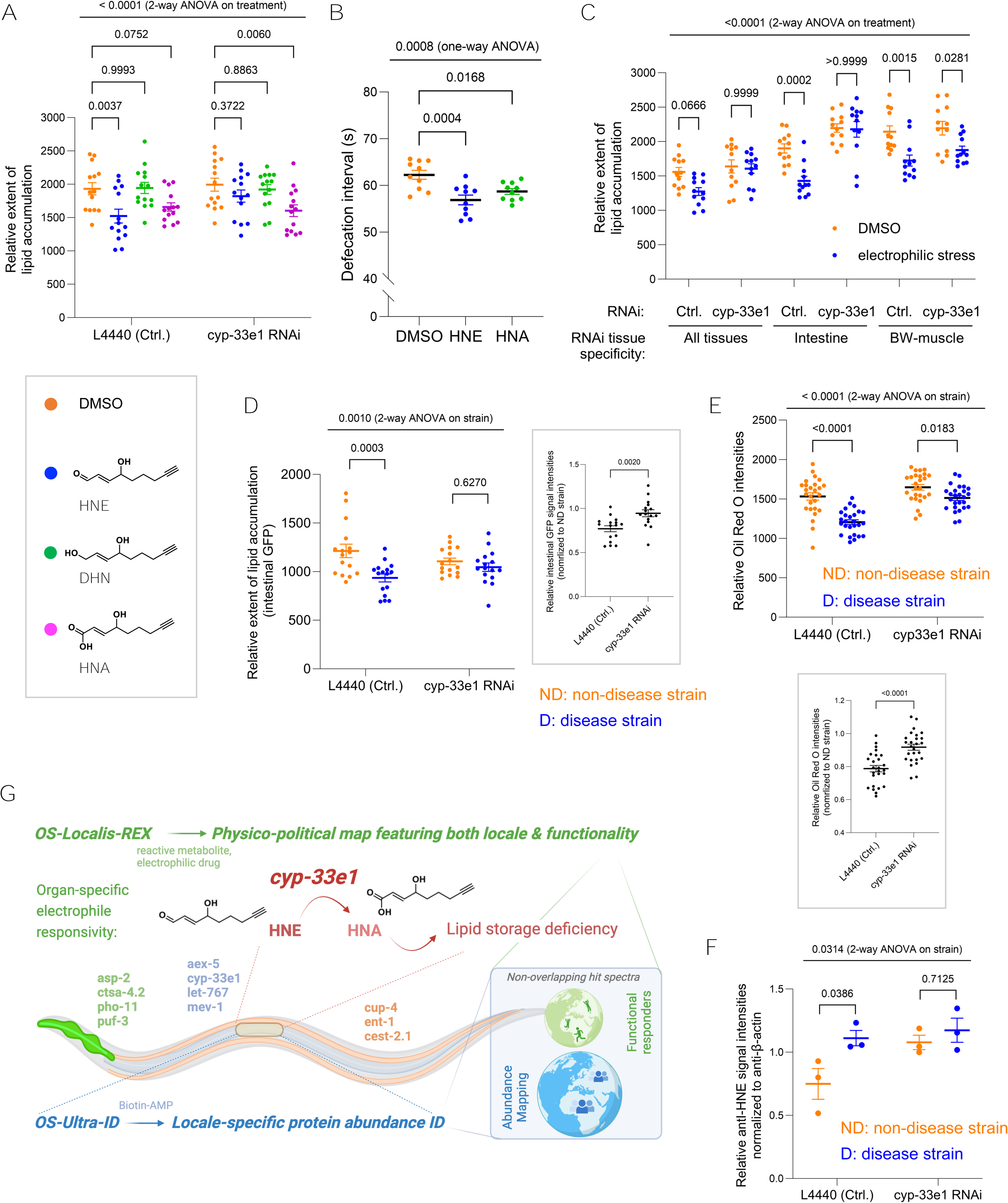
cyp-33e1 HNE sensing generates a metabolite (HNA) that promotes lipid depletion; similar results are found in diseased models with heightened endogenous HNE stress. **(A)** Administration of HNA, an enzymatic product of cyp-33e1, phenocopied HNE-induced lipid depletion in the wild-type (L4440, Ctrl.) animals, and maintained this phenotype in the presence of cyp-33e1 RNAi. P-values calculated by Dunnett’s multiple comparisons test. See also **Figure S7A**. ***Inset***: chemical structures of small molecules. See **Figure S7B** and **S7C** for results obtained using an orthogonal Oil Red O staining. **(B)** Average DMP intervals of live worms (WT, or indicated cyp-33e1 mutant knock-in), following HNE(alkyne) (2 mM, 1 h) treatment. Each data point represents 4-7 (average 6) cycles from individual animals. P-values calculated by Dunnett’s multiple comparisons test. Data from first 2 columns are replicas of the first 2 columns within **Figure 5H**. See also **Movie 11**, **12**, and **17** in **Data S6**. **(C)** Despite similar abundance of cyp-33e1 in both intestine and BW-muscle tissues (**Figure 1D**), BW-muscle-specific depletion of cyp-33e1 failed to block stress-induced lipid depletion, observed during ubiquitous and intestine-specific cyp-33e1 knockdown. Ctrl.: control RNAi. P-values calculated by Šídák’s multiple comparisons test. See also **Figure S7D; Method S2D-G**. **(D)** Quantification of relative lipid droplet extent in indicated diseased (D) and non-diseased (ND) animals. See also **Figure S7G** and **S7H**, Key Resources Table. P-values calculated by Šídák’s multiple comparisons test. ***Inset*** shows comparison between fold change in ND versus D strains for control and cyp-33e1 RNAi worms. **(E)** Similar to **Figure 7D** but deployed Oil Red O staining. P-values calculated by Šídák’s multiple comparisons test. See also **Figure S7I**. ***Inset*** (bottom) shows a comparison between fold change in ND versus D strains for control and cyp-33e1 RNAi worms. **(F)** Quantification of the extent of HNEylated proteomes in D vs. ND animals subjected to cyp-33e1 RNAi or L4440 (Ctrl.) RNAi. P-values calculated by Šídák’s multiple comparisons test. See also **Figure S7E** and **S7F**. **(G)** A combination of profiling electrophile responders [using organ-specific precision localized electrophile generation (OS-Localis-REX)] and mechanistic investigations, provides a detailed annotated map of *chemically actionable* targets within specific organs in *C. elegans*. ***Inset*** (bottom right) shows effectively non-overlapping target spectra between OS-Localis-REX and OS-Ultra-ID (organ-specific mapping of localized protein abundance, performed in the same 3 organs as in OS-Localis-REX). All data present mean±SEM. Sample sizes and decision tree for statistical treatment in **Method S3**.

### Tissue-specific electrophile responsivity of cyp-33e1

At this juncture, because cyp-33e1 is present in numerous locales at similar levels (**Figure 1D**), it remains unclear from our data to what extent cyp-33e1 activity must localize in the gut to exert its effect on lipid storage. We used tissue-specific RNAi^67^ to address this question. We first generated and validated the strains necessary to execute tissue-specific gene silencing, in either intestine or BW-muscles (**Method S2D-G**). Ubiquitous cyp-33e1 knockdown (**Figure 7C**, left set) reproduced our previous observations (**Figure 7A**, left set). BW-muscles-specific cyp-33e1 depletion was did not block stress-induced lipid depletion (**Figure 7C**, left vs. middle sets, and **Figure S7D**). Intestine-targeted cyp-33e1-RNAi blocked lipid depletion (**Figure 7C**, left vs. right-most sets, and **Figure S7D**). Thus, HNE-induced lipid depletion is mediated by intestinal cyp-33e1.

### Physiological significance of cyp-33e1 in animals with elevated endogenous stress

We exploited *C. elegans* harboring the transgene [unc-54p::Htt513(Q128)::YFP::unc-45 3’UTR]^23–25,68^, expressing YFP-tagged polyQ-expanded disease-associated fragment of human Hungtinton’s disease (HD) protein Htt (Key Resources Table). Endogenous HNE production and proteome HNEylation are upregulated in diseased animals compared to Htt513(Q15) expressing non-disease animals. We crossed both strains into the DHS-3::GFP lipid reporter line (**Method S2H**). Anti-HNE western blots confirmed elevated protein HNEylation, against age-matched healthy controls (**Figure S7E**: two left-hand lanes). Oxy-blot, a measure of overall protein carbonylation, showed minimal changes between diseased and control animals (**Figure S7F**). Thus, HNEylation is an operative stressor in diseased worms.

The resulting double transgenic lines showed hallmarks of Htt513 aggregation only in diseased worms (**Figure S7G**). Comparison of lipid droplet levels in these two strains when treated with control RNAi showed that there was a depletion of lipid droplets in diseased strain, compared to the control strain (**Figure 7D** and **Figure S7H**: top). There was no difference between these strains when they were treated with cyp-33e1 RNAi (**Figure 7D** and **Figure S7H**: bottom). Similar data were obtained with ORO staining (**Figure 7E**, **Figure S7I**), although in this instance, cyp-33e1-RNAi caused an increase in lipid accumulation selectively in the diseased strain (**Figure 7E**, compare 2^nd^ and 4^th^ columns). This observation further reinforces the importance of cyp-33e1 activity in HNE-induced depletion of lipid accumulation. To determine if these outcomes could be explained simply by total protein HNEylation, we examined HNEylation in these backgrounds. Following cyp-33e1 RNAi, the difference in the extent of HNEylation between diseased and healthy animals was no longer apparent (**Figure 7F**, **Figure S7E**: two right-hand lanes). Thus proteome HNEylation is not correlated with changes in lipid storage; activity of cyp-33e1 is correlated with such changes.

## Discussion

Several lines of evidence indicate that our spatiotemporal *and functional* proteomics approach identified genuine locale-specific responders. First, hits were not dominated by expression: in several instances, reported and our own tissue-specific expression datasets indicated that proteins sensed electrophiles in locales where they were of reduced expression than other locales investigated. Second, and no less remarkably, our hit proteins did not gravitate to locale-indiscriminate electrophile responders, indicating that OS-Localis-REX is indeed able to undertake organ-specific actionability profiling, surmounting issues that have to date hindered development of *functionally-annotated* maps. These results further point to there being significant context dependence among chemically-actionable proteins across organs. These observations could reflect changes in responder proteins’ interactomes, locales, modifications, or other context-dependent regulatory nuances across tissues. All the above information is indeed difficult to extract by other proteomics/sequencing methods, including APEX, and as shown here, OS-Ultra-ID. Clearly, as our own data with OS-Ultra-ID-expressing worms show, OS-Localis-REX cannot replace the power of such methods to identify locale-specific proteomes, because it is blind to proteins that have little chemical function. Hence OS-Localis-REX is complementary to general profiling methods. However, as we outline in the introduction, it is often functionality or actionability that constitutes the most important parameters. Moreover, with an expanding toolset of proto-electrophiles applicable to REX technologies, a wealth of activities can be probed using the same approach, likely extending the possibilities of finding locale-specific actionable proteins.

Our investigations into the role of context-dependent responders uncovered several known and novel phenotypic effects associated with OS-Localis-REX hits. Several were also newly linked to electrophile-induced effects on egg-laying rate. Such phenotypic changes in electrophile response are, we submit, compelling evidence that OS-Localis-REX functionally-annotated maps report on validated stress responders whose electrophile engagement is linked to how worms manage electrophilic challenges.

One specific responder, cyp-33e1, stood out. Subsequent mechanistic examinations documented that this metabolic enzyme manifests context-dependent roles in regulating gut health and response to electrophilic stress. cyp-33e1 thus emerged to be responsible for converting HNE to a signal, acid metabolite (HNA), ushering lipid deficit. This behavior is not positively linked to cyp-33e1 HNEylation (it is antagonistic, in vitro). These results demonstrate that our method has the power to identify proteins that are metabolically active against specific electrophiles, providing that there is partitioning between metabolism and labeling.

Overall this result adds a new dimension to our profiling data, and further underscores the ability of OS-Localis-REX to decode cellular signaling pathways enacted by enzymatic processes and their specific products at the tissue-specific level. This is far from a trivial discovery. Cyp-isoform-induced sensitization of molecules causing toxicity and disease, such as cancer, is well known. This occurs, for instance, in drug metabolism^60^, and inadvertent toxification of xenobiotics, e.g., methanol (acidosis) and benzene (alkylation). However, several Cyp-isoforms are also involved in reactive small-molecule signaling pathways, not unakin to what we delineate here. Such pathways in mammals, from where the majority of these data are derived, include CYP1-4 families that metabolize eicosanoids. Regulatory loops for other CYP family enzymes have been proposed^69^: e.g., several CYP enzymes, including CYP26 family enzymes, metabolize *all trans* retinoic acid^70^. It is interesting to note that retinoic acid is an important regulator of hepatic lipid homeostasis^70,71^. It is thus tempting to propose that CYP metabolism of lipophilic entities may form regulatory loops controlling lipid storage, and hence regulate response to stresses which are *broadly* conserved across taxa. Unfortunately, despite reports of several *C. elegans* cyp-enzymes performing lipid-metabolism roles^72^, precise molecular and pathway details are lacking. As we demonstrate here, localized reactive-lipid generation may be one way to progress in this area.

<u>Limitations of the Study</u>. As noted above, OS-Localis-REX is blind to total protein levels in specific tissues. For the moment, at least, these parameters are admittedly the most commonly used, although our data question their overall utility. We note that combination of OS-Ultra-ID with OS-Localis-REX is perhaps an ideal solution to this issue, as we demonstrate here. Moreover, as the electrophile released during OS-Localis-REX is relatively low concentration, our hits are limited to the very best responders. It is possible that modification of the REX protocols^19,20^ may enable more depth to be achieved in a single experiment, without resorting to using a fleet of photocaged electrophiles.

At the biochemical level, we cede that how HNA functions to reduce lipid droplets remains to be defined. Likely this molecule interacts with (a) lipid receptor(s). However, as HNA is a non-reactive molecule, it is not applicable to REX technologies, and indeed overall methods to study such interactomes are lacking. HNA befited with photo-crosslinkers and similar strategies are quite likely to be of use to achieve this goal, but are outside the scope of this study.

Bearing these limitations in mind, subsequent work investigating such behaviors, and indeed profiling other tissues, will help establish a more detailed understanding of how locale-specific responders vary across worms. Similar profiling experiments across other organisms will bring further insights that can be compared across organisms/expression patterns, to give a selective comprehensive lens into actionable protein targets against the local proteome noise.

## Supporting information

All SI figures

## Acknowledgements.

Prof. Pierre Gönczy, Ms. Coralie Busso (EPFL-SV): RNAi clones (Ahringer’s library). Prof. Mario de Bono, Ms. Ekaterina Lashmanova (IST:Austria): MG1655:BioBKan strain. Drs. Krishna Tripathi, Kevin Fridianto, Chaosheng Luo (Aye Lab): photocaged-electrophile probe synthesis. <u>Research</u>: ERC:CoG financed by SERI, Switzerland (Y.A.) (2023-present); SNSF:310030_184729 (Y.A.) (2019-2023); SNSF:R-Equip (Y.A.) for EPFL proteomics facility instrumentation (2023); SNSF:NCCR master program (J.L.) (2019); Novartis Foundation (M.J.C.L.).

## Author Contributions

J.L., performed all experiments (except photocaged-electrophile synthesis and transgenic/knock-in worm generations), data analysis, compilation of figures, legends, and methods; A.K., collaborated with J.L. on phenotypic assays, RNAi validations involving initial 15 genes; Y-Q.G. collaborated with J.L. on phenotypic assays, MS validations of protein modification, small-molecule MS analysis, compilation of methods; D.A.U., initial development of *C. elegans* Localis-REX workflows; R.H. collaborated with J.L. and Y.A. on TMT-proteomics experiments, and with J.L., Y-Q.G., and Y.A. on MS-PTM mapping. B.A.N., collaborated with J.L. and A.K. on phenotypic assays involving initial 15 genes; M.J.C.L. experimental design and data analysis, writing (manuscript critical review and editing); Y.A. conceptualization, project development and administration, experimental design and data analysis, writing (manuscript critical review and editing), supervision, funding acquisition. All authors assisted with manuscript final proofing.

## KEY RESOURCES TABLE

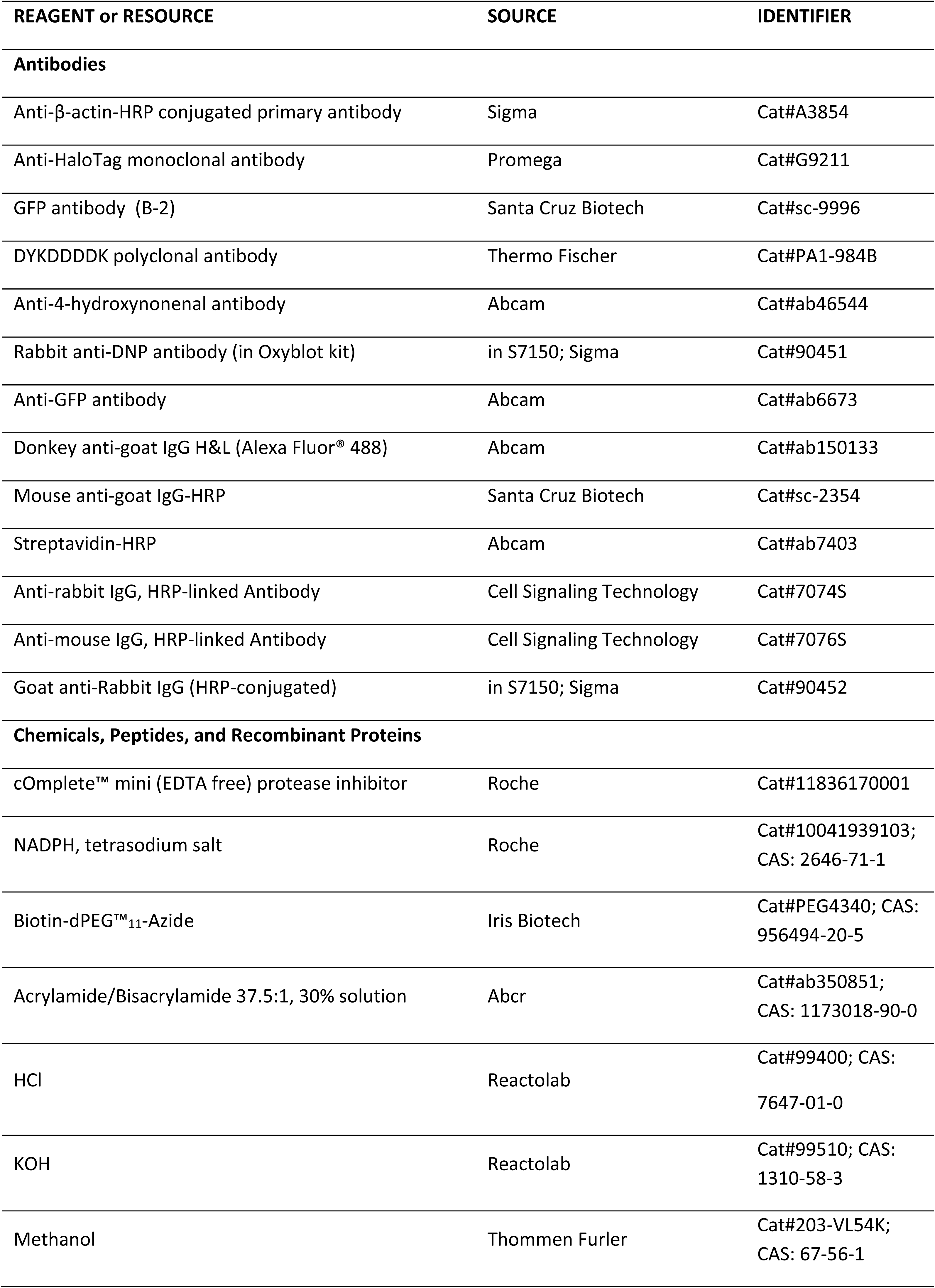
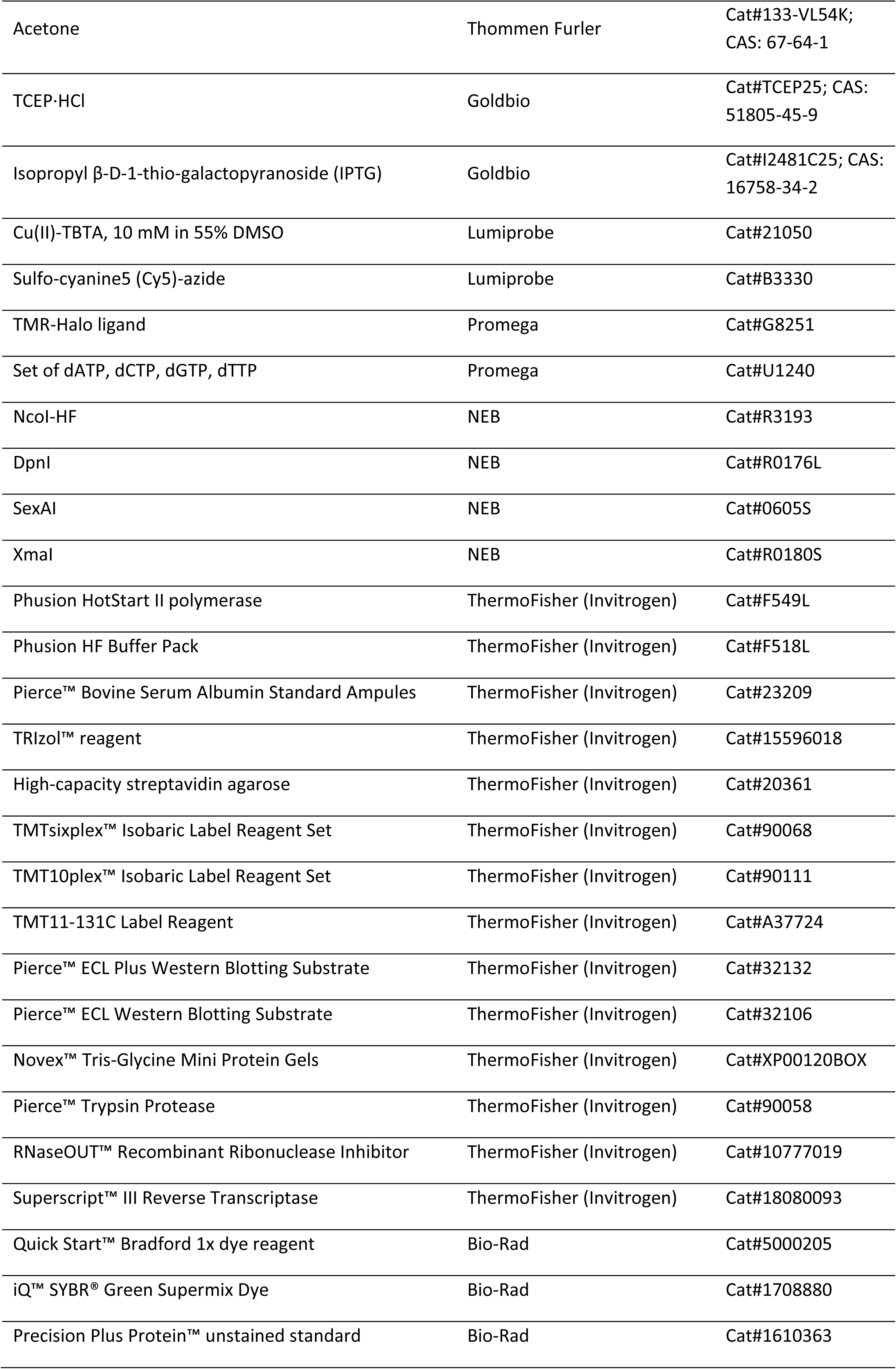
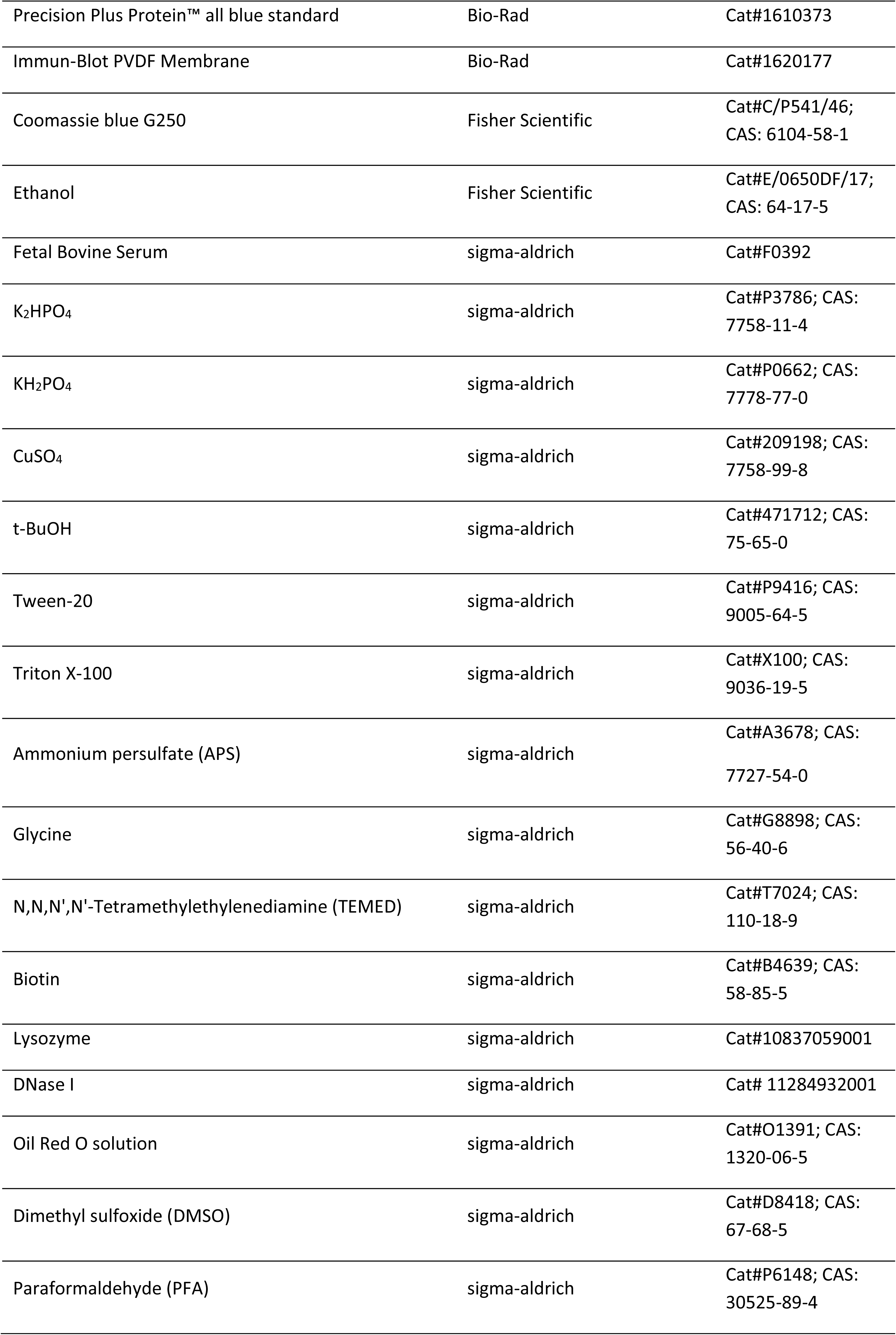
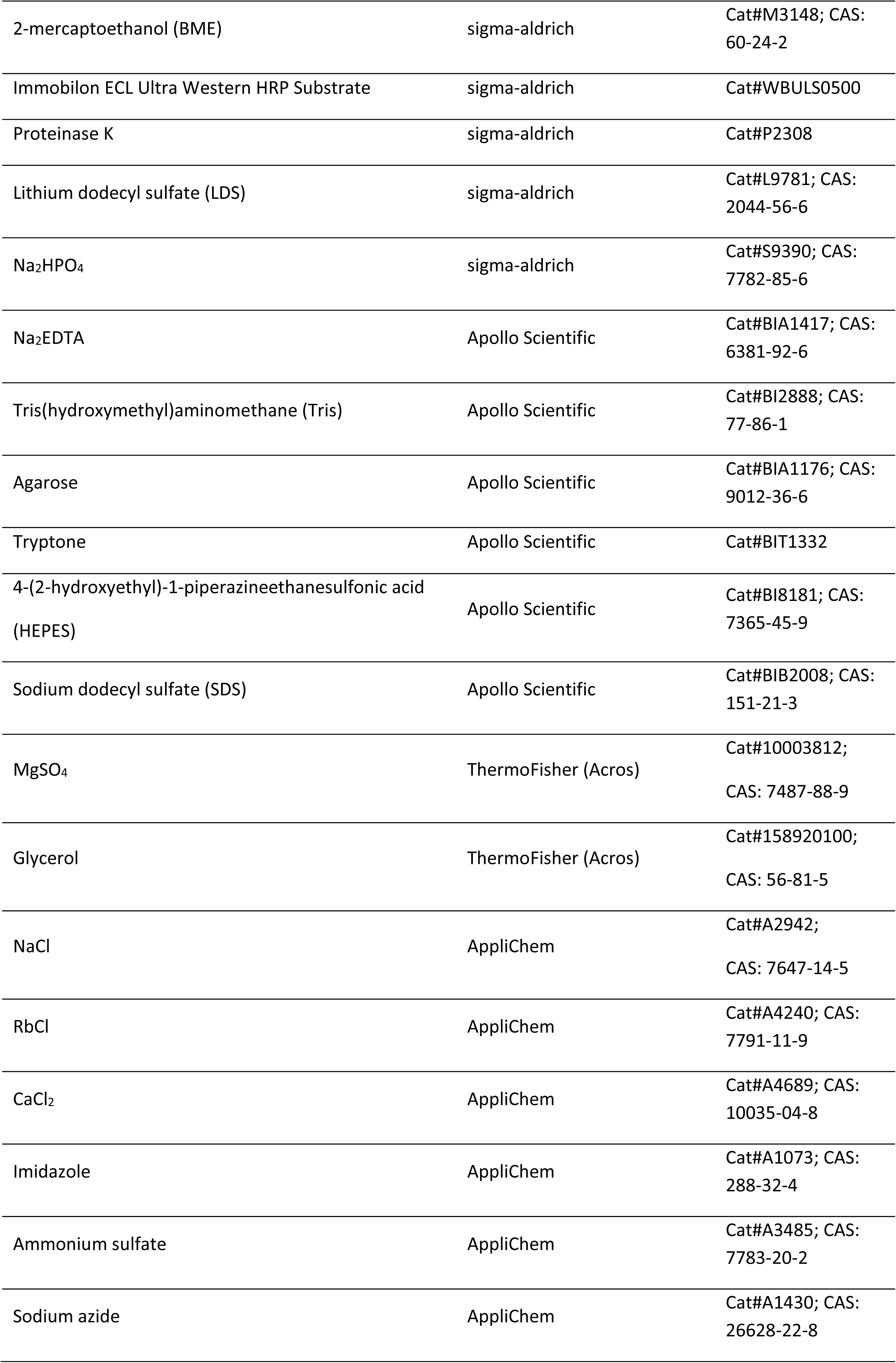
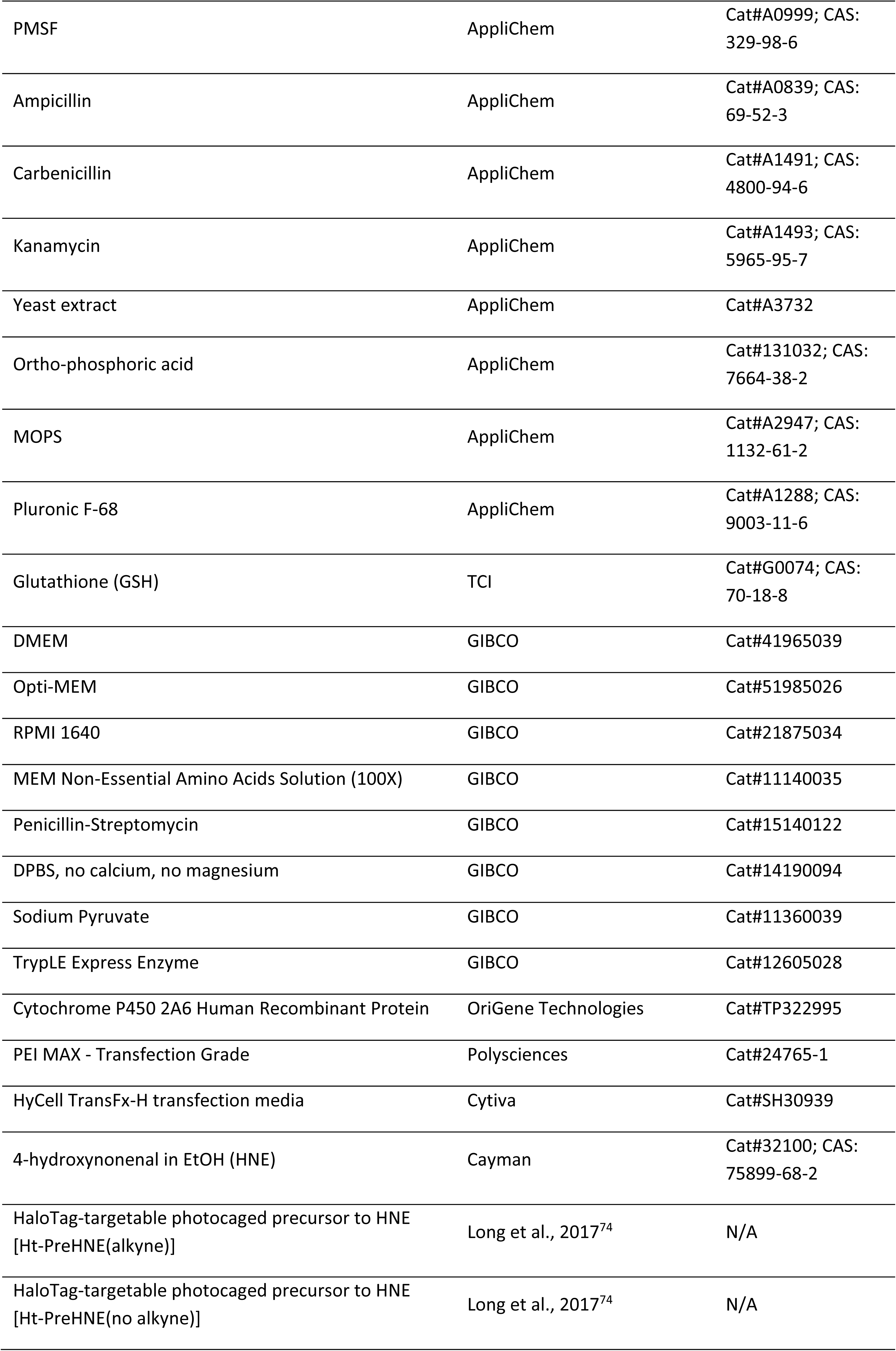
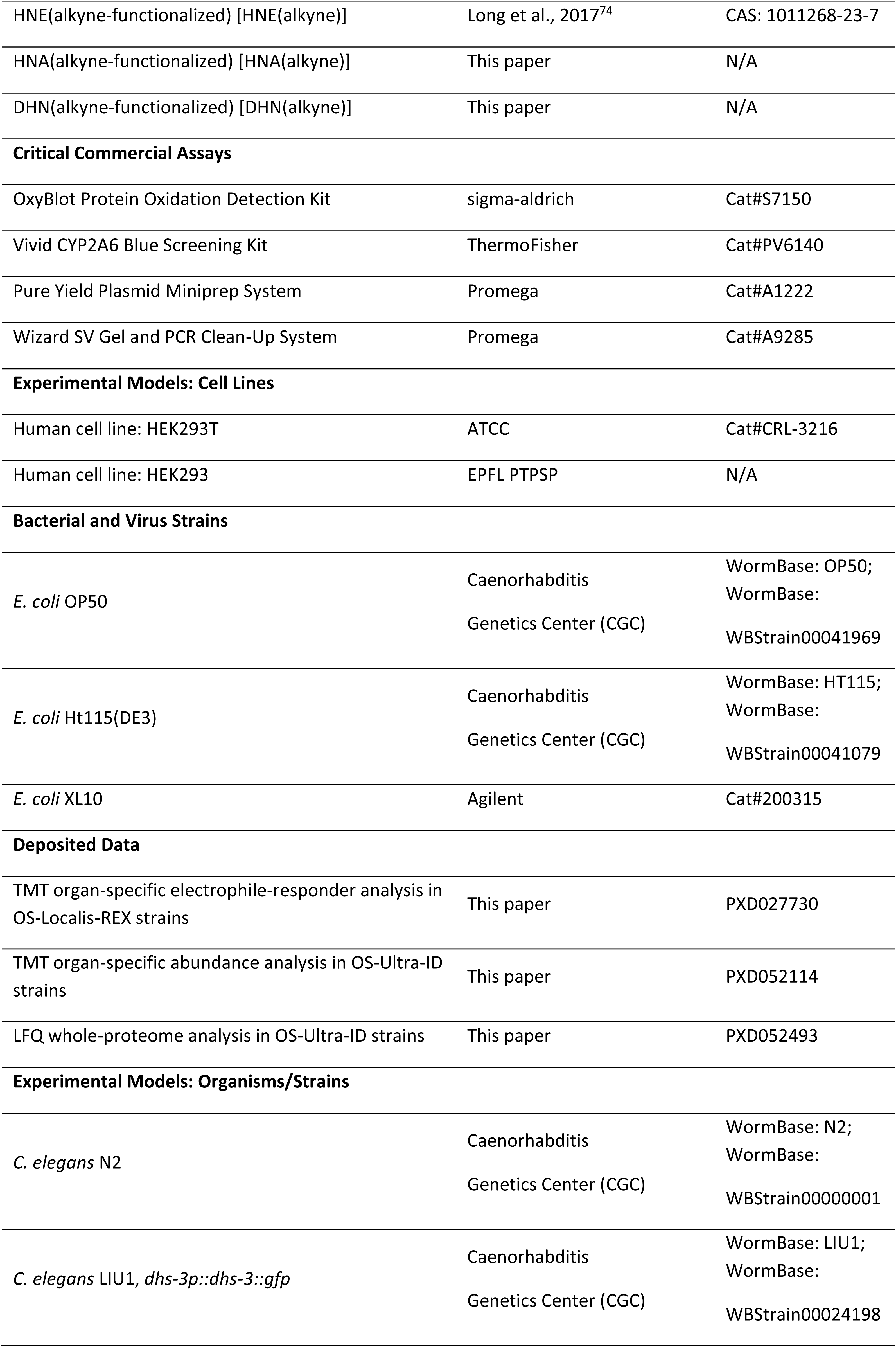
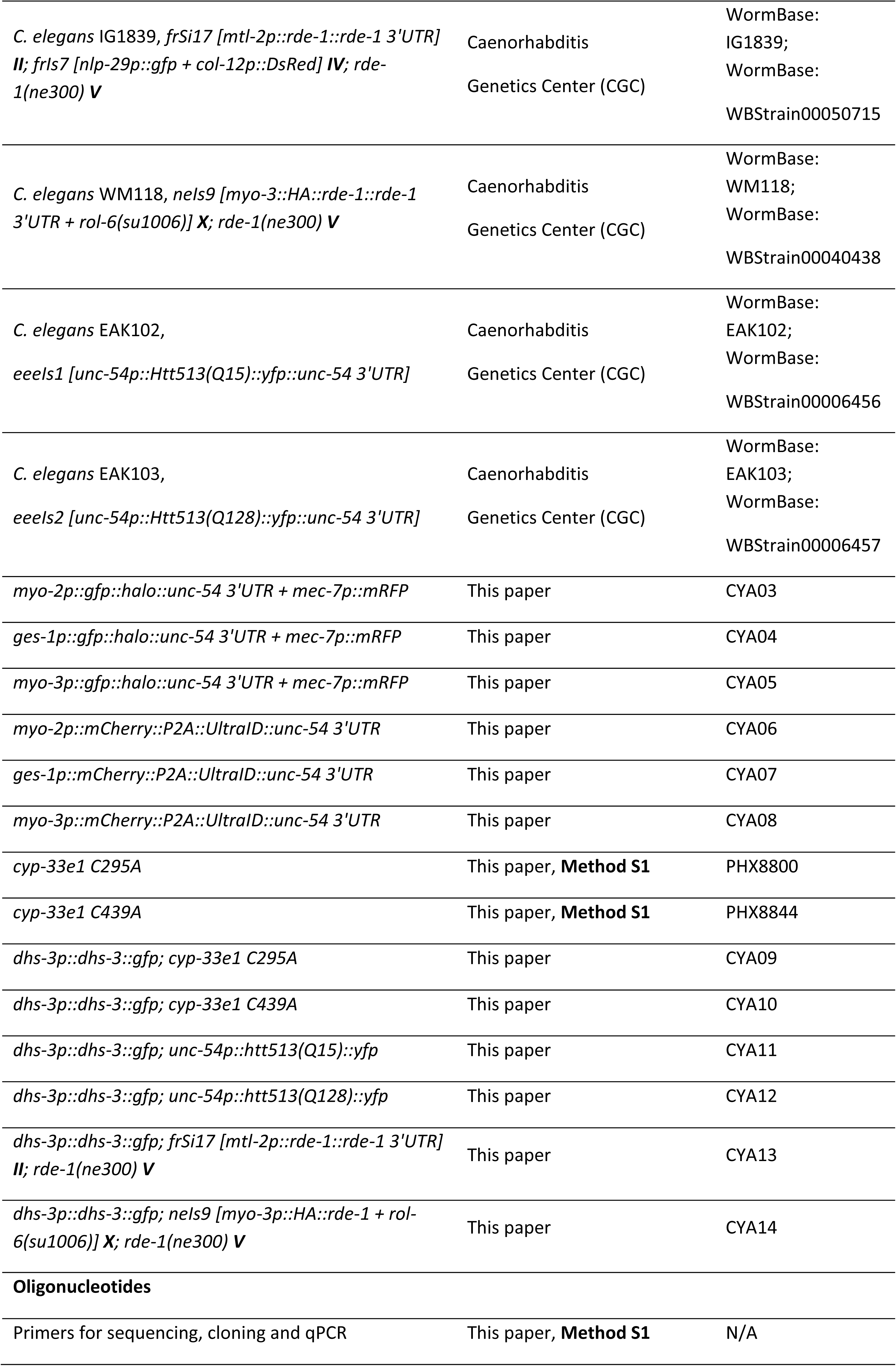
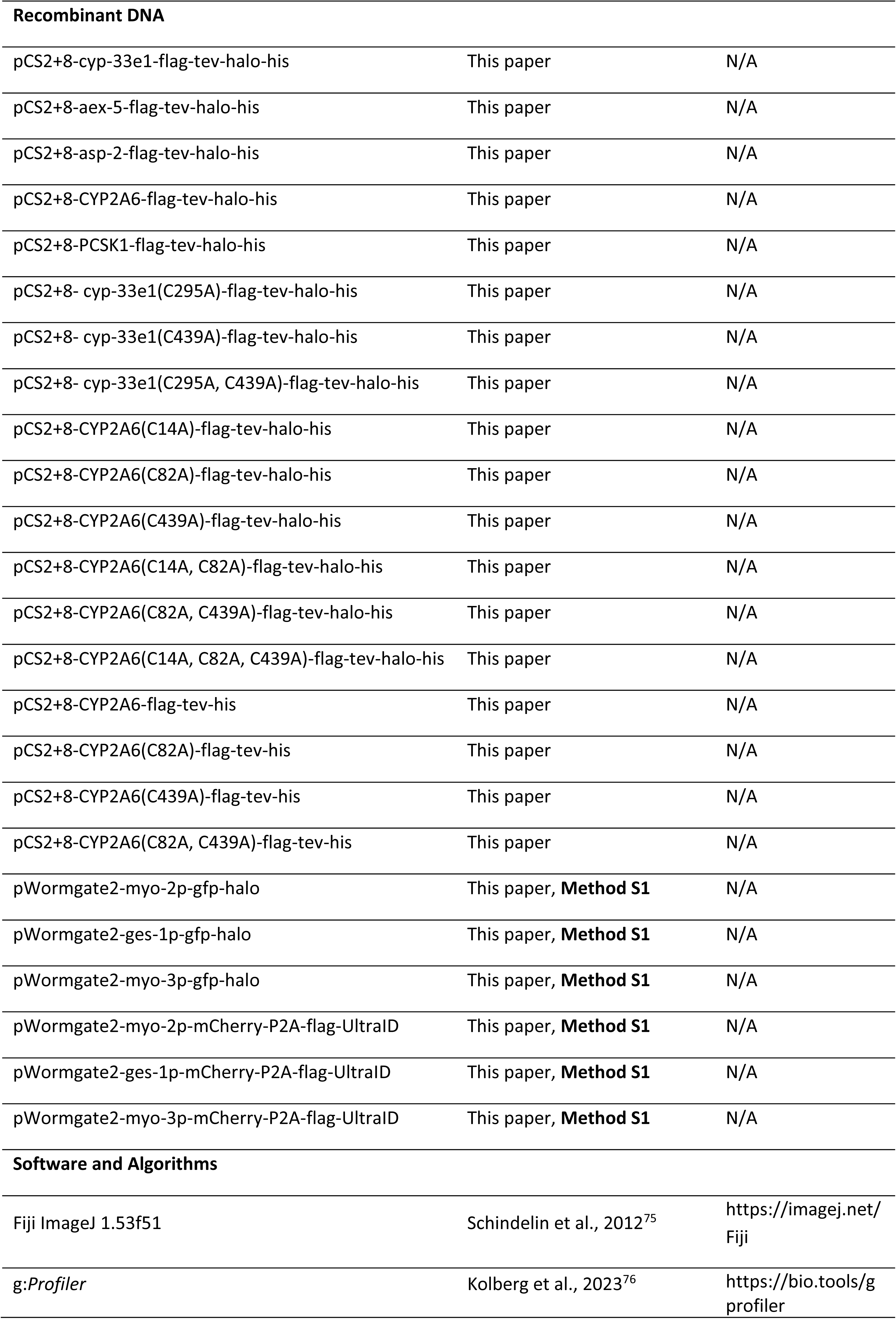
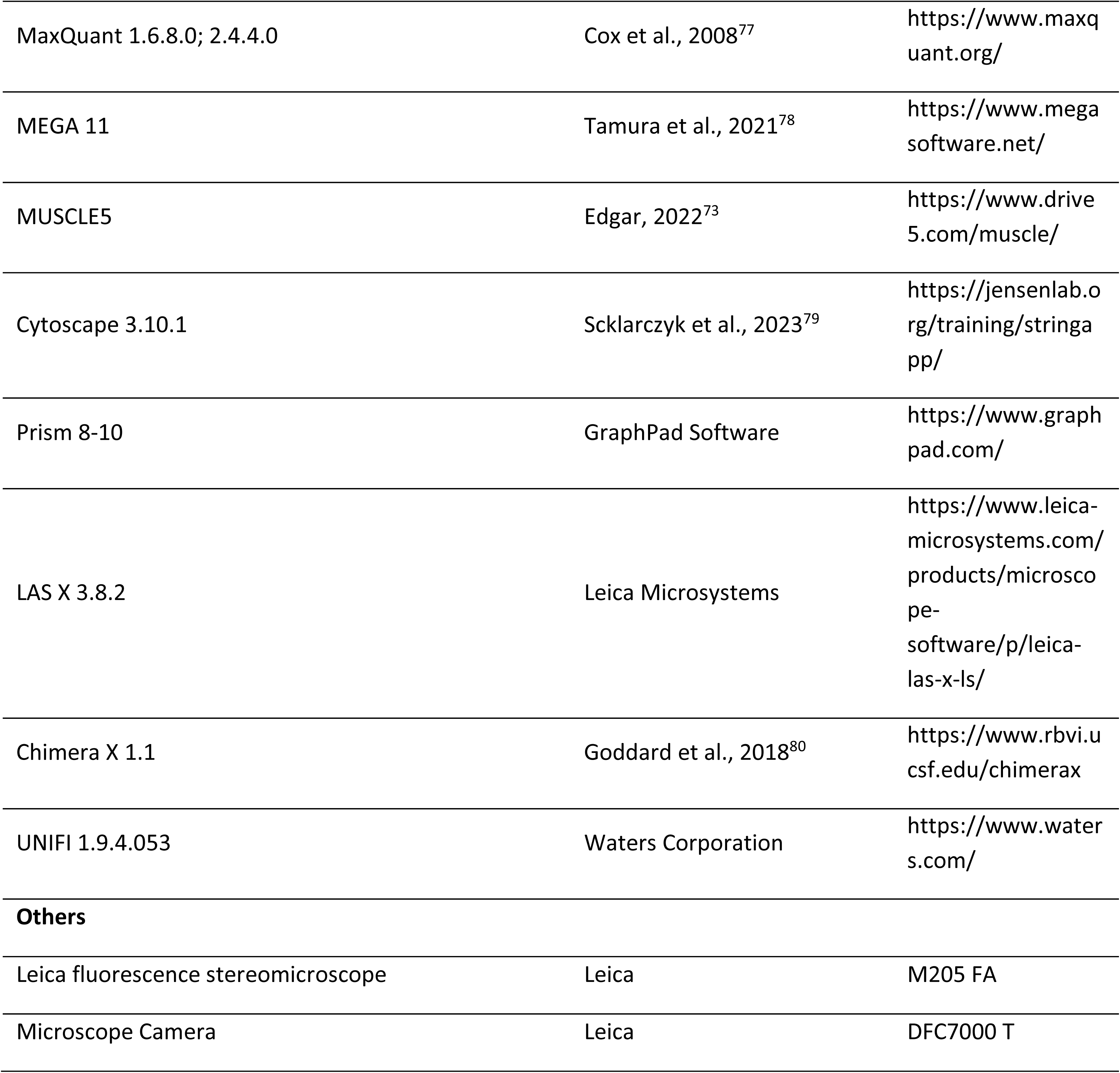

### RESOURCES AVAILABILITY

#### Lead contact

All unique materials will be made available upon reasonable request to the lead contact, Yimon Aye (, valid until 09/30/24; or)

#### Materials availability

The extrachromosomal and integrant C. elegans strains generated in this study (Key Resources Table) are being distributed to Caenorhabditis Genetics Center (CGC). All recombinant DNA generated in this study (Key Resources Table) are being distributed to addgene. Further information and requests for resources and reagents should be directed to the Lead Contact.

#### Data and Code availability

Proteomics data that support the findings of this study: PRIDE ProteomeXchange identifier number is **PXD027730** (TMT organ-specific electrophile-responder analysis in OS-Localis-REX strains, **Figure 1C** and **1D**; **Data S1** and **S2**), **PXD052114** (TMT organ-specific abundance analysis in OS-Ultra-ID strains, **Figure 2B-2D**; **Data S5**), and **PXD052493** (LFQ whole-proteome analysis in OS-Ultra-ID strains, **Figure S2B**; **Data S4**).

## EXPERIMENTAL MODEL AND SUBJECT DETAILS

### Technical Services and CROs

TMT-multiplex data processing and analysis were performed by Dr. Florence Armand (EPFL SV proteomics core facility) in collaboration with R.H., J.L., and Y.A. *C. elegans* transgenic and knock-in lines were designed by J.L., D.A.U., M.J.C.L., and Y.A., and subsequently generated in collaboration with the SUNY Biotech CRO. Small-molecule MS analysis data acquisition, processing and analysis were performed by Dr. Daniel Ortiz Trujillo (EPFL ISIC mass-spectrometry core facility) in collaboration with Y-Q.G., and J.L.

### Construction of plasmids

Sequences of all primers used for cloning and mutagenesis are listed in Method S1. For protein overexpression in cultured HEK293T cells, all constructs were generated in pCS2+8 vector, using ligase-free Hawaiian cloning. Cyp-33e1, aex-5, and asp-2 sequences were amplified from *C. elegans* cDNA, and cloned into pCS2+8 vector using the appropriate forward and reverse primers with Phusion Hotstart II following the manufacturer’s protocol. Prior to insertion, the empty pCS2+8 plasmid was digested with appropriate restriction enzymes (NEB). The products were digested by DpnI (NEB) for 1 hour before transformation into XL10 Gold Ultracompetent cells (Agilent). The inserts generated were fully validated by Sanger sequencing using Microsynth sequencing service, Lausanne, Switzerland.

For cyp-33e1 and CYP2A6 mutagenesis in pCS2+8 vector, the appropriate forward and reverse primers listed in Method S1 containing desired mutations were designed by Agilent QuikChange tool. Under different annealing temperatures (from 56 °C to 68 °C), pCS2+8 backbone and the primers were incubated in the thermocycler with Phusion Hotstart II, following the manufacturer’s protocol. The products were digested by DpnI (NEB) for 2 hours before transformation into XL10 Gold Ultracompetent cells. The mutation sites were fully validated by Sanger sequencing using Microsynth sequencing service, Lausanne, Switzerland.

#### *C. elegans* maintenance and culture

Previously-reported procedures were followed^81^. Briefly, All *C. elegans* strains were grown at 17 °C for maintenance unless otherwise indicated. All worms were derived from the Bristol N2 *C. elegans* strain. Nematode growth medium (NGM-agarose) contained: 0.79 g Tris base (6.5 mM), 2 g NaCl (34 mM), 3 g tryptone, 17 g agarose, pH = 8.0, per liter of ddH_2_O. Post autoclaving, when the temperature had become 50 -60 °C, 1 mL (5 mg/L) cholesterol in EtOH was added per liter of media to render the final concentration as 5 μg/L. For a 10-cm plate, 15 mL NGM-agarose media was pipetted using sterile procedures. In parallel, to prepare OP50, bacteria were inoculated into 3 mL LB media (containing the following components in final concentrations: 10 g/L NaCl, 10 g/L tryptone, 5 g/L yeast extract). The cultures were incubated overnight at 37 °C with shaking, and 1 mL was to 200 mL flasks with ∼60 mL LB media. Then the flasks were incubated overnight at 37 °C with shaking. 1.5 mL OP50 was dispensed into the middle of solidified 10 cm NGM-agarose plates (described above) at room temperature in sterile biosafety cabinets. When OP50 was dry, the plates were subsequently stored at 4 °C. 20 healthy L4-young adult *C. elegans* were selected using platinum picks and grown on 10 cm NGM-agarose plates for 4-5 days at 20 °C until harvesting, which was performed when the plate had become densely populated but prior to starvation. Unless otherwise indicated, worm ages were not synchronized.

#### *C. elegans* Strains Generation

**(i) Transgenic strains for OS-Localis-REX.** (See **Method S1** for gene sequence and primers used to sequence the injected plasmids. Plasmids and worm strains will be deposited to Addgene and CGC repositories, respectively, post manuscript publication)
**(ii) Transgenic strains for OS-Ultra-ID.** (See **Method S1** for gene sequence and primers used to sequence the injected plasmids. Plasmids and worm strains will be deposited to Addgene and CGC repositories, respectively, post manuscript publication)

For transgenic strain generation, 10 μL corresponding plasmids were injected into the gonads of 30 adult wild-type P_0_ worms. For OS-Localis-REX strains, the plasmids concentration was 100 ng/μL, mixed with 100 ng/μL plasmids (of the same backbone) with mec-7p::mRFP transgene marker. For OS-Ultra-ID strains, the plasmids’ concentrations were: 10 ng/μL for myo-3p strains, 20 ng/μL for ges-1p strain, and 5 ng/μL mixed with empty vector pSL1190 plasmid (95 ng/uL) for myo-2p strain. The F_1_ progeny carrying transgenic arrays were further selected and isolated in individual NGM plates for culturing. After the stable transgenic lines were isolated, the single worm PCR was applied to the extrachromosomal lines containing target plasmid for verification.

**(iii) Knock-in strains using CRISPR-cas9.** (See **Method S1** for gene sequence and primers used in single-worm PCR for mutation-site validations. Plasmids and worm strains will be deposited to Addgene and CGC repositories, respectively, post manuscript publication)

The sgRNAs were designed for both knock-in strains: Sg1: CCGTAAGCAGGCTACAGATAGAC; Sg2: CCGACCGACCTAAACTCAACTAT for PHX8800 (cyp-33e1 syb8800, C295A); Sg1: CCAAAATGGAACTATTCTTGTAT;

Sg2: CCTGAACCATACGAATTCAAACC for PHX8844 (cyp-33e1 syb8844, C439A). A ∼1200 kb sequence near the mutation site was designed as the repair template. The sgRNA plasmids, repair template, Cas9 plasmid and co-injection marker were mixed and injected into the gonads of 30 adult P_0_ worms. The F1 progeny with the precise expected sequence were further selected and isolated in individual NGM plates for culturing, verified by single worm PCR and restriction enzyme cutting. The stable homozygous progeny was isolated and validated by single worm PCR and sequencing.

#### Cell Culture

HEK293T (obtained from ATCC) and were cultured in MEM (Gibco) supplemented with 10% v/v fetal bovine serum (Sigma), penicillin/ streptomycin (Gibco), sodium pyruvate (Gibco), and nonessential amino acids (Gibco) at 37 °C in a humidified atmosphere of 5% CO2. Media were changed every 2–3 days. Mycoplasma test was performed tri-monthly using Venor GeM PCR-based mycoplasma detection kit. The kit was used as stated in the manual and purchased from Sigma.

## METHOD DETAILS

### Validation of antibodies

To validate primary antibodies used for the western blot analyses of the lysates originating from *C. elegans*, Bristol N2 (wild-type) strain, which does not express Halo/GFP, was used. Identical procedures of sample handling and data processing were performed for N2 worms as for transgenic worms. Antibodies used for the western blot analyses of the lysates originating from HEK293T cells (Key resource table) have been previously validated in the literature. Antibodies in the Oxyblot kit were validated by eliminating the carbonyl group derivatization using 2,4-Dinitrophenylhydrazine (DNPH) solution provided by the manufacturer.

### Western blotting

Cells lysates, worm lysates, or appropriate samples as described in 1x Laemmli dye, were loaded on 11% or 15% SDS gel, separated by SDS-PAGE, transferred to PVDF either at 30 V overnight or 80V for 1 hour, followed by 55 V for 2 hours. The membrane was blocked in either 5% skimmed milk or 1% BSA recommended by the manufacturer for 2 hours, and then incubated with the appropriate antibodies (Key resource table). Detection was carried out on the Fusion FX imager (Vilber) using ECL Western Blotting Substrate (Pierce) or Immobilon ECL Ultra Western HRP Substrate (Sigma). Western blot data were quantitated using the Gel Analysis tool in Fiji ImageJ 1.53f51.

### Worm lysis

Worm pellets were removed from -80 °C and thawed on ice. 1x Roche cOmplete^TM^ mini (EDTA free) protease inhibitor (in the final concentration recommended by manufacturer) was dissolved in 4.48 mL 50 mM HEPES (pH = 7.6), to which 50 μL 100 mM TCEP in 50 mM HEPES (pH = 7.0) and 0.5 mL 10% Triton-X100 in 50 mM HEPES (pH = 7.6) were added to generate the lysis buffer, which was freshly prepared just before use and kept chilled on ice. Per each worm pellet, 2-8 x pellet volumes of lysis buffer (400 µL) and 1/4 volume of zirconia beads were applied. (In pulldown experiments, volumes of pellets used where: typically, 60-100 µL). Once the worm pellets were thawed, the lysate buffer and the beads were added and the samples were vortexed for 15 seconds then freeze-thawed using liquid nitrogen (repeated 3 times). Afterwards, the samples were centrifuged at 20,000 x g, at 4 °C for 10 minutes. The supernatant was collected, and the protein concentration (in independent triplicate) was measured by Bradford assay. In each replicate of the Bradford assay, 1 µL lysate was mixed with 1 mL Bradford reagent. Then 200 µL of the mixture was pipetted to 96-well plates. Absorbance at 595 nm was measured using a plate reader and protein concentration (mean value over triplicate measurements) was assessed using BSA as a standard. The lysates were immediately used after Bradford quantitation.

### Optimization of probe concentration for Halo-binding site saturation *in vivo*

The following procedure was used to identify the concentration of Ht-PreHNE(alkyne) that can saturate Halo protein binding site in vivo. All assays were performed under dim-red light. All plates of (densely packed, but not starved) worms were harvested with 3 mL of S buffer [6.45 mM K_2_HPO_4_, 43.6 mM KH_2_PO_4_, 100 mM NaCl, filter-sterilized (0.22 μm)], and the samples were pipetted into 15 mL falcon tubes using glass pipettes. [In terms of the number of worms, per each Click reaction condition, 3 x 10 cm plates of confluent worms (densely packed, but not starved) were typically used]. To wash away OP50, the tubes were centrifuged at 3000 x g for 2 min. The supernatant was removed, and the pellets were washed with 10 mL S buffer until the supernatant was clear. Worm pellets were resuspended in 1 mL S buffer, pooled then split equally into 15 ml falcon tubes. To prevent starvation, the following procedure was performed: 3 mL of a saturated OP50 culture was centrifuged and the resulting OP50 pellet was resuspended in 2.5 mL of S buffer. This resulting mixture was added to each of the falcon tubes above containing pelleted worms to fully resuspend the worms. In parallel, Ht-PreHNE(alkyne) was diluted in S buffer to prepare a 10x stock (1 mM). 500 µL, 250 µL, 120 µL, and 60 µL of this 1 mM stock solution were added to the samples above to render a gradient of final concentrations. The final volume of each condition was made up to 5.0 mL by adding S buffer [respectively, to give final concentrations of Ht-PreHNE(alkyne), as follows: 100 µM, 50 µM, 25 µM, and 12 µM]. After 6 hours of incubation involving end-to-end rotation under dim-red light at room temperature, the samples were washed twice (each time with 10 mL S buffer) and a third time with 50 mM HEPES (pH = 7.6), each wash/rotation lasting 30 minutes. Between each wash, the worms were pelleted by centrifugation at 3000 x g for 2 min. Subsequently, the samples were transferred into 1.5 mL tubes and pelleted, then flash-frozen in liquid nitrogen. The resulting pellets could be kept at -80 °C, for up to 1 week. Worm pellets were lysed (see section ‘worm lysis’), and lysates were made to a concentration of 1 mg/mL and treated with 10 µM of Halo-TMR and incubated at 37 °C for 30 min. 10 µL 4X Laemmli buffer with 6% BME was added to quench the reaction. The tubes were either stored at –80 °C, protected from light, or directly analyzed by SDS-PAGE. For each Click reaction condition, 3 x 10 cm plates of (densely packed, but not starved) worms were used.

### Precision localized electrophile delivery in tissue-specific Halo transgenic *C. elegans*

For technical controls deployed (i.e., independent experiments that offer meaningful comparisons in terms of effects of probe, UV light, etc., in non-targeted backgrounds), please refer to main text discussion and individual figure legends. All plates of (densely packed, but not starved) worms were washed with 3 mL of S buffer, and the samples were pipetted into 15 mL falcon tubes using glass pipettes. The samples were centrifuged at 3000 x g for 2 min. The supernatant was removed, and the pellets were washed with 10 mL S buffer until the supernatant was clear. Worm pellets were resuspended in 1 mL S buffer, pooled then split equally into 15 ml falcon tubes. 6 mL of a saturated OP50 culture was centrifuged and resuspended in 5 mL of S buffer per tube. To prevent starvation, the OP50 in S buffer was added to each tube, as described in the section above. Afterwards, Ht-PreHNE(alkyne) was diluted in S buffer to make a 10x stock (120 µM), and 1000 µL of this solution was added to the samples. The final volume for each condition was made up to 10.0 mL by adding S buffer. [The final concentration of Ht-PreHNE(alkyne) was 12 µM]. After 6 hours of incubation involving end-to-end rotation under dim-red light at room temperature, the samples were washed twice (each time with 10 mL S buffer) and a third time with 50 mM HEPES (pH = 7.6). Each wash/incubation lasted 30 minutes. Between each wash, the worms were pelleted by centrifugation at 3000 x g for 2 min. Subsequently, the samples were transferred to a 6-well plate. The experimental group and the control group (as described in **Figure 1b** and elsewhere) were placed under a UV lamp (366 nm, 5 mW/cm^2^) for 5 min. Then the samples were transferred to 1.5 mL tubes and pelleted, and flash-frozen in liquid nitrogen. The resulting pellets could be kept at -80 °C for up to 1 week.

### Time-dependent photo-uncaging assay *in vivo*

Following Ht-PreHNE(alkyne) treatment and washing steps as described in the section above, worm samples were placed under a UV lamp (366 nm, 5 mW/cm^2^) for 0, 1, 3, 5, and 15 min. The protein concentration of lysates (assessed by Bradford assay described elsewhere) was normalized to 1 mg / mL in 200 µL. 250 µL 5x stock Click mixture for 1250 µL total reaction volume was prepared by mixing 62.6 µL 20% SDS, 62.6 µL t-BuOH, 12.5 µL 100 mM CuSO_4_, 12.5 µL Cu(II)TBTA (10 mM in 55% DMSO), 50 µL 50 mM HEPES (pH = 7.6), 25 µL 500 µM Cy5-azide. 25 µL of 100 mM TCEP stock solution [freshly-prepared in Argon-saturated 500 mM HEPES (pH = 7.0)] was added just before use. 50 µL of 5x Click mixture was added to 200 µL of 1 mg/mL lysates. The final concentration of Click reaction components were 1% SDS, 5% t-BuOH, 1 mM CuSO_4_, 0.1 mM Cu (II)TBTA, 2 mM TCEP, 10 µM Cy5-azide in 250 µL samples. The samples were incubated at 37 °C for 30 minutes following vortexing (a few seconds). 30 µL samples were pipetted to new 1.5 mL tubes. 10 µL 4x Laemmli dye with 6% BME was added to each tube to quench the reaction. The samples were subjected to in-gel fluorescence analysis.

### Biotin-Click pulldown for enrichment of tissue-specific electrophile-sensor proteins

For each pulldown assay for target enrichment, 6 x 10 cm plates were used per condition. (Typically, the volume of worm pellets obtained from 6 x 10 cm plates is 100 µL). Final lysate concentration and volume of the lysates were standardized to 2.5 mg/mL and 1000 µL, respectively. 100 µL bed volume of pre-rinsed streptavidin beads were prepared with 1 mL ddH_2_O then 2x 1 mL 0.5% LDS in 50 mM HEPES (pH = 7.6) by end-to-end rotation for 5 min. Between each step, the beads were pelleted by 500 x g centrifuge for 1 min and the supernatant was removed. Subsequently, the samples were added to the beads. After 2 h incubation on a rotator at room temperature, tubes were centrifuged at 2000 x g for 1 min and lysate was removed from the beads.

1.25mL of 5x Click mixture for 6.25 mL total volume was prepared by mixing 313 µL of assay buffer containing in final concentration: 20% SDS, 313 µL t-BuOH, 62.5 µL 100 mM CuSO_4_, 62.5 µL 10 mM Cu(II)TBTA in 55% DMSO, 361.5 µL 50 mM HEPES (pH = 7.6), 12.5 µL 100 mM biotin azide. 125 µL of 100 mM TCEP [freshly prepared in argon-saturated 500 mM HEPES (pH = 7.0)] was added just before use. 250 µL of 5x Click mixture was added to 1000 µL 2.5 mg/mL lysates. The final concentration of Click components in the assay mixture were: 1% SDS, 5% t-BuOH, 1 mM CuSO_4_, 0.1 mM Cu(II)TBTA, 2 mM TCEP, and 200 μM biotin azide. The mixture was incubated at 37 °C for 30 min after vortexing (a few seconds). Subsequently, 4x volume of pre-chilled (−20 °C) pure ethanol was added to each lysate, followed by storage at -80 °C, from overnight to 3 days.

After precipitation, the samples were centrifuged at 20,000 x g for 30 min at 4 °C. The supernatant was removed, and the pellets were washed twice with 1 mL ice-cold 70% ethanol by vortexing for 1 minute, then one wash with acetone. Between each wash, the proteins were pelleted by 20,000 x g, at 4 °C for 10 min. Following the last wash, acetone was removed and evaporated. Next, 80 μL of resuspension buffer: 8% LDS and 1 mM EDTA in 50 mM HEPES (pH = 7.6) was added to each pellet. The pellets were resolubilized by vortexing and sonicating using Bioruptor® Pico sonication device (cat. # B01060010) over 10 minutes. This process was repeated until the pellets were fully solubilized. Samples were diluted with 50 mM HEPES (pH = 7.6) to a final concentration of 0.5% LDS and centrifuged at 20,000 x g for 10 minutes. 30 μL sample was removed and treated with 10 μL 4 x Laemmli dye to be used as an input sample.

The remaining samples were moved to a fresh tube with 50 µL bed-volume of streptavidin beads that had been washed with 250 µL of ddH2O (washed twice) followed by 250 µL of buffer [50 mM Hepes (pH 7.6) (final wash). After 2 h incubation on a rotator at room temperature, the samples were spun down at 2000 x g for 1 min and a small sample (30 μL) of supernatant was removed and treated with 10 μL 4 x Laemmli dye to be used as a flow-through sample. The beads were washed 3 times with 1 mL 0.5% LDS for 30 minutes at room temperature then centrifuged at 500 x g for 5 min, room temperature. After all of the wash buffer, including residual amount, was carefully removed, 30 μL of 2 x Laemmli dye with 6% BME was added to beads. Then the samples were boiled for 10 min at 98 °C, then centrifuged at 20,000 x g for 5 min. The supernatant was removed as the elution sample.

### Optimization of external biotin treatment time and dose for OS-Ultra-ID biotinylation activity saturation in live *C. elegans*

For all OS-Ultra-ID-assocaited experiments, 20 healthy transgenic adults (P_0_) were picked onto standard 10 cm NGM plates. After the plates were full of the F_2_ progeny (not starved) at the larval stage, all worms were collected and then redistributed to freshly-made MG1655:BioBKan plates as a 1 to 6 ratio of 10 cm plates (i.e., 1 plate to 6 plates). The following procedure was used to identify the external biotin treatment time and dose that can saturate biotinylation activity in vivo. All assays were performed under dim-red light. All plates of (densely packed, but not starved) worms were harvested with 3 mL of S buffer [6.45 mM K_2_HPO_4_, 43.6 mM KH_2_PO_4_, 100 mM NaCl, filter-sterilized (0.22 μm)], and the samples were pipetted into 15 mL falcon tubes using glass pipettes. [In terms of the number of worms, per each condition, one 10 cm plate of confluent worms (densely packed, but not starved) was typically used]. To wash away MG1655:BioBKan, the tubes were centrifuged at 3000 x g for 2 min. The supernatant was removed, and the pellets were washed with 10 mL S buffer until the supernatant was clear. Worm pellets were resuspended in 1 mL S buffer, pooled then split equally into Eppendorf tubes. To prevent starvation, the following procedure was performed: 10 mL of a saturated OP50 culture was centrifuged and the resulting OP50 pellet was resuspended in 2.5 mL of S buffer. 250 µL of this resulting 4x mixture was added to each of the Eppendorf tubes above containing pelleted worms to fully resuspend the worms. In parallel, biotin was diluted/resuspended in S buffer to prepare a 4x stock. This stock solution was added to the samples above to render a gradient of final concentrations as 0, 0.2, 1, 5, and 25 mM. The final volume of each condition was made up to 1.0 mL by adding S buffer. After 0.5, 1, 2, and 4 hours of incubation involving end-to-end rotation under dim-red light at room temperature, the samples were washed twice (each time with 1 mL S buffer) and a third time with 50 mM HEPES (pH = 7.6), each wash/rotation lasting 30 minutes. Between each wash, the worms were pelleted by centrifugation at 3000 x g for 2 min. Subsequently, the samples were flash-frozen in liquid nitrogen. The resulting pellets could be kept at -80 °C, for up to 1 week. Worm pellets were lysed (see section ‘worm lysis’), and lysates were made to a concentration of 1.25 mg/mL. 4X Laemmli buffer with 6% BME was added before being loaded onto gel and analyzed by streptavidin blot.

### Biotinylated protein enrichment for OS-Ultra-ID strains

For all OS-Ultra-ID-associated experiments, 20 healthy transgenic adults (P_0_) were picked onto normal 10 cm NGM plates. After the plates were full of the F_2_ progeny (not starved) at the larval stage, all worms were collected and then redistributed to freshly-made MG1655:BioBKan plates as a 1 to 6 ratio of 10 cm plates (1 plate to 6 plates). (*Note*: Biotin deficiency limits the number of F_2_ progeny that could be raised from 20 P_0_ adults in MG1655:BioBKan plates). In terms of the amount of worms, per each mass-spec sample, 6 x 10 cm plates of confluent worms (densely packed, but not starved) were typically used. The external biotin treatment concentration was 5 mM and the time was 4 hours. The other procedural steps were similar to those described above. After the worm lysis and protein concentration normalization, 30 μL sample was removed and treated with 10 μL 4 x Laemmli dye to be used as an input sample for LFQ analysis. The input samples were collected for the streptavidin blot analysis. The remaining lysates were moved to a fresh tube with 50 µL (bed-volume) of streptavidin beads that had been washed with 250 µL of ddH_2_O (washed twice) followed by 250 µL of buffer [50 mM Hepes (pH 7.6) (final wash). After 2 h incubation on a rotator at room temperature, the samples were spun down at 2000 x g for 1 min and a small sample (30 μL) of supernatant was removed and treated with 10 μL 4 x Laemmli dye to be used as a flow-through sample. The beads were washed 3 times with 1 mL 0.5% LDS for 30 minutes at room temperature then centrifuged at 500 x g for 5 min, at room temperature. After all of the wash buffer, including residual amount, was carefully removed, 30 μL of 2 x Laemmli dye with 6% BME was added to beads. Then the samples were boiled for 10 min at 98 °C, then centrifuged at 20,000 x g for 5 min. The supernatant was removed as the elution sample for TMT-11plex analysis.

### Mass spectrometry sample preparation in OS-Localis-REX and OS-Ultra-ID experiments

For OS-Localis-REX samples, following the biotin-click pulldown, all elution samples were loaded into separate wells of a home-made 11% SDS-PAGE gel and were run until the dye-front had penetrated 1-2 cm into the gel to desalt the samples. For OS-UltraID samples, the samples were loaded into Novex™ Tris-Glycine mini protein gels. The gel was cut into small pieces which were washed twice in 50% ethanol and 50 mM ammonium bicarbonate (AB) for 20 min and dried by vacuum centrifugation. Sample reduction was performed with 10 mM DTT for 1 h at 56°C. A washing-drying step as described above was repeated prior to alkylation with 55 mM Iodoacetamide for 45 min at 37°C in the dark. Samples were washed and dried again and digested overnight at 37°C using Mass Spectrometry grade Typsin at a concentration of 12.5 ng/µL in 50 mM AB and 10 mM CaCl_2_. Resulting peptides were extracted in 70% ethanol, 5% formic acid twice for 20 min with permanent shaking. Samples were further dried by vacuum centrifugation and stored at -20 °C. Peptides were desalted on C18 StageTips and dried by vacuum centrifugation. For TMT labeling, peptides were first reconstituted in 8 μL HEPES 100 mM (pH 8.5) and 3 μL of TMT solution (20 µg/μL in pure acetonitrile) was then added. Labeling was performed at room temperature for 1.5 h and reactions were quenched with hydroxylamine to a final concentration of 0.4% (v/v) for 15 min. TMT-labeled samples were then pooled at a 1:1 ratio across all samples. The combined sample was vacuum centrifuged near dryness and subjected to fractionation using the Pierce High pH Reversed-Phase Peptide Fractionation Kit following the manufacturer’s instructions.

### Mass spectrometry data analysis in OS-Localis-REX and OS-Ultra-ID experiments

The individual samples for LFQ analysis of the whole-proteome abundance in individual OS-Ultra-ID worms, or the resulting 12 fractions for TMT analysis of tissue-specific abundance identification within individual OS-Ultra-ID worms, the resulting fractions were resuspended in 2% acetonitrile, 0.1% FA and nano-flow separations were performed on a Dionex Ultimate 3000 RSLC nano UPLC system (Thermo Fischer Scientific) on-line connected with a Qexactive HF Orbitrap mass spectrometer (Thermo Fischer Scientific). A capillary precolumn (Acclaim Pepmap C18, 3 μm-100Å, 2 cm x 75μm ID) was used for sample trapping and cleaning. A 50cm long capillary column (75 μm ID; in-house packed using ReproSil-Pur C18-AQ 1.9 μm silica beads) was then used for analytical separations at 250 nl/min over 150 min biphasic gradients. Acquisitions were performed through Data-Dependent Acquisition (DDA). First MS scans were acquired with a resolution (ion mass/mass peak width) of 120,000 (at 200 m/z) and the 15 most intense parent ions were then selected and fragmented by High energy Collision Dissociation (HCD) with a Normalized Collision Energy (NCE) of 32% using an isolation window of 0.7 m/z. Fragmented ions were acquired with a resolution of 30,000 (at 200 m/z) and selected ions were then excluded for the following 40 s.

For OS-Localis-REX, the raw data were processed using MaxQuant 1.6.8.0 against a concatenated database consisting of the UniProt *C. elegans* complete reference proteome (26991 entries_LM190929) and a HaloTag-GFP protein. For OS-Ultra-ID, the raw data were processed using MaxQuant 2.4.4.0 against a concatenated database consisting of the UniProt *C. elegans* complete reference proteome (26695 entries_LM240314) and a mcherryUltraID protein. Carbamidomethylation was set as fixed modification, whereas oxidation (M), phosphorylation (S, T, Y), acetylation (Protein N-term) and glutamine to pyroglutamate were considered as variable modifications. A maximum of two missed cleavages were allowed and “Match between runs” option was enabled. For LFQ data from whole-proteome abundance analysis of individual OS-Ultra-ID strains, proteins with at least 2 valid values in at least one group were kept for the following analysis: imputation of missing values is performed following a normal distribution with a width = 0.3 and a down-shift = 1.8. For TMT data analysis of tissue-specific abundance from individual OS-Ultra-ID strains, a minimum of 2 unique peptides (in OS-Localis-REX) or 1 unique peptide (in OS-UltraID) were required for protein identification and the false discovery rate (FDR) cutoff was set to 0.01 for both peptides and proteins. Hits were further selected using criteria described in main text. See also **Figure S1I** and **S2F**.

### Age synchronization of *C. elegans*

For synchronization, 3 x 10 cm plates of indicated worm strains were used. All plates were harvested using 5 mL of M9 media [5.8 g Na_2_HPO_4_·7H_2_O (22 mM); 3.0 g KH_2_PO_4_ (22 mM); 5.0 g NaCl (86 mM); 0.25g MgSO_4_·7H_2_O (1.0 mM) for 1 L, that had been filter-sterilized (0.22 μm) before use]. The samples were pipetted into a 15 mL falcon tube using glass pipettes. The tube was centrifuged at 2000 x g for 1 min to pellet the worms. After aspirating the supernatant media, 15 mL of fresh 20% alkaline hypochlorite solution (3.0 mL bleach, 3.75 mL 1 M NaOH and 8.25 mL ddH_2_O) was added to the tube. Once most of the worm bodies had dissolved (typically after 5 minutes of end-to-end rotation), the M9 media was added to fill the tube and then the tube was centrifuged at 4000 x g for 1 min to pellet the eggs. After aspirating the hypochlorite solution, the pellet was washed with 3 x 10 mL M9 media. Subsequently, the pellet was resuspended with 7 mL M9 media and incubated overnight at room temperature with gentle rocking to hatch the eggs. The L1 worms were distributed onto 3 x 10 cm plates the next day.

### RNA interference assay in *C. elegans*

The previously reported procedures were followed. Briefly, the normal nematode growth medium (NGM-agarose) was prepared as described elsewhere. After autoclaving and cooling to 50 - 60 °C, 1 mL of 5 mg/L cholesterol in EtOH, 1 mL of 1 M IPTG, and antibiotics (Ampicillin, 50 μg/mL) were added. For 10 cm plates, 15 mL NGM-agarose media was pipetted using sterile procedures. The respective Ahringer’s dsRNA-encoding plasmids or L4440 empty vector were transformed into HT115 competent cells and were grown in LB/Ampicillin (50 μg/mL) at 37 °C overnight. The concentrated bacterial culture was then seeded onto the NGM plates with IPTG and left to dry at room temperature for overnight induction of dsRNA expression. Synchronized L1 worms were raised on these plates (for targeted knockdown) at 22 °C for 40-48 hours.

### RNA isolation and reverse transcription

Synchronized late L4 worms were collected from RNAi plates and washed 3 times with M9 media to remove bacteria. The worms were then resuspended in 1 mL TRIzol™ reagent and homogenized. Samples were used immediately for RNA isolation or stored at -80 °C until further use. To isolate the RNA, 0.2 mL of chloroform was added to 1 mL TRIzol™ reagent then mixed by shaking. After incubation for 2-3 min, the samples were centrifuged at 12000 x g at 4 °C for 15 min. The aqueous phase containing RNA was transformed into a new Eppendorf tube. 0.5 mL of isopropanol was added to precipitate the RNA. The aqueous phase was incubated with isopropanol for 10 min at 4 °C and then centrifuged at 12000 x g at 4 °C for 10 min. After discarding the supernatant, the RNA pellet was washed with 1 mL 75% (v/v) ethanol. The samples were vortexed briefly before centrifugation at 7500 x g at 4 °C for 5 min. The resulting pellet was resuspended with 20-50 µL Rnase-free water. Extracted RNA was quality-controlled by Agarose gel electrophoresis analysis and concentration was determined by A_260nm_ using a BioTek Cytation3 microplate reader with a Take3 accessory. cDNA was synthesized with reverse transcription kits using RNaseOUT™ Recombinant Ribonuclease Inhibitor and Superscript™ III Reverse Transcriptase.

### Quantitative real-time PCR (qRT-PCR)

For each sample (and associated technical and biological replicates as described in figure legends), PCR was performed using iQ SYBR Green Supermix (Bio-Rad) and primers specific to the gene of interest following the manufacturer’s protocol. Amplicons were chosen that were around 200 bp in length and had no predicted off-target binding predicted by NCBI Primer BLAST. For genes with multiple splice variants, primers were chosen that amplified conserved sequences across all splice variants. Primers were validated using standard curves generated by amplification of serially-diluted cDNA; primers with a standard curve slope between –0.8 and 1.2 and R^2^≥0.97 were considered acceptable. Single PCR products were confirmed by melting analysis following the PCR protocol. Quantitative PCR data was carried out using QuantStudio™ 7 Pro. Samples with a threshold cycle >35 or without a single, correct melting point were not included in data analysis. Normalization was carried out using a single housekeeping gene (β-actin) as indicated in each dataset and the ΔΔC_t_ method. See **Method S1** for primer sequences.

### Scoring of fertilized eggs laid by *C. elegans*

Plates for RNAi-targeted knockdown (6 cm, 8 mL of NGM-media) were prepared as described elsewhere. For each phenotypic assay, 12 L4-young adult worms were picked from 10 cm plates (subjected to RNAi), at the end of 40 hours post plating of L1. The worms were transferred individually to 6 cm plates (subjected to RNAi). On the morning and the evening of Day 1 and 2 post plating of L1, the worms were transferred to fresh 6 cm plates (subjected to RNAi), and further on each subsequent evening, until no new eggs were observed. The number of eggs was counted at specific time intervals - 3 times on Days 1 and 2; 2 times on Days 3 and 4, once on subsequent days, until no new eggs were observed. All counted eggs were validated as healthy embryos by assessment of hatching.

### *C. elegans* viability analysis

RNAi-fed worms (from 6 cm plate size, prepared using 8 mL of NGM-media) were prepared as described above. For each phenotypic assay, 10 L4-young adult worms were picked from 10 cm RNAi plates 40 hours post plating of L1, and were transferred individually to 6 cm RNAi plates. The worms were monitored for their survival. The worms were transferred to fresh 6 cm RNAi plates in the morning and the evening of Day 1 and Day 2; and in the evening of subsequent days, until no new eggs were observed. A worm was considered dead if no response was observed after 3 gentle touches on the head with a platinum pick and it displayed no pharyngeal pumping.

### *C. elegans* body bend assay

RNAi-fed worms (from 6 cm plate size, prepared using 8 mL of NGM-media) were prepared as described elsewhere. For each phenotypic assay, 10 L4-young adult worms were picked from 10 cm RNAi plates 40 hours post plating of L1, and were transferred individually to 6 cm RNAi plates. The number of bends at the pharynx was counted for 2 minutes for every worm, every alternate day until death. Data were recorded only when the worm was away from the edges of the bacterial food lawn. The worms were transferred to fresh 6 cm RNAi plates in the morning and the evening of Day 1 and Day 2; only in the evening of subsequent days until no new eggs were observed.

### *C. elegans* egg-laying assay

RNAi-fed worms (from 6 cm plate size, prepared using 8 mL of NGM-media) were prepared as described elsewhere. For each phenotypic assay, 3 x 10 cm plates were prepared and synchronized L1 worms were raised on them for 40 hours at 22 °C. The worms were then collected and washed 3 times with M9 media to remove bacteria. The worms were then resuspended in 0.5 mL M9 media and transferred to 1.5 mL tubes. HNE-alkyne-functionalized (in vehicle DMSO) master stock solution was prepared in M9 media and added to each tube such that final concentration was 2 mM in final volume of 0.7 mL. For each HNE-alkyne-treatment condition, corresponding final volume of DMSO vehicle was used as a control. After one hour of incubation involving end-to-end rotation, samples were washed 3 times with 0.5 mL M9 media and resuspended in 0.2 mL M9 media. These worm suspensions were seeded onto freshly-prepared RNAi plates. The worms were allowed to revive for 4 hours at 20 °C. 12 L4-young adult worms were then picked from these RNAi plates and transferred individually to 6 cm RNAi plates. On the next evening (Day 2), the worms were transferred to fresh 6 cm RNAi plates, and subsequent transfers were performed in the morning and evening of Days 2-3; and on the evening of subsequent days until no new eggs were observed. The number of eggs was counted at specific time intervals - 3 times on Days 1-2; 2 times on Days 3-4; once on subsequent days until no new eggs were observed.

### *C. elegans* food race assay

Food race plates were prepared at least two days prior to experiment, which were NGM plates with 50 µL of OP50 seeded as a dot 1 cm away from the edge of the plate on the equator, as established and described. Large population of worms was bleached-synchronized and recovered on either NGM plates or RNAi plates (10 cm plates, as described elsewhere) at 22°C for 40 hours. The worms were then collected and treated by different doses of HNE(alkyne) or by the equivalent volume of its vehicle solvent in M9 buffer for 1 hour with end-to-end rotation, washed 3 times by M9, and recovered on NGM plates for 4 hours. The worms were then collected again and transferred to a marked position also 1 cm away from the edge of the plate on the opposite side of the OP-50 lawn on the equator drop-wise. The timer starts when the M9 has evaporated, and the number of worms that have reached the lawn was recorded at 30, 60, 90, and 120 min, and the next day, matched against the total number of alive worms.

### *C. elegans* defecation assay

RNAi-fed worms (from 10 cm plate size, prepared using 15 mL of NGM media) were prepared as described elsewhere. The food lawn was normalized to a 1.2 OD value before being seeded on the NGM media specially for this assay. For each phenotypic assay group, two 10 cm plates were prepared and synchronized L1 worms from the same pool were raised on these plates for 48 hours at 22 °C. The half worms were for imaging without the treatment. For each condition, 12 individual worms are recorded individually for 10 x defecation cycles by Leica M205 FA microscope. The other half worms were then collected and resuspended in 0.5 mL M9 media and transferred to 1.5 mL tubes. HNE-alkyne-functionalized (in vehicle DMSO) master stock solution was prepared in M9 media and added to each tube such that the final concentration was 2 mM in a final volume of 0.75 mL. For each HNE-alkyne-treatment condition, the corresponding final volume of DMSO vehicle was used as a control. After one hour of incubation involving end-to-end rotation, samples were washed 3 times with 0.5 mL M9 media and resuspended in 0.2 mL M9 media. These worm suspensions were seeded onto freshly prepared 10 cm RNAi plates. The worms were cultured at 20 °C afterward. 16 hours later, 20 worms are recorded individually for 6 x defecation cycles for each condition. The defecation cycle was determined by intervals between two pBoc behaviors.

### *C. elegans* male induction and crossing

20 x L4-young adult hermaphrodites were picked onto 2-3 x 10 cm plates, which were incubated for 6 hours at 31 °C afterward. The plates were transferred to 20 °C until the healthy F1 worms grew after the induction. Then all the males were picked to another 6 cm plate with healthy L4-young hermaphrodites at a 3:1 to 4:1 ratio. The males from the next generation were used for crossing. The hermaphrodites from another strain were picked together with these males to 35 mm plates for crossing at a similar male-to-hermaphrodite ratio. After 3 days, heterozygous F1 worms from crossing were picked to fresh 35 mm plates individually to ensure their self-crossing. Subsequently, 16-20 x hermaphrodite F2 worms were picked to 35 mm plates separately. Once enough F3 progenies from F2 self-crossing were observed on the plates, the F2 mothers were picked out and checked by single worm PCR. 10 x individual F3 worms were picked and checked by single worm PCR to ensure the homozygosity from the F2 mother containing the target transgene or mutation. Sometimes more self-crossing steps were required depending on the F3 homozygosity. Only the homozygous worms would be used for subsequent experiments. All crossing and culture temperatures are 20 °C.

### Single worm PCR

Single adult worms were picked into a 10 μL worm digestion buffer in a PCR tube (1X HF buffer, 0.5 mg/mL proteinase K). The tubes were spun down at 14000 g for 15 seconds before digestion. The worms were digested by flash-freezing in liquid nitrogen for 10 min followed by a 90-minute incubation at 65 °C and a 15-minute protease K inactivation at 95 °C. Then the mixture was used directly for PCR as the template. For the PCR reaction, 2 μL DMSO, 0.2 μM corresponding primer pair, 0.2 mM dNTPs mixture, and 0.04U/ μL Phusion polymerase were added to a final volume at 50 μL 1X HF buffer. The PCR cycle was set to 98 °C for 50 seconds, and 30 times 98 °C for 10s; 53 °C for 30 seconds; 72 °C for 45 seconds. After the PCR reaction, the product was checked with 1% agarose in NEB buffer. DNA electrophoresis gel and Sanger sequencing after cleaning with the PCR clean-up kit, if necessary.

### Lipid droplet imaging

RNAi-fed worms containing dhs-3p::dhs-3::gfp transgene (from 10 cm plate size, prepared using 15 mL of NGM-media) were prepared as described elsewhere. For each phenotypic assay group, one 10 cm plate was prepared and synchronized L1 worms from the same pool were raised on these plates for 48 hours at 22 °C. The worms were then collected and resuspended in 0.5 mL M9 media and transferred to 1.5 mL tubes. HNE-alkyne-functionalized or its reduced version DHN; oxidized version HNA (in vehicle DMSO) master stock solution was prepared in M9 media and added to each tube such that the final concentration was 2 mM in a final volume of 0.75 mL. For each small-molecule-treatment condition, the corresponding final volume of DMSO vehicle was used as a control. After one hour of incubation involving end-to-end rotation, samples were washed 3 times with 0.5 mL M9 media and resuspended in 0.2 mL M9 media. These worm suspensions were seeded onto freshly prepared 10 cm RNAi plates. The worms were cultured at 20 °C afterward. 16 hours later, individual worms were picked and imaged under both stereo objective and 5x HR objective (Leica FluoCombi III) with a Leica M205 FA microscope. For Oil Red O staining experiments, instead of imaging the worms directly, the worms were collected by M9 to 1.5 mL tubes and washed 3 times. Worm pellets without supernatant were fixed by adding 500 μL 60% isopropanol. 500 μL freshly 0.45-μm filtered 0.3% Oil Red O in 60% isopropanol (by mixing 0.5% Oil Red O in 100% isopropanol and ddH_2_O the day before) was added to the worms to enable the staining. After 18 hours of end-to-end rotation at 25 °C in a wet chamber (web paper towels and aluminum foil covered), the supernatant was removed. The remaining worm pellets were washed with 500 μL 60% isopropanol twice and then added with 250 μL 0.01% Triton X-100 in S-basal buffer. Stained worms can be imaged or stored at 4 °C for a month. The GFP signals were quantified by selecting the intestinal region and analyzed in Fiji ImageJ 1.53f51 under the 5x HR objective mode. The Oil Red O images were taken under the RGB mode with the 5x HR objective; the signals were quantified by selecting the body region (without the pharynx) in the blue channel after inverting the images in Fiji ImageJ 1.53f51.

### *C. elegans* immunofluorescent imaging following bulk HNE treatment

Healthy, age-synchronized L4-young adult worms single adult worms were collected, suspended in 0.5 mL M9 media, and transferred to 1.5 mL tubes. HNE-alkyne-functionalized (in vehicle DMSO) master stock solution was prepared in M9 media and added to each tube such that the final concentration was 2 mM in a final volume of 0.75 mL. For each HNE-alkyne-treatment condition, the corresponding final volume of DMSO vehicle was used as a control. After one hour of incubation involving end-to-end rotation, samples were washed 3 times with 0.5 mL M9 media and fixed in 1 mL 4% paraformaldehyde overnight with gentle end-to-end rotation at 4 °C. After pelleting at 2000x g and removal of paraformaldehyde, the worms were incubated in methanol at -20 °C for 24 h. Subsequently, the worms were washed for 30 min each, twice with PBS including 0.015% Tween-20, and twice with 50 mM HEPES buffer including 150 mM NaCl, pH = 7.6. The signal of alkyne functionalized HNE was visualized by click reaction: the worms were incubated in the freshly made cocktail click mixture, with the final concentration containing 50 mM HEPES (pH = 7.6), 150 mM NaCl, t-BuOH (5%), CuSO4 (1 mM), Cu(II)TBTA (0.1 mM), TCEP [2 mM; made as a 100 mM stock in 500 mM HEPES (pH 7.6)] and Cy5-azide (10 μM). The TCEP was added at the end of the mixture preparation. The worms were incubated at room temperature in the Eppendorf thermomixer at 300 g for 1 hour.

Afterward, the samples were washed 3 times in PBS including 0.015% Tween-20, and then incubated overnight with gentle end-to-end rotation at 4 °C. For the subsequent incubation with antibodies, the worms need to be washed twice in 1 mL PDT buffer (PBS, 0.1% Tween-20, 1% DMSO) for 30 min each at room temperature and blocked in 1 mL blocking solution (PBS, 0.05% Tween-20, 10% v/v FBS, 2% w/v BSA) for 1 hour at room temperature. After the blocking, the worms were incubated with goat anti-GFP (B2) (Abcam, ab6673) solution (1:500 in blocking solution) at 4 °C overnight. Similarly, the worms were washed twice in 1 mL PDT buffer (PBS, 0.1% Tween-20, 1% DMSO) for 30 min each at room temperature and blocked in 1 mL blocking solution (PBS, 0.05% Tween-20, 10% v/v FBS, 2% w/v BSA) for 1 hour at room temperature. Then Donkey pAb to Goat IgG AlexaFluor 488-conjugated (Abcam, ab150133) solution (1:500 in blocking solution) was added to the samples and incubated for 1.5 hours. After 3 times washing in 1 mL PDT buffer, 30 min each at room temperature, the worms were imaged under both stereo objective and 5x HR objective (Leica FluoCombi III) with a Leica M205 FA microscope. The fluorescent signals were quantified by selecting the intestinal region and analyzed in Fiji ImageJ 1.53f51 under the 5x HR objective mode.

### Cell growth, transfection, and targetable electrophile sensing test (T-REX)

HKE293T and Hela cells were maintained in 1 x MEM + Glutamax media supplemented with 10% FBS, 1× NEAA, 1× sodium pyruvate, and 1× Pen-Strep in 60 mm growth factor treated plates. When the cell density was ∼70%, cells were transfected with specific halo-POI plasmid using TransIT-2020 transfection reagent per the manufacturer’s recommendation. All subsequent steps were done under a dim-red light. 36 hours after transfection, the cells were treated with 15 μM Ht-PreHNE(alkyne) in serum-free media and incubated for 2 h, followed by gentle rinsing with serum-free media three times every 30 min over the next 1.5 h. The preheated UV light (10-min on before using, 365 nm, ∼ 5 mW/cm^2^) was placed 2 cm above the sample for 5 min for photo-uncaging alkyne functionalized HNE. The lids of cultural dishes were removed during the irradiation. Afterward, cells were harvested by TrypLE express enzyme into 1.5 mL Eppendorf tubes, washed two times of ice-cold DPBS, once of 50 mM HEPES (pH = 7.6), and flash-frozen in liquid nitrogen. Cell pellets were lysed in 400 µL freshly prepared lysis buffer containing 50 mM HEPES (pH 7.6), 150 mM NaCl, 1% Triton-X100, 1× Roche cOmplete mini, EDTA-free protease inhibitor mixture, and 0.3 mM TCEP by rapid freeze-thaw and 15-seconds vortex for three times. The cell debris was removed by centrifugation at 20000 g for 30 min at 4 °C. Protein concentrations in cell lysate were determined to be 1.25 mg/mL using Bradford assay. Subsequent biotin-click pulldown steps are similar to methods in worms as described before.

### Enzymatic activity test of CYP2A6 with HNE inhibition

Vivid™ CYP450 Screening Kit (PV6140) from ThermoFisher Scientific was used to test the enzymatic activity of CYP2A6. From 0.25 μM to 32 μM, a series of double concentrations of 3-cyano coumarin (substrate) was added to an opaque 96-well plate to determine the *K*_m_. The enzyme concentration was 10 nM, with 30 μM NADPH in 100 μL reaction volume. The fluorescent signals at 460 nm were recorded every minute, over the course of 1 hour, using Cytation-5 multimode plate reader. The enzymatic reaction velocity was calculated by linear fitting of signals observed within the initial 20-minute period, the data from which were used for fitting to Michaelis-Menten kinetics and calculating the *K*_m_. For HNE-inhibition assays, CYP2A6 was incubated with different concentrations of HNE or EtOH (vehicle) for 20 min, prior to addition of 30 μM NADPH and 100 μM 3-cyano coumarin, rendering final concentrations of CYP2A6 to be 10 nM, and HNE as indicated in corresponding figure, during the measurement of remaining enzyme activity. The fluorescent signals at 460 nm were recorded every minute, over the course of 1 hour, using Cytation-5 multimode plate reader. Enzymatic reaction velocity for subsequent fitting and analysis was calculated based on the data observed during the initial 30-minute period. See main text discussion and corresponding figure legends for the equations used.

### Overexpression and enzymatic activity test of microsomal CYP2A6

HEK293 cells were transfected in 20*10^6^cells /mL in RPMI1640 media (Thermo Fischer) with 0.1% pluronicF68 (Applicam). 15 μg sterile plasmids containing CYP2A6 mutants were added to 10 mL media for transfection. After mixing by shaking, 30 μg Pei-Max (Polyscience, Chemie Brunschwig) were added to the cells, mixed, and cultured at 37 °C with stirring for 1.5 hours. The cells were diluted to a concentration of 1*10^6^cells/mL in Hycell TransFxH Media (Cytiva). After a 3-day culture with stirring at 37°C, cells were harvested by centrifugation. Cell pellets can either be kept at – 80 °C or lysed in 400 µL (For 5 mL media) 50 mM potassium phosphate buffer (pH =8.0), by rapid freeze-thaw and 15-second vortex three times. The supernatant was removed after centrifugation at 12000 g for 5 min at 4 °C. Protein concentration (c, mg/mL) normalization was based on the Bradford assay for the supernatant in each sample. The remaining cell debris was resuspended in 50 mM potassium phosphate buffer (pH =8.0) again, by matching the buffer volume to the corresponding protein concentrations in supernatant (volume of resuspending buffer = 40*c µL). The subsequent enzymatic activity test steps were similar to the commercial CYP450 Screening Kit as described elsewhere, by substituting the 10 nM enzyme to 50 µL CYP2A6 microsomes. For the activity test of different mutants, the 3-cyano coumarin deployed was 100 μM. For the HNE-inhibition assays, CYP2A6 microsomes were incubated with 400 μM HNE or EtOH (vehicle) for 20 min, prior to addition of 30 μM NADPH and 10 μM 3-cyano coumarin as the final concentration. CYP2A6 amount was determined by western blot analysis.

### *In vitro* HNEylation of CYP2A6 for validation of HNEylation using Cy5 in-gel fluorescence analysis

Recombinant CYP2A6 (TP322995) from Origene was used to test the in vitro HNEylation of CYP2A6. Before the reaction, 90 μM HNE-alkyne and 1.8 μM CYP2A6 were preheated in a 37 °C metal bath. 15 μL CYP2A6 stock (final concentration: 1.5 μM) and 3 μL HNE-alkyne stock (final concentration: 15 μM) were mixed and incubated at 37 °C to start the reaction. 2 μL solution was pipetted to 28 μL ice-cold 50 mM HEPES with 1x Click mixture, at 0s, 15s, 30s, 60s, 90s, 120s, 180s, 240s after mixing. The final concentration for the click reaction contained 1% SDS, 50 mM HEPES (pH = 7.6), t-BuOH (5%), CuSO_4_ (1 mM), Cu(II)TBTA (0.1 mM), TCEP [2 mM; made as a 100 mM stock in 500 mM HEPES (pH 7.6)] and Cy5-azide (10 μM). The mixture of the HNEylated CYP2A6 and click reagents was incubated at 37 °C for 30 min, followed by adding 12.5 μL 4x loading dye with 6% BME. The cy5 signals were read out by in-gel fluorescence and the protein amounts were read out by colloidal Coomassie blue staining. Gels were quantified in Fiji ImageJ 1.53f51 and the data relevant to time-dependent HNE modification was fitted to Y=0.020533 + (1-0.020533)*(1-exp(-k_obs_’*x)) in GraphPad Prism. To enable comparison with the previously-reported electrophile-sensing rates of kinetically-privileged HNE-sensor proteins, The adjusted second-order observed rate constant of *k*_obs_= (3.7 ± 0.8) x 10^8^ M^-2^·s^-1^ was derived by considering the concentration of the protein and HNE-alkyne as described in the main text and corresponding figure legends.

### Sample preparation for HNEylation site identification by mass spectrometry

1 μM of recombinant CYP2A6 (Origene), approximately 2.83 μg in 45 μL, was prewarmed at 37°C on a shaker in a red light room for 5 min. 0.8 μM or 10 μM final concentration of in-house-synthesized HNE(alkyne) in DMSO was added to the protein (5 μL) for the final volume of 50 μL, and incubated for 5 min in a red light room. Samples were then resuspended in 4 M urea, diluted by 5 volumes of with 100 mM Tris-HCl pH 8, and digested overnight at 37°C by 1/20 w/w enzyme-to-protein ratio of mass spectrometry-grade trypsin. The reaction was stopped by the addition of formic acid and peptides were dried by vacuum centrifugation. Peptides were resuspended in 0.1% TFA solution and were desalted in StageTips using 4 C18 disks (Empore 3M) based on the standard protocol^82^. Desalted peptides were dried down by vacuum centrifugation.

### Mass spectrometry (MS) analysis for HNEylation site identification

The tryptic peptides were resuspended in 10 μL of 0.1% FA and 2% acetonitrile (Biosolve) for nano-LC-ESI-MS analysis, which was carried out using an Exploris 480 Orbitrap Mass Spectrometer (Thermo Fischer Scientific) coupled with a Dionex UltiMate 3000 RSLCTM (Thermo, Sunnyvale, CA). 2 μL were injected onto a capillary precolumn (Acclaim Pepmap C18, 3 μm-100Å, 2 cm x 75μm ID) for sample trapping and cleaning. A 50 cm long capillary column (75 μm ID; in-house packed using ReproSil-Pur C18-AQ 1.9 μm silica beads; Dr. Maisch) was then used for analytical separations at 250 nL/min over 90 min biphasic gradients. Acquisitions were performed through Top Speed Data-Dependent Acquisition mode using a cycle time of 2 seconds. First MS scans were acquired with a resolution of 120,000 (at 200 m/z) and the most intense parent ions were selected and fragmented by High energy Collision Dissociation (HCD) with a Normalized Collision Energy (NCE) of 30% using an isolation window of 2 m/z. Fragmented ions were acquired with a resolution of 30,000 (at 200m/z) and selected ions were then excluded for the following 20 s. All data were acquired under Xcalibur 3.0 operation software (Thermo-Fisher Scientific).

### Data analysis for digest-MS-based HNEylation site identification

All MS and MS/MS raw spectra were processed by Proteome Discoverer v2.5 (Thermo Fisher Scientific). Database search was performed against the *H. Sapiens* Uniprot database (UP000005640, revised on 2023 Oct 19) with 82,685 entries and 20,596 genes. Parent mass tolerance was set at 10.0 ppm and fragment mass tolerance was set to 0.05 Da, with search type set to monoisotopic and maximum number of missed cleavages of 2. Variable modifications were set with N-terminal acetylation, carbamidomethylation, and pyro-Q to E. In addition to these, variable modifications of HNE(alkyne) (C_9_H_12_O_2_, Mmi 152.0837), reduced HNE(alkyne) (C_9_H_14_O_2_, Mmi 154.0994), dehydrated HNE(alkyne) (C_9_H_10_O, Mmi 134.0732), and reduced and dehydrated HNE(alkyne) (C_9_H_12_O, Mmi 136.0888) for Cys, His, and Lys were added. Peptide identifications were accepted if they could be established at greater than 84.0% probability to achieve an FDR less than 1.0% by the Percolator posterior error probability calculation. Protein identifications were accepted if they could be established at greater than 99.0% probability to achieve an FDR less than 1.0% and contained at least 2 identified peptides. Protein probabilities were assigned by the Protein Prophet algorithm. Proteins that contained similar peptides and could not be differentiated based on MS/MS analysis alone were grouped to satisfy the principles of parsimony. Scaffold (Scaffold v5.3.1, Proteome Software Inc., Portland, OR) was used to validate MS/MS-based peptide and protein identifications.

### *In vitro* reaction of CYP2A6-mediated HNE(alkyne) turnover, and preparation for product identification by mass spectrometry

The procedure was modified from previous published methods, using reagents from CYP2A6 Vivid Blue Screening Kit (Cat #PV6140). Reaction volume was 20 μL per tube, each containing 10 μL 1 μM CYP2A6 Baculosomes (final 0.5 μM) and 10 μL master mix of 1X Vivid P450 Buffer (final 0.5X), 3.0 mM NADPH (final 1.5 mM), 10.0, 3.0, and 1.0 mM of HNE (final 5.0, 1.5, and 0.5 mM, diluted in ddH_2_O from 10 mM stock in vehicle EtOH), and matching volumes of EtOH. The two mixtures were mixed together and placed at 37°C for 30 min and flash frozen. All samples were thawed together on room temperature shaker with the addition of 1.8 μL 100% TFA to precipitate protein. The thawed solution was centrifuged at 20,000g for 5 min at 4°C, and 1.8 μL of 9.56 M NaOH was added to the collected supernatant. The solution was incubated at 37°C for 30 min on shaker with 3.9 μL fresh 7X mM GSH such that final HNE:GSH ratio of ∼1:2 based on the HNE concentration. 10 μL of the sample was diluted in 90 μL MeOH and placed on ice until mass spectrometry analysis immediately after.

### Mass spectrometry analysis of the product from CYP2A6-assisted HNE(alkyne) turnover

Samples were kept in chilled autosampler at 10°C during analysis. Analysis was performed in negative ion mode on Vion IMS QTOF (Waters) in-line with Acquity UPLC Class I (Waters) Sample Manager, Binary Solvent Manager, and Column Manager, operated under UNIFI software (Waters). 5 μL of each sample was injected onto 40°C heated Acquity Premier BEH 1.7 μm 2.1 x 50 mm C18 Column (Waters, P/N 186009452) at 0.4 μL/min flow rate of 100% Solvent A (0.1% FA) and 0% Solvent B (methanol). Between 0.5 and 3.0 min, Solvent B increases to 100%, and remains until 5.0 min. Between 5.0 and 5.1 min, Solvent B decreases back to 0%, and remains until 6.0 min. The QTOF analyzer was set to sensitivity mode, at 1.4 kV capillary voltage, 120°C and 500°C for source and desolvation temperatures, respectively. Scan range was set between 50-1200 *m/z* at 1 s scan time, lock mass correction every 5 min, and Intelligent Data Capture enabled. MS2 was enabled with collision energy of 4.0 eV at low energy, and between 20.0-40.0 eV for high energy ramping.

Raw data processing was performed in UNIFI via Full Application Processing with the following compounds added for identification: GSH (C_10_H_17_N_3_O_6_S, 307.0838 Da), HNA (C_9_H_16_O_3_, 172.1099 Da), GS-HNE (C_19_H_33_N_3_O_8_S, 463.1988 Da), and GS-HNA (C_19_H_33_N_3_O_9_S, 479.1938 Da). Molecules were identified based on matching to the *m/z*, the retention time (RT), and the Collision Cross Section (CCS) values of these molecules, determined through a separate experiment with standards. Target identification has an RT tolerance of 0.1 min, CCS tolerance of 3%, and a mass tolerance of 10 ppm. Peak smoothing was performed with mean smoothing algorithm at 2 half width with 2 iterations. Peak integration was performed with automatic peak width and threshold detection, integration at 0.0% liftoff and 0.5% touchdown. The output of this processing generates a Review where the observed *m/z*, mass error ppm, observed RT, observed CCS in angstroms, response, and adducts information was available. Response values were used as the integrated area value, used for quantification of the amount of GS-HNA and GS-HNE molecules in each sample.

### Synthesis of alkyne functionalized HNA and DHN

The synthesis until HNA-methyl ester was described as published.

For methyl [(4-chlorophenyl)sulfinyl]acetate (**6**), 3.62 g (25 mmol) starting material **3** was added to 40 mL DMF, then 3.08 g (27.5 mmol) K_2_CO_3_ was added. After stirring for 0.5 h at rt., 3.83 g (25 mmol) **4** and 0.96 g KI were added to the system. After stirring overnight at rt., brine was added to the reaction. The product was extracted by EtOAc; washed with brine and cold water 4-5 times. The organic layer was dried with sodium sulfate and concentrated in a vacuum.

5.02 g (23 mmol) starting material **5** was added to 20 mL CHCl_3_, then 5.75 g (25 mmol) mCPBA was added slowly. After stirring overnight, the reaction was quenched by saturated NaHCO_3_. The product was extracted by CH_2_Cl_2_ and washed with NaHCO_3_ 3 times. The organic layer was dried with sodium sulfate and concentrated in a vacuum. The product was purified by flash silica chromatography (80% hexane, 20% EtOAc) and analyzed by ^1^H-NMR spectroscopy.

For non-2-ene-1,4-diol, 220 mg (1.2 mmol) starting material **7** was dissolved in 20 mL CH_2_Cl_2_ and cooled to 0 °C, then 4.8 mL DIBAL-H in hexane (1M solution) was added drop by drop. After stirring for 3 hours, the reaction was quenched by adding cold water and saturated Rochelle salt solution, followed by CH_2_Cl_2_ extraction. The white-solid product was dried and concentrated, purified by flash silica chromatography (70% hexane, 30% EtOAc), and analyzed by ^1^H-NMR spectroscopy (400 MHz, CDCl_3_): δ 7.06 (dd, *J* = 15.6, 4.6 Hz, 1H), 6.08 (dd, *J* = 15.6, 1.7 Hz, 1H), 4.41 (ddt, *J* = 6.5, 4.7, 2.4 Hz, 1H), 2.26 (td, *J* = 6.7, 2.6 Hz, 2H), 1.98 (t, *J* = 2.6 Hz, 1H), 1.84 – 1.57 (m, 4H).

For 4-Hydroxy-2-nonenoic acid, 220 mg (1.2 mmol) starting material **7** was dissolved in 15 mL THF/water at a 1:1 ratio, then 0.48 g LiOH (20 mmol) was added. After stirring for 2 hours, the reaction was washed by ether two times. Subsequently, 2 M HCl was added to the aqueous layer slowly until pH = 2-3. The yellow-oil product was extracted by EtOAc, dried and concentrated, purified by flash silica chromatography (60% hexane, 40% EtOAc), and analyzed by ^1^H NMR (400 MHz, CDCl_3_) spectroscopy: δ 5.84 (dt, *J* = 15.6, 5.1 Hz, 1H), 5.75 (m, 1H), 4.15 (t, *J* = 5.2 Hz, 3H), 2.22 (tt, *J* = 4.3, 2.5 Hz, 2H), 2.06 (s, 2H), 1.96 (t, *J* = 2.7 Hz, 1H), 1.70 – 1.53 (m, 4H).

## QUANTIFICATION AND STATISTICAL ANALYSIS

The quantification methods were explained in the corresponding methods. Samples generated from the same pool of animals (e.g., 6 x 100 mm plates, depending on specific experiments) were considered one biological replicate. Data analysis and statistical treatments are detailed in corresponding figure legends. P-values were calculated either from t-test or adjusted comparisons in ANOVA, explained in the decision tree for the t-test or ANOVA test (**Method S3**). Data were plotted/fit and statistics were generated using GraphPad Prism. Bonferroni correction was applied where applicable and indicated, typically when multiple comparisons elevate the chance of Type 1 errors. When images are shown, they are representative of multiple independent samples. For experiments involving cultured cells, samples generated from individual wells or plates were considered biological replicates. For *C. elegans* experiments, designation of how biological replicates were considered is described in figure legends. In the figure legend for each experiment, how the data are presented in the figure (typically mean ± SEM) is clearly indicated.

