## Supplementary material for "Organ-specific electrophile responsivity mapping in live *C. elegans*": All SI figures

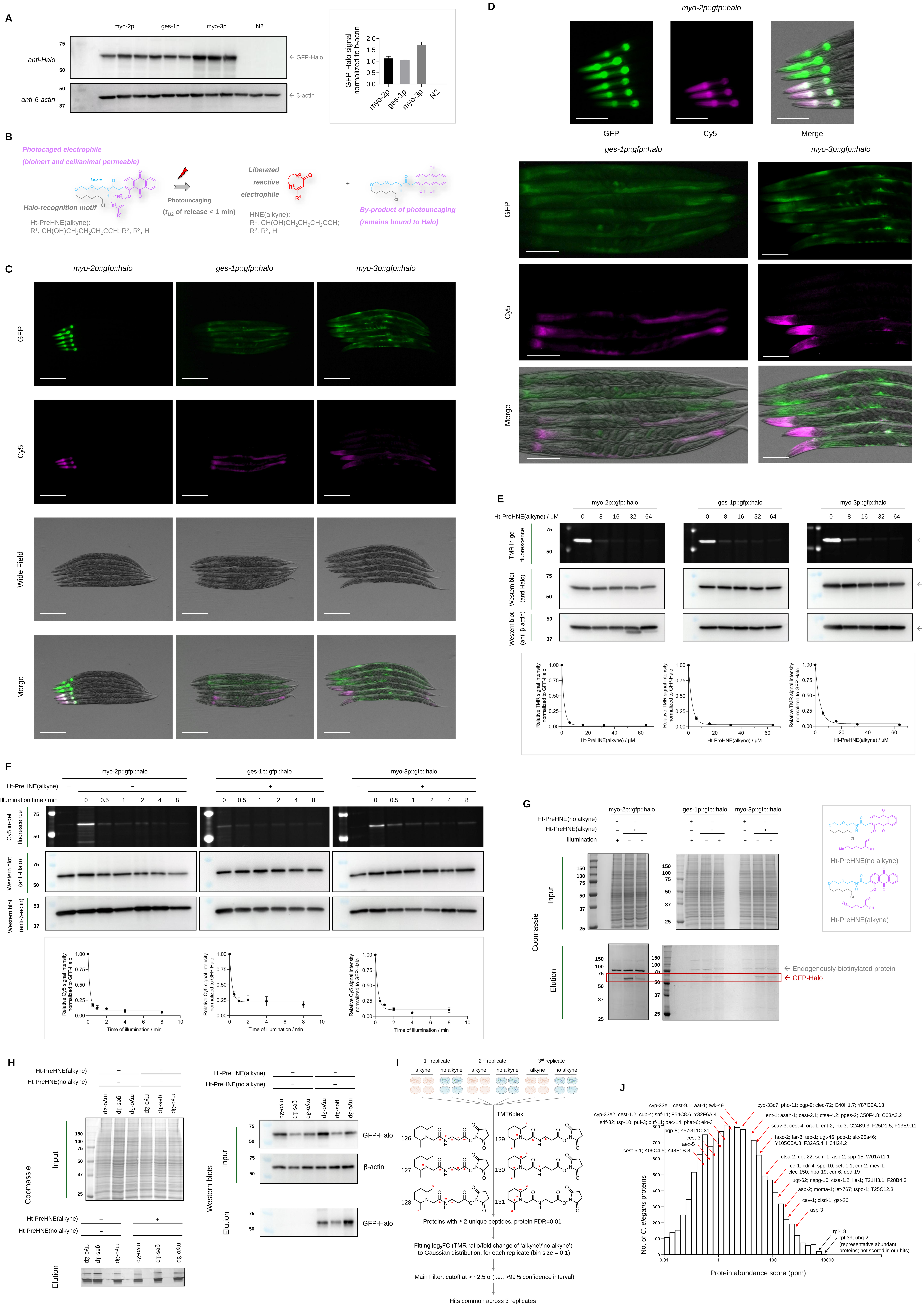

A
myo-2p
ges-1p
myo-3p
N2
anti-Halo
anti-β-actin
← GFP-Halo
← β-actin
GFP-Halo signal normalized to b-actin
B
Photocaged electrophile
(bioinert and cell/animal permeable)
Linker
Halo-recognition motif
Ht-PreHNE(alkyne):
R¹, CH(OH)CH₂CH₂CH₂CCH; R², R³, H
Photouncaging
(t1/2 of release < 1 min)
Liberated reactive electrophile
HNE(alkyne):
R¹, CH(OH)CH₂CH₂CH₂CCH;
R², R³, H
By-product of photouncaging
(remains bound to Halo)
C
myo-2p::gfp::halo
ges-1p::gfp::halo
myo-3p::gfp::halo
GFP
Cy5
Wide Field
Merge
D
myo-2p::gfp::halo
GFP
Cy5
Merge
ges-1p::gfp::halo
myo-3p::gfp::halo
E
myo-2p::gfp::halo
ges-1p::gfp::halo
myo-3p::gfp::halo
Ht-PreHNE(alkyne) / μM
TMR in-gel fluorescence
Western blot (anti-Halo)
Western blot (anti-β-actin)
← GFP-Halo
← β-actin
Relative TMR signal intensity normalized to GFP-Halo
Ht-PreHNE(alkyne) / μM
F
myo-2p::gfp::halo
ges-1p::gfp::halo
myo-3p::gfp::halo
Ht-PreHNE(alkyne)
Illumination time / min
Cy5 in-gel fluorescence
Western blot (anti-Halo)
Western blot (anti-β-actin)
Relative Cy5 signal intensity normalized to GFP-Halo
Time of illumination / min
G
myo-2p::gfp::halo
ges-1p::gfp::halo
myo-3p::gfp::halo
Ht-PreHNE(no alkyne)
Ht-PreHNE(alkyne)
Illumination
Coomassie
Input
Elution
← Endogenously-biotinylated protein
← GFP-Halo
Ht-PreHNE(no alkyne)
Ht-PreHNE(alkyne)
H
Ht-PreHNE(alkyne)
Ht-PreHNE(no alkyne)
myo-2p
ges-1p
myo-3p
Coomassie
Input
Elution
Western blots
GFP-Halo
β-actin
I
1st replicate
2nd replicate
3rd replicate
alkyne
no alkyne
TMT6plex
126
127
128
129
130
131
Proteins with ≥ 2 unique peptides, protein FDR=0.01
Fitting log₂FC (TMR ratio/fold change of 'alkyne'/'no alkyne') to Gaussian distribution, for each replicate (bin size = 0.1)
Main Filter: cutoff at > ~2.5 σ (i.e., >99% confidence interval)
Hits common across 3 replicates
J
No. of C. elegans proteins
Protein abundance score (ppm)
cyp-33e1; cest-9.1; aat-1; twk-49
cyp-33c7; pho-11; pgp-9; clec-72; C40H1.7; Y87G2A.13
cyp-33e2; cest-1.2; cup-4; snf-11; F54C8.6; Y32F6A.4
srlf-32; tsp-10; puf-3; puf-11; oac-14; phat-6; elo-3
ent-1; asah-1; cest-2.1; ctsa-4.2; pges-2; C50F4.8; C03A3.2
pgp-8; Y57G11C.31
scav-3; cest-4; ora-1; ent-2; inx-3; C24B9.3; F25D1.5; F13E9.11
cest-3
aex-5
cest-5.1; K09C4.5; Y48E1B.8
faxc-2; far-8; tep-1; ugt-46; pcp-1; slc-25a46; Y105C5A.8; F32A5.4; H34I24.2
ctsa-2; ugt-22; scm-1; asp-2; spp-15; W01A11.1
fce-1; cdr-4; spp-10; selt-1.1; cdr-2; mev-1; clec-150; hpo-19; cdr-6; dod-19
ugt-62; nspg-10; ctsa-1.2; ile-1; T21H3.1; F28B4.3
asp-2; moma-1; let-767; tspo-1; T25C12.3
cav-1; cisd-1; gst-26
asp-3
rpl-18
rpl-39; ubq-2
(representative abundant proteins; not scored in our hits)

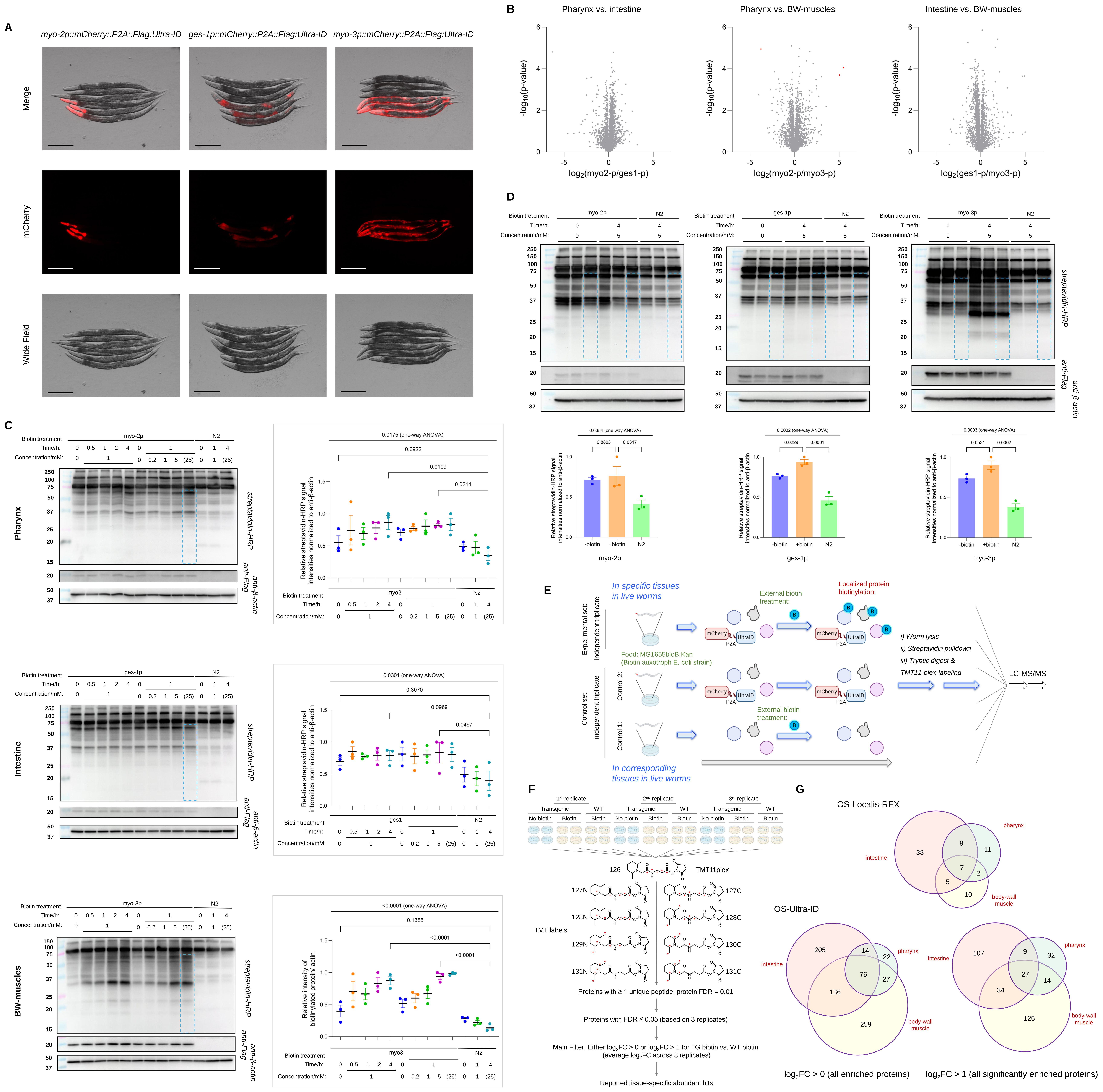

A
myo-2p::mCherry::P2A::Flag:Ultra-ID
ges-1p::mCherry::P2A::Flag:Ultra-ID
myo-3p::mCherry::P2A::Flag:Ultra-ID
Merge
mCherry
Wide Field
B
Pharynx vs. intestine
Pharynx vs. BW-muscles
Intestine vs. BW-muscles
-log10(p-value)
log2(myo2-p/ges1-p)
log2(myo2-p/myo3-p)
log2(ges1-p/myo3-p)
C
Biotin treatment
Time/h:
Concentration/mM:
myo-2p
ges-1p
myo-3p
N2
Pharynx
Intestine
BW-muscles
streptavidin-HRP
anti-Flag
anti-β-actin
0.0175 (one-way ANOVA)
0.6922
0.0109
0.0214
0.0301 (one-way ANOVA)
0.3070
0.0969
0.0497
<0.0001 (one-way ANOVA)
0.1388
<0.0001
<0.0001
Relative streptavidin-HRP signal intensities normalized to anti-β-actin
Relative intensity of biotinylated protein/ actin
myo2
ges1
myo3
D
myo-2p
ges-1p
myo-3p
N2
0.0354 (one-way ANOVA)
0.8803
0.0317
0.0002 (one-way ANOVA)
0.0229
0.0001
0.0003 (one-way ANOVA)
0.0531
0.0002
-biotin
+biotin
N2
E
In specific tissues in live worms
Experimental set: independent triplicate
External biotin treatment:
Localized protein biotinylation:
Food: MG1655bioB:Kan (Biotin auxotroph E. coli strain)
Control set: independent triplicate
Control 2:
Control 1:
mCherry
P2A
UltraID
i) Worm lysis
ii) Streptavidin pulldown
iii) Tryptic digest &
TMT11-plex-labeling
LC-MS/MS
In corresponding tissues in live worms
F
1st replicate
2nd replicate
3rd replicate
Transgenic
WT
No biotin
Biotin
126
TMT11plex
127N
127C
128N
128C
129N
130C
131N
131C
TMT labels:
Proteins with ≥ 1 unique peptide, protein FDR = 0.01
Proteins with FDR ≤ 0.05 (based on 3 replicates)
Main Filter: Either log2FC > 0 or log2FC > 1 for TG biotin vs. WT biotin (average log2FC across 3 replicates)
Reported tissue-specific abundant hits
G
OS-Localis-REX
intestine
pharynx
body-wall muscle
OS-Ultra-ID
log2FC > 0 (all enriched proteins)
log2FC > 1 (all significantly enriched proteins)

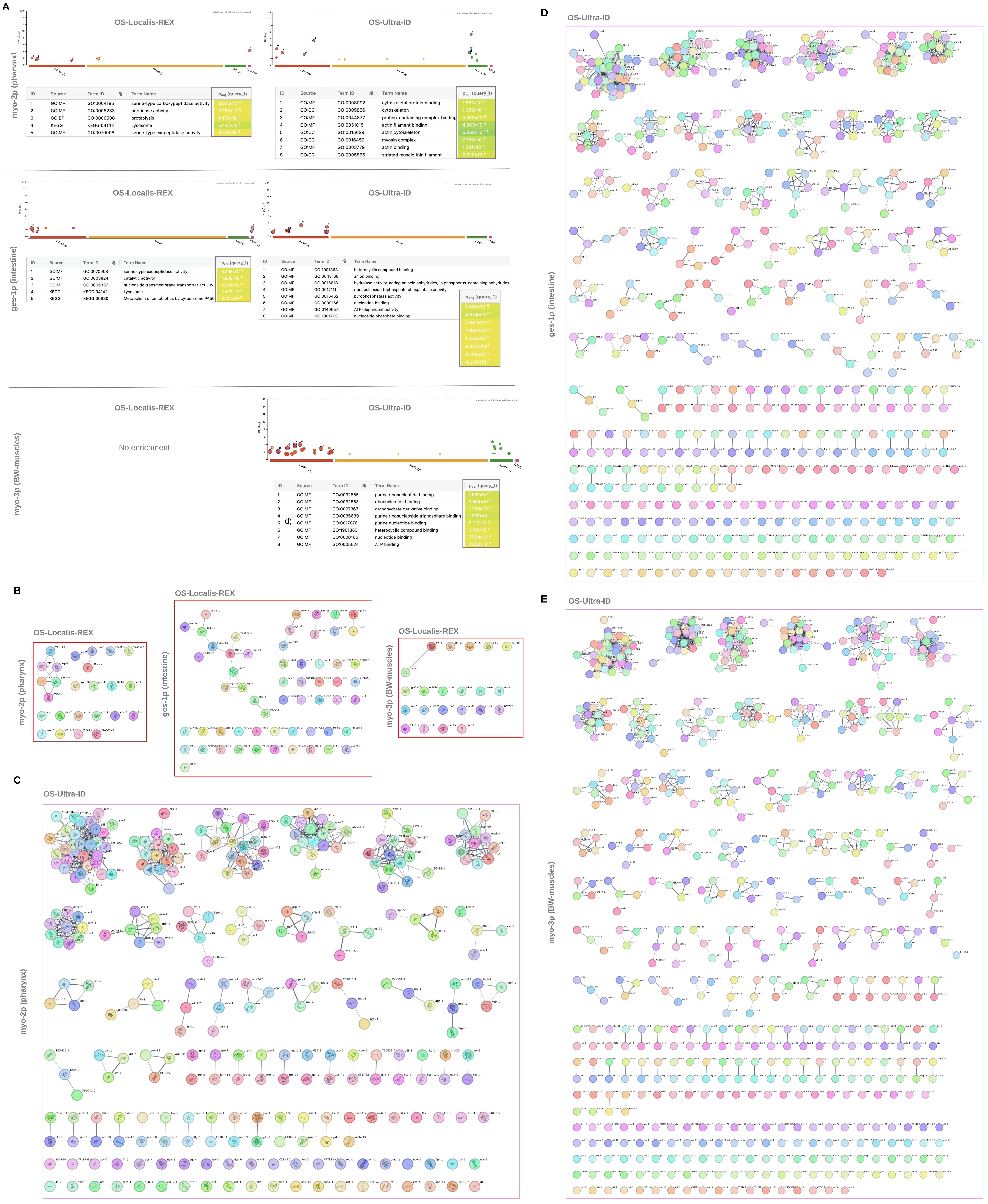


myo-2p (pharynx), OS-Localis-REX

| ID | Source | Term ID | Term Name | $p_{adj}$ (query_1) |
|---|---|---|---|---|
| 1 | GO:MF | GO:0004185 | serine-type carboxypeptidase activity | $9.025\times10^{-3}$ |
| 2 | GO:MF | GO:0008233 | peptidase activity | $2.267\times10^{-2}$ |
| 3 | GO:BP | GO:0006508 | proteolysis | $1.472\times10^{-2}$ |
| 4 | KEGG | KEGG:04142 | Lysosome | $5.447\times10^{-5}$ |
| 5 | GO:MF | GO:0070008 | serine-type exopeptidase activity | $4.715\times10^{-2}$ |

myo-2p (pharynx), OS-Ultra-ID

| ID | Source | Term ID | Term Name | $p_{adj}$ (query_1) |
|---|---|---|---|---|
| 1 | GO:MF | GO:0008092 | cytoskeletal protein binding | $1.662\times10^{-4}$ |
| 2 | GO:CC | GO:0005856 | cytoskeleton | $1.360\times10^{-4}$ |
| 3 | GO:MF | GO:0044877 | protein-containing complex binding | $5.093\times10^{-4}$ |
| 4 | GO:MF | GO:0051015 | actin filament binding | $4.063\times10^{-8}$ |
| 5 | GO:CC | GO:0015629 | actin cytoskeleton | $6.835\times10^{-10}$ |
| 6 | GO:CC | GO:0016459 | myosin complex | $7.350\times10^{-6}$ |
| 7 | GO:MF | GO:0003779 | actin binding | $1.281\times10^{-5}$ |
| 8 | GO:CC | GO:0005865 | striated muscle thin filament | $2.522\times10^{-4}$ |

ges-1p (intestine), OS-Localis-REX

| ID | Source | Term ID | Term Name | $p_{adj}$ (query_1) |
|---|---|---|---|---|
| 1 | GO:MF | GO:0070008 | serine-type exopeptidase activity | $3.338\times10^{-3}$ |
| 2 | GO:MF | GO:0003824 | catalytic activity | $4.908\times10^{-3}$ |
| 3 | GO:MF | GO:0005337 | nucleoside transmembrane transporter activity | $4.004\times10^{-2}$ |
| 4 | KEGG | KEGG:04142 | Lysosome | $2.472\times10^{-3}$ |
| 5 | KEGG | KEGG:00980 | Metabolism of xenobiotics by cytochrome P450 | $3.105\times10^{-2}$ |

ges-1p (intestine), OS-Ultra-ID

| ID | Source | Term ID | Term Name | $p_{adj}$ (query_1) |
|---|---|---|---|---|
| 1 | GO:MF | GO:1901363 | heterocyclic compound binding | $1.749\times10^{-3}$ |
| 2 | GO:MF | GO:0043168 | anion binding | $4.554\times10^{-4}$ |
| 3 | GO:MF | GO:0016818 | hydrolase activity, acting on acid anhydrides, in phosphorus-containing anhydrides | $2.080\times10^{-2}$ |
| 4 | GO:MF | GO:0017111 | ribonucleoside triphosphate phosphatase activity | $2.646\times10^{-2}$ |
| 5 | GO:MF | GO:0016462 | pyrophosphatase activity | $1.753\times10^{-2}$ |
| 6 | GO:MF | GO:0000166 | nucleotide binding | $4.637\times10^{-2}$ |
| 7 | GO:MF | GO:0140657 | ATP-dependent activity | $4.718\times10^{-2}$ |
| 8 | GO:MF | GO:1901265 | nucleoside phosphate binding | $4.637\times10^{-2}$ |

myo-3p (BW-muscles), OS-Ultra-ID

| ID | Source | Term ID | Term Name | $p_{adj}$ (query_1) |
|---|---|---|---|---|
| 1 | GO:MF | GO:0032555 | purine ribonucleotide binding | $1.557\times10^{-4}$ |
| 2 | GO:MF | GO:0032553 | ribonucleotide binding | $2.504\times10^{-4}$ |
| 3 | GO:MF | GO:0097367 | carbohydrate derivative binding | $1.335\times10^{-3}$ |
| 4 | GO:MF | GO:0035639 | purine ribonucleoside triphosphate binding | $1.037\times10^{-4}$ |
| 5 | GO:MF | GO:0017076 | purine nucleotide binding | $8.119\times10^{-4}$ |
| 6 | GO:MF | GO:1901363 | heterocyclic compound binding | $1.142\times10^{-2}$ |
| 7 | GO:MF | GO:0000166 | nucleotide binding | $7.964\times10^{-3}$ |
| 8 | GO:MF | GO:0005524 | ATP binding | $7.303\times10^{-3}$ |

**A**

| | Pharynx | Intestine | Body_wall_muscle | Human ortholog | Human protein function | Human disease / behavioral role |
|---|---|---|---|---|---|---|
| cyp-33e2 | | | | (CYP2J2) | Cytochrome P450 - 2J2; Monooxygenase/oxidoreductase | Hypertension, lipodystrophy |
| ent-1 | | | | SLC29A1 | Equilibrative nucleoside transmembrane transporter | Fetal erythroblastosis, periampullary and gallbladder adenocarcinoma, pancreatic cancer, diabetes |
| asah-1 | | | | ASAH1 | Acid ceramidase | Farber disease, spinal muscular atrophy, keloid disorder |
| cest-2.1 | | | | (CES2) | Cocaine carboxylesterase | Large intestine adenoma |
| cdr-4 | | | | FAXC | Failed axon connections homolog; Stress response to cadmium in *C. elegans* | Deafness |
| cav-1 | | | | CAV1 | Caveolin-1 molecular adaptor | Hypertension, lipodystrophy |
| pho-11 | | | | (ACP2) | Putative acid phosphatase 11 | Acid phosphatase deficiency |
| puf-3 | | | | (PUM1) | Pumilio homolog RNA-binding protein | Growth and body weight regulator |
| ctsa-4.2 | | | | CTSA | Serine carboxypeptidase | Galactosialidosis, leukoencephalopathy |
| asp-2 | | | | (CTSE) | Cathepsin E; aspartic endopeptidase | Gastric adenocarcinomas |
| aex-5 | | | | PCSK1 | Neuroendocrine convertase 1(Proprotein convertase subtilisin/kexin type 5); Serine endopeptidase inhibitor | Neurodegenerative diseases |
| cyp-33e1 | | | | CYP2A6 | Cytochrome P450 – 2A6; Oxidoreductase | Cardiovascular diseases |
| let-767 | | | | HSD17B12 | Very-long-chain 3-oxoacyl-coA reductase; testosterone dehydrogenase | Platelet deficiency, central hypoventilation syndrome |
| mev-1 | | | | SDHC | Succinate dehydrogenase (mitochondrial) | Cowden syndrome, gastrointestinal stromal tumor, paragangliomas |
| cup-4 | | | | (CHRNA9) | Neuronal acetylcholine receptor subunit alpha-9 (ion channel) | Tobacco addition, deafness |

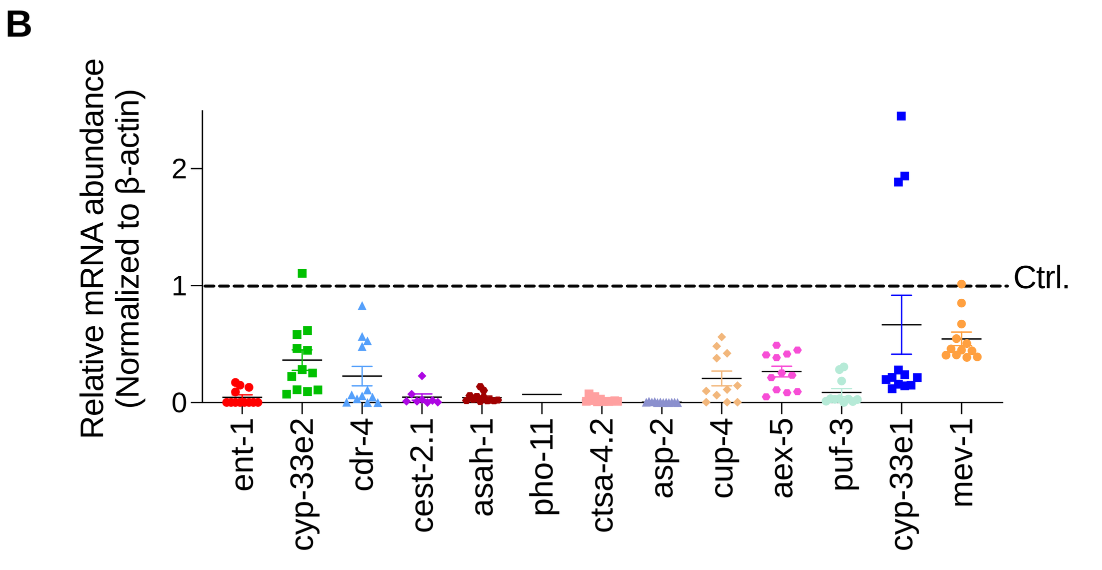


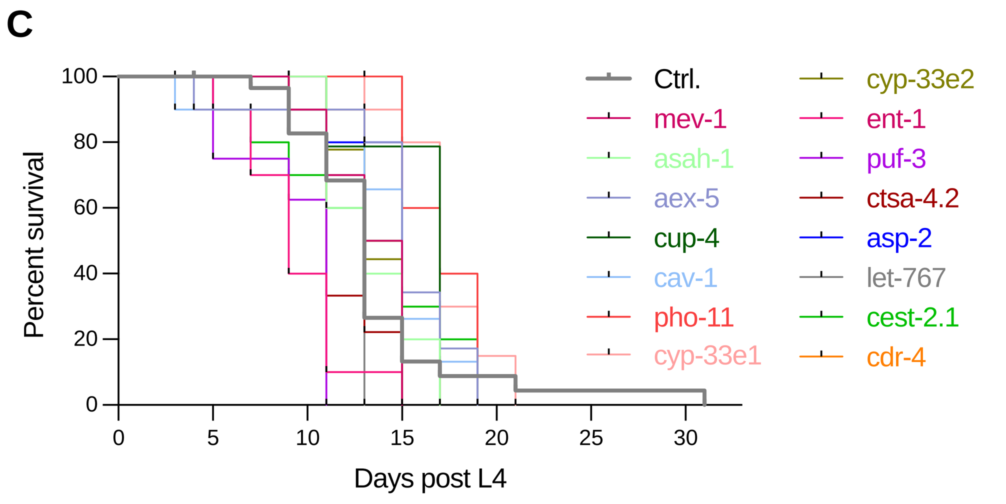


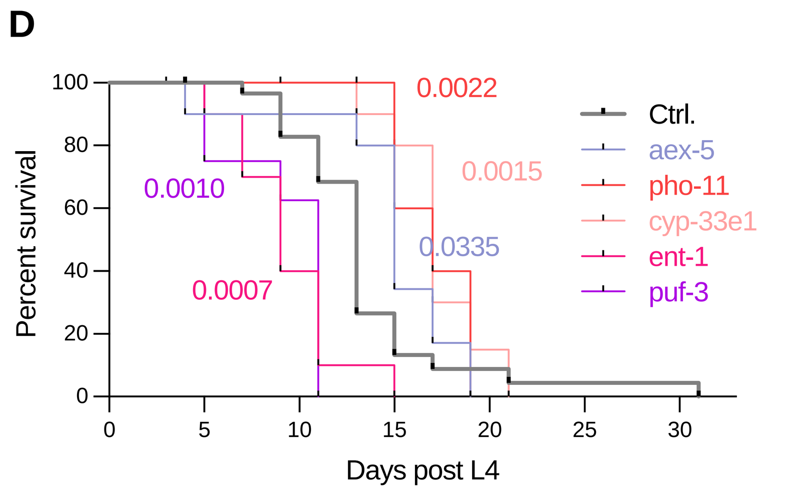


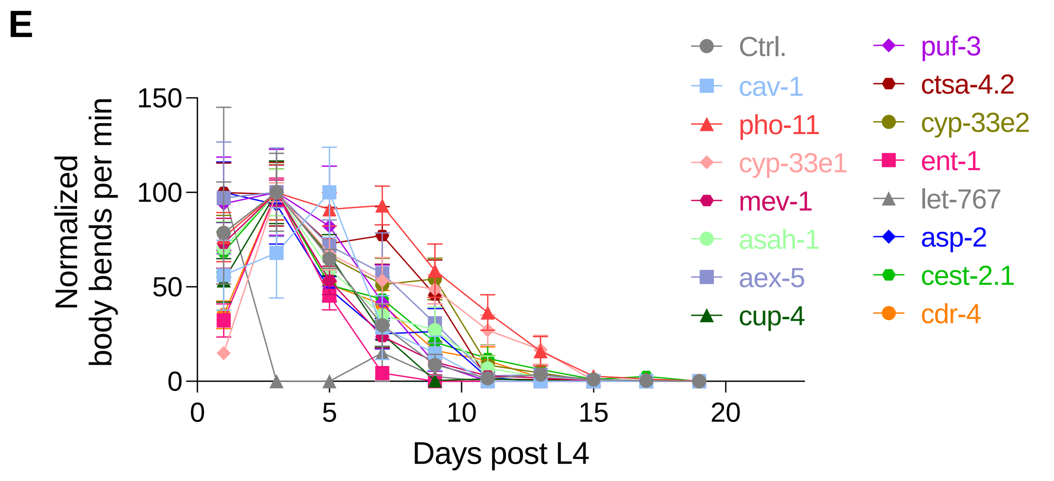


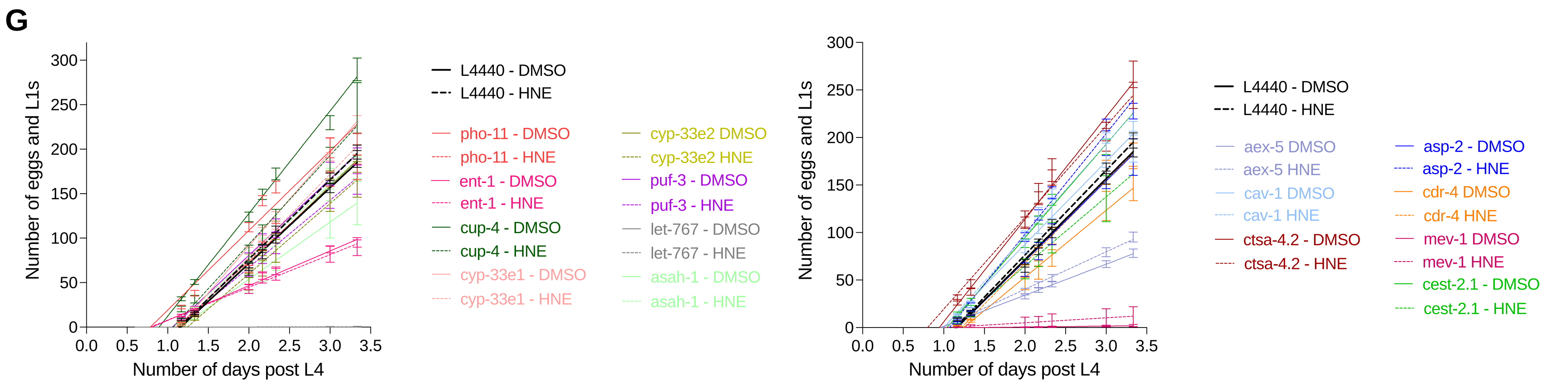


**F**

| Developmental stage | < -2σ | < -1σ | > +1σ | > +2σ | > +3σ |
|---|---|---|---|---|---|
| Day 1 post L4 (young adult) | - | cyp-33e1 | cyp-33e2<br>cest-2.1 | - | asp-2 |
| Day 3 post L4 (young adult) | - | let-767 | - | - | - |
| Day 5 post L4 (adult) | let-767 | aex-5<br>puf-3 | - | - | - |

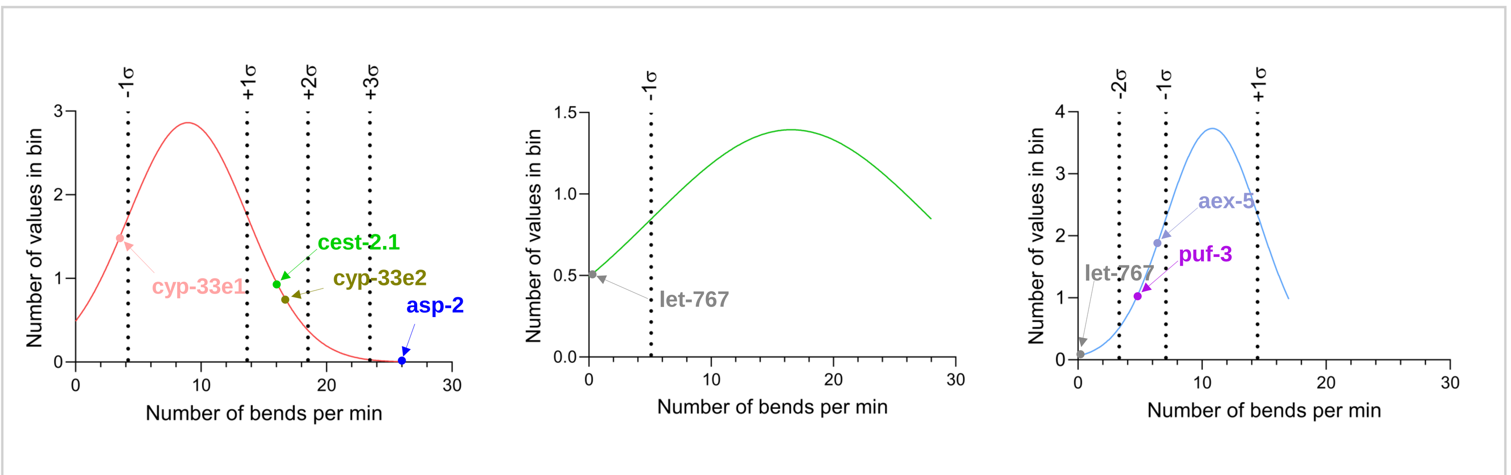


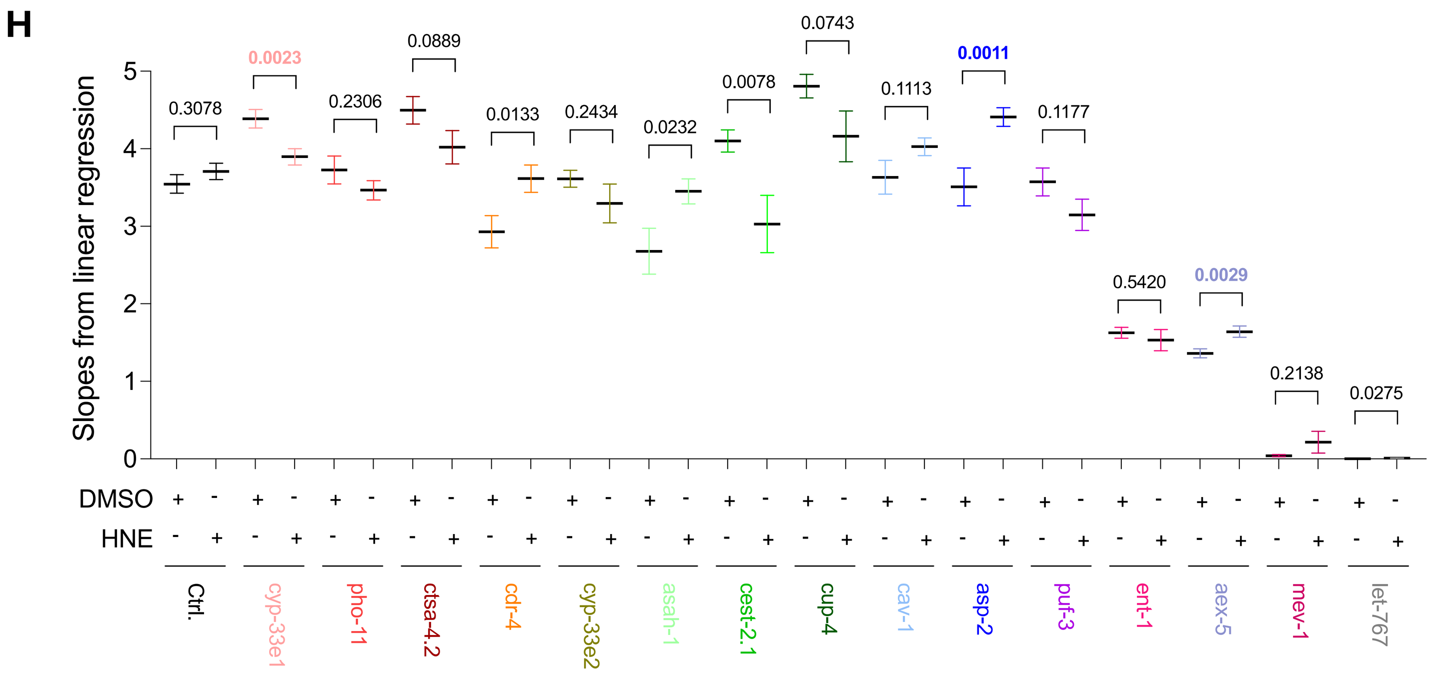


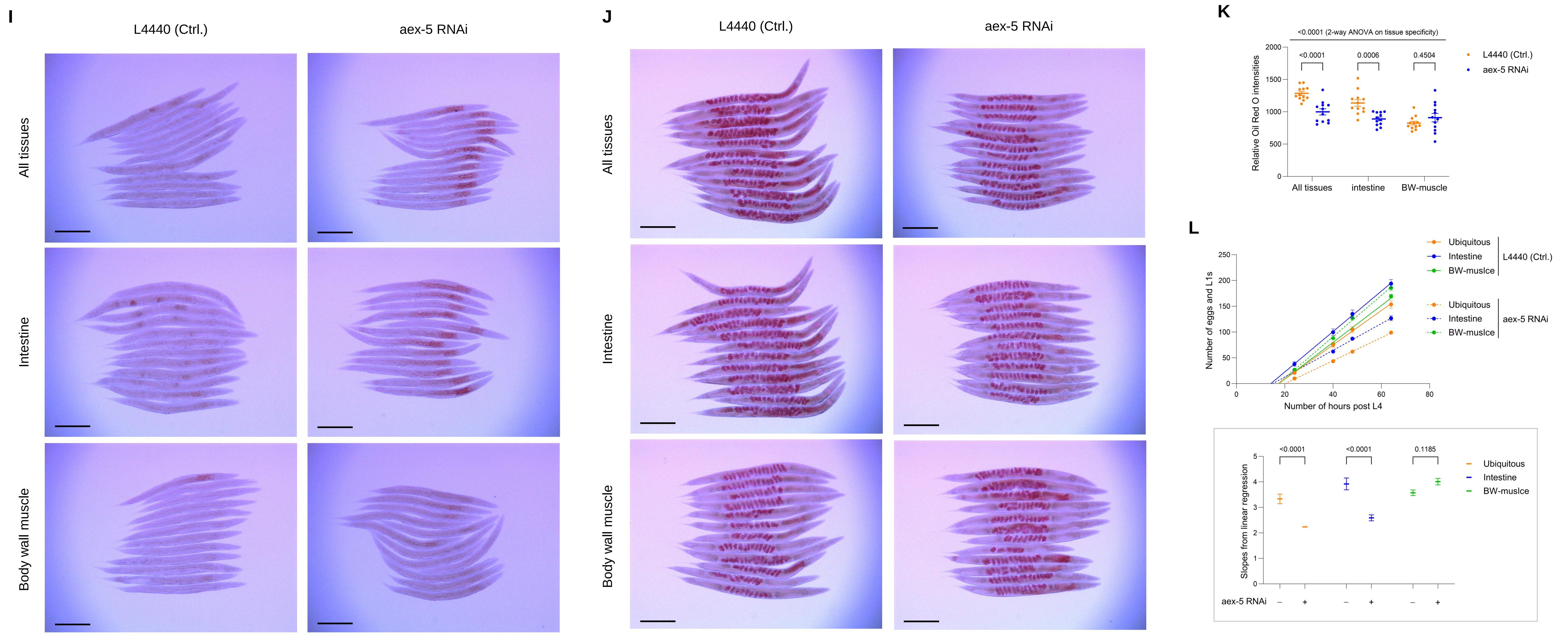

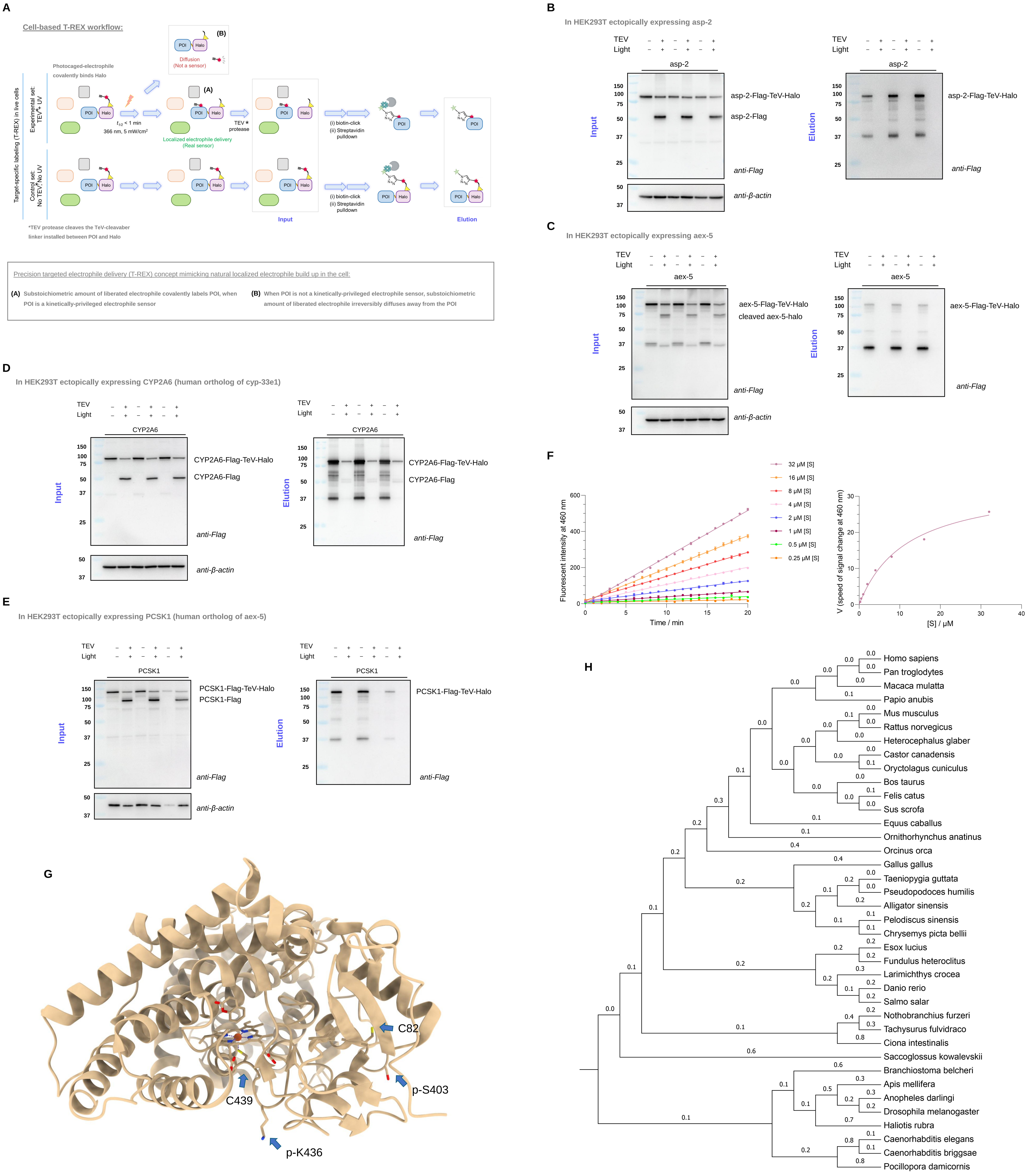
A
Cell-based T-REX workflow:
Photocaged-electrophile covalently binds Halo
Target-specific labeling (T-REX) in live cells
Experimental set: TEV, UV
Control set: No TEV, No UV
366 nm, 5 mW/cm²
TEV protease
Localized electrophile delivery (Real sensor)
Diffusion (Not a sensor)
(i) biotin-click
(ii) Streptavidin pulldown
POI
Halo
Input
Elution
*TEV protease cleaves the TeV-cleavable linker installed between POI and Halo
Precision targeted electrophile delivery (T-REX) concept mimicking natural localized electrophile build up in the cell:
(A) Substoichiometric amount of liberated electrophile covalently labels POI, when POI is a kinetically-privileged electrophile sensor
(B) When POI is not a kinetically-privileged electrophile sensor, substoichiometric amount of liberated electrophile irreversibly diffuses away from the POI
B
In HEK293T ectopically expressing asp-2
TEV
Light
asp-2
asp-2-Flag-TeV-Halo
asp-2-Flag
anti-Flag
anti-β-actin
Input
Elution
C
In HEK293T ectopically expressing aex-5
aex-5
aex-5-Flag-TeV-Halo
cleaved aex-5-halo
aex-5-Flag-TeV-Halo
D
In HEK293T ectopically expressing CYP2A6 (human ortholog of cyp-33e1)
CYP2A6
CYP2A6-Flag-TeV-Halo
CYP2A6-Flag
E
In HEK293T ectopically expressing PCSK1 (human ortholog of aex-5)
PCSK1
PCSK1-Flag-TeV-Halo
PCSK1-Flag
F
Fluorescent intensity at 460 nm
Time / min
32 μM [S]
16 μM [S]
8 μM [S]
4 μM [S]
2 μM [S]
1 μM [S]
0.5 μM [S]
0.25 μM [S]
V (speed of signal change at 460 nm)
[S] / μM
G
C82
C439
p-S403
p-K436
H
Homo sapiens
Pan troglodytes
Macaca mulatta
Papio anubis
Mus musculus
Rattus norvegicus
Heterocephalus glaber
Castor canadensis
Oryctolagus cuniculus
Bos taurus
Felis catus
Sus scrofa
Equus caballus
Ornithorhynchus anatinus
Orcinus orca
Gallus gallus
Taeniopygia guttata
Pseudopodoces humilis
Alligator sinensis
Pelodiscus sinensis
Chrysemys picta bellii
Esox lucius
Fundulus heteroclitus
Larimichthys crocea
Danio rerio
Salmo salar
Nothobranchius furzeri
Tachysurus fulvidraco
Ciona intestinalis
Saccoglossus kowalevskii
Branchiostoma belcheri
Apis mellifera
Anopheles darlingi
Drosophila melanogaster
Haliotis rubra
Caenorhabditis elegans
Caenorhabditis briggsae
Pocillopora damicornis

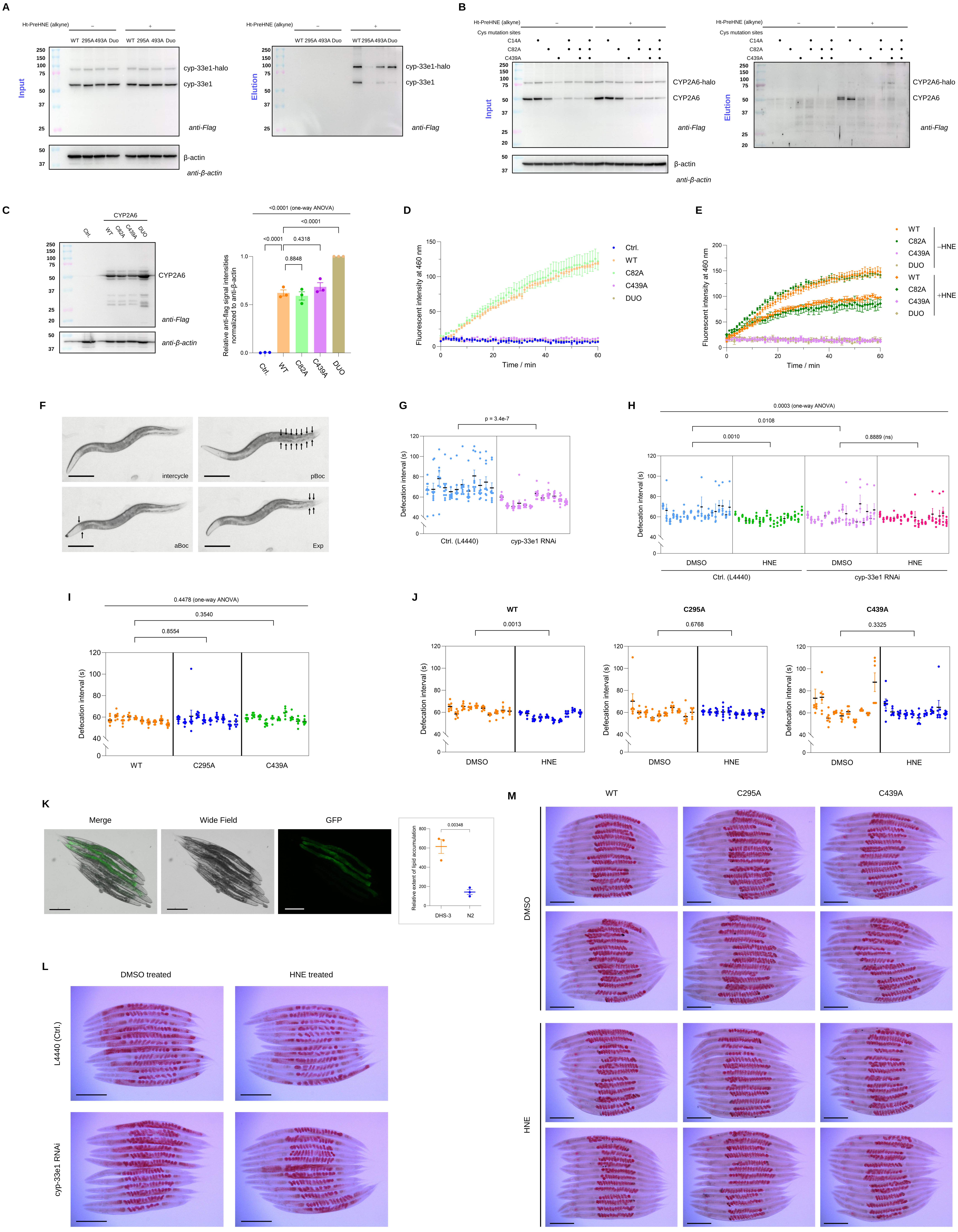
A
B
C
D
E
F
G
H
I
J
K
L
M
Input
Elution
anti-Flag
anti-β-actin
cyp-33e1-halo
cyp-33e1
CYP2A6-halo
CYP2A6
β-actin
Time / min
Fluorescent intensity at 460 nm
Defecation interval (s)
Ctrl. (L4440)
cyp-33e1 RNAi
DMSO
HNE
WT
C295A
C439A
Merge
Wide Field
GFP
DMSO treated
HNE treated
L4440 (Ctrl.)

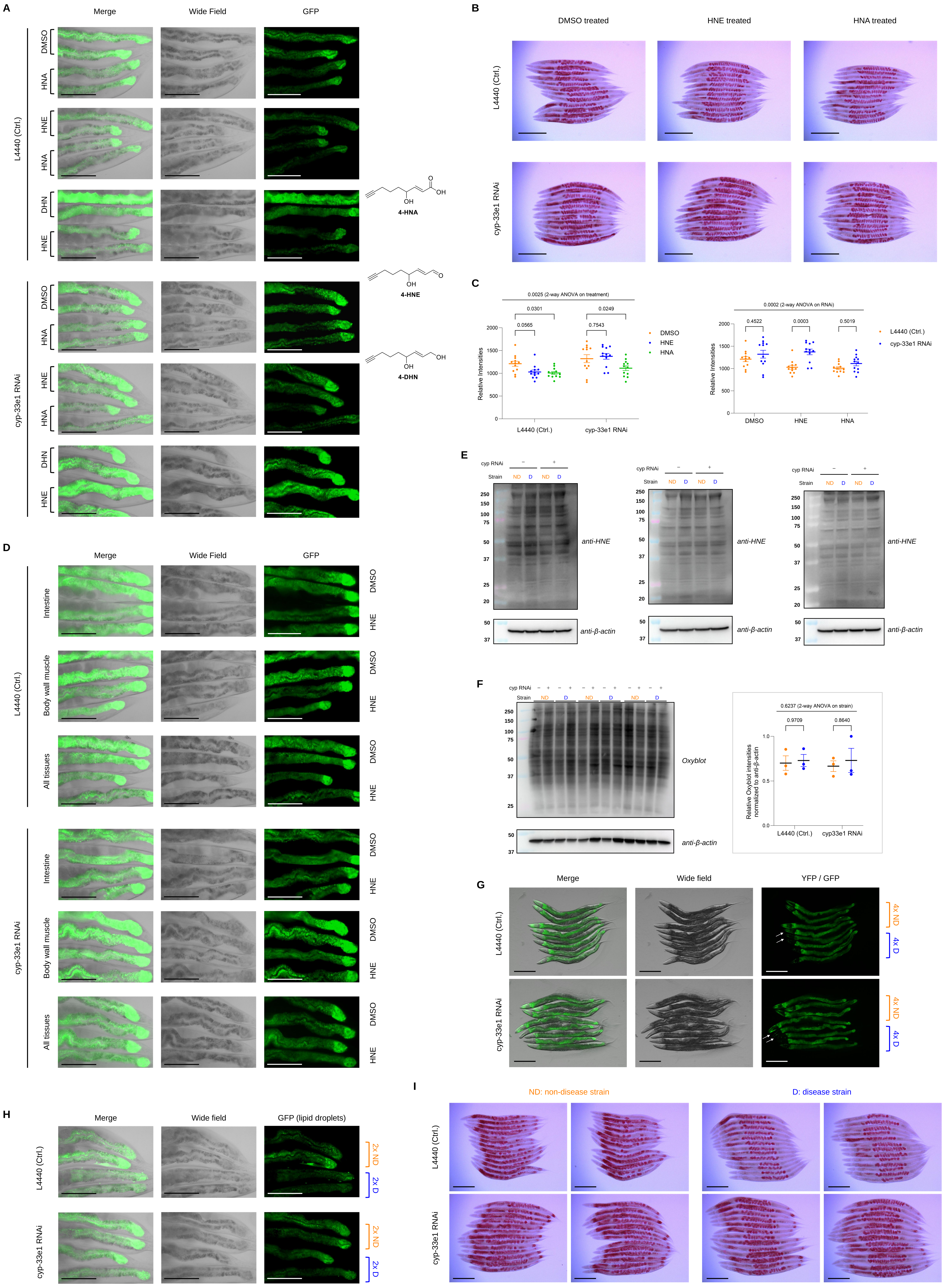
A
Merge
Wide Field
GFP
L4440 (Ctrl.)
cyp-33e1 RNAi
DMSO
HNA
HNE
DHN
4-HNA
4-HNE
4-DHN
OH
O
B
DMSO treated
HNE treated
HNA treated
C
0.0025 (2-way ANOVA on treatment)
0.0301
0.0565
0.0249
0.7543
Relative Intensities
DMSO
HNE
HNA
0.0002 (2-way ANOVA on RNAi)
0.4522
0.0003
0.5019
D
Intestine
Body wall muscle
All tissues
E
cyp RNAi
Strain
ND
D
anti-HNE
anti-β-actin
F
Oxyblot
0.6237 (2-way ANOVA on strain)
0.9709
0.8640
Relative Oxyblot intensities normalized to anti-β-actin
cyp33e1 RNAi
G
Wide field
YFP / GFP
4x ND
4x D
H
GFP (lipid droplets)
2x ND
2x D
I
ND: non-disease strain
D: disease strain
