## Supplementary material for "Organ-specific electrophile responsivity mapping in live *C. elegans*": SI figure legends

### SUPPLEMENTAL FIGURE LEGENDS

**Figure S1. OS-Localis-REX establishment and results, related to Figure 1**

**(A)** Assessments of transgene expression. Expression level of GFP-Halo transgene product in three organ-specific transgenic strains. 20 healthy L4-young adult transgenic *C. elegans* were selected and grown on nematode growth medium (NGM-agarose) plates (10 cm in diameter) for 4-5 days at 20 °C. Worms were then lysed and normalized lysates were subjected to western blot analysis using anti-Halo antibody. Representative data shown from n = 3 independent biological replicates. Samples from Bristol N2 wild-type strain were used for comparison. ***Inset:*** quantification (by ImageJ) where GFP-halo signals were normalized against β-actin loading control. See also **Figure 1A**.

**(B)** Schematic of chemistry underlying precision localized electrophile delivery platform, used to identify electrophile-actionable protein targets in specific organs of live *C. elegans*. See also **Figure 1B**.

**(C)** Immunofluorescence and Click-imaging validate the organ-specific anchorage of photocaged-electrophile probe [Ht-PreHNE(alkyne)] binding. Myo-2, ges-1, and myo-3 promoters respectively drive GFP-Halo expression in pharynx, intestine, and body-wall muscles expression, enabling the organ-specific photocaged-electrophile probe anchoring. Each image shows 3 respective age-synchronized organ-specific GFP-Halo-expressing hermaphrodites at day 1 adult stage with Ht-PreHNE(alkyne) treatment (3 lower worms), against 3 age-matched worms with Ht-PreHNE(no-alkyne) treatment (3 upper worms). Scale bars: 200 µm. ***Note:*** *these animals are mosaic, and hence GFP expression patterns are not identical across all GFP-Halo-transgenic worms in the same group.* See also **Method S1**.

**(D)** Zoomed images of **Figure S1C**. Scale bars: 100 µm. ***Note:*** *these animals are mosaic, and hence GFP expression patterns are not identical across all GFP-Halo-transgenic worms in the same group.*

**(E)** Assessments of minimum probe concentration to saturate Halo. Minimum Ht-PreHNE(alkyne) probe concentration required for saturating GFP-Halo active site in live *C. elegans* determined by TMR-Halo-ligand-blocking analysis. Ht-PreHNE(alkyne)-treated (6 h) or corresponding DMSO-treated live worms were subjected to lysis, followed by treatment of resulting lysate with TMR-Halo ligand (10 µM, 30 min), which labeled the Halo not bound by Ht-PreHNE(alkyne) in worms. Representative data are shown from n = 3 independent biological replicates. ***Inset:*** quantification (by ImageJ) of dose-dependent blockage of TMR-Halo-ligand binding, normalized against western-blot signals for total GFP-Halo and β-actin loading control. $EC_{50}^{\text{[Ht-PreHNE(alkyne) for saturating GFP-Halo in vivo]}}$ = 1.3 ± 0.2 µM for *myo-2p::gfp::halo* strain; 1.9 ± 0.3 µM for *ges-1p::gfp::halo* strain; 2.5 ± 0.3 µM for *myo-3p::gfp::halo* strain.

**(F)** Assessments of photouncaging efficiency. Time-dependent photo-uncaging of Ht-PreHNE(alkyne) in live *C. elegans*. Ht-PreHNE(alkyne)-treated (12 µM, 6 h) live worms were subjected to a low-power hand-held lamp (366 nm, 5 $mW/cm^2$) for indicated time. Representative data are shown from n = 3 independent biological replicates. ***Inset:*** quantification (by ImageJ) of time-dependent liberation of HNE(alkyne) from Halo, reported by Cy5-signal depletion, normalized against GFP-Halo signal. Data were fit to exponential curve: $t_{1/2}$ (photo-uncaging in vivo) = 0.2 ± 0.1 min for *myo-2p::gfp::halo* strain; 0.2 ± 0.1 min for *ges-1p::gfp::halo* strain; 0.2 ± 0.1 min for *myo-3p::gfp::halo* strain.

**(G)** OS-Localis-REX validation of selective enrichment of electrophile-responder proteins from indicated organ-specific Halo-transgenic strains. Representative coomassie-stained input and elution gels following biotin-streptavidin enrichment of electrophile(alkyne)-tagged proteins (and photocage-bound GFP-Halo) from *myo2p::gfp::halo, ges1p::gfp::halo* and *myo3p::gfp::halo* strains. Selective enrichment was observed in samples treated with both Ht-PreHNE(alkyne) and light-illumination, against two respective controls, namely, Ht-PreHNE(no alkyne) and light-illumination; Ht-PreHNE(alkyne) alone treatment. ***Inset:*** corresponding photocaged-probe structures. The signal of GFP-Halo band (~60 kDa) decreased in samples treated with Ht-PreHNE(alkyne) and light-illumination, as expected, as a result of photouncaging, leaving less amount of alkyne-functionalized photocaged-probe covalently bound to Halo. ~75 kDa band likely represents an endogenously-biotinylated protein, serving as an internal control.

**(H)** Validation of enrichment workflow for TMT6-plex target-ID of organ-specific electrophile responders by OS-Localis-REX. Left: Representative Coomassie-stained input (top) and elution (bottom) gels for organ-specific TMT6-plex mass spectrometry analysis from indicated Halo transgenic strains. TMT6-plex per each strain constituted 2 sets of conditions and 3 independent biological replicates, namely: Ht-PreHNE(no alkyne) and light treatment (triplicate control set); and Ht-PreHNE(alkyne) and light treatment (triplicate experimental set) (see also **Figure 1B**). Elution gel allows desalting prior to in-gel trypsin digest and TMT labeling. The thick bands in elution gel were streptavidin from high-capacity streptavidin agarose. Right: Same as the left one, except that samples were analyzed by western blot using anti-GFP and anti-β-actin antibodies. β-actin was used as the loading control. As expected, control groups did not enrich (no-alkyne)probe-bound-GFP-Halo.

**(I**) Selection criteria for quantitative ranking of organ-specific electrophile responders by OS-Localis-REX. TMT6-plex labeling and filtering workflow for organ-specific electrophile-responder target-ID. See **Figure 1B** for preceding experimental details performed in live *C. elegans*. Alkyne and no-alkyne designate corresponding photocaged-probes used (see ***inset***). For each replicate, hits (representing proteins with ≥2 unique peptides, protein FDR=0.01) were first ranked by TMT-ratiometric analysis against no-alkyne-probe-treated controls and these enrichment ratios (specifically, $log_2FC$) were fit to Gaussian distribution analysis. Hits with >2.58 $\sigma$ (corresponding to >99% confidence interval) from the mean, were combined from all 3 replicates per strain, and the resultant hits common to 3 replicates per strain were scored as significant hits. Also see **Figure 1C**; **Data S1** and **S2; Method S1**.

**(J)** Abundance profile of indicated OS-Localis-REX hits. OS-Localis-REX hits (see also **Figure 1D** and **Data S2**), mapped against whole-genome protein abundance in *C. elegans* (from PAXdb), reflect a wide-ranging native protein abundance across the electrophile-responders identified.

All data present mean±SEM. All sample sizes are listed in **Method S3**.

**Figure S2. OS-UltraID establishment and results, related to Figure 2**

**(A)** Expression, morphological, and viability validations of OS-Ultra-ID strains by live imaging. Fluorescence imaging, post levamisole immobilization of live worms, confirms organ-specific mCherry-P2A-UltraID expression: *myo-2*, *ges-1*, and *myo-3* promoters respectively drive mCherry-P2A-Flag-UltraID expression in pharynx, intestine, and body-wall muscles. Each image shows 3 respective age-synchronized organ-specific mCherry-P2A-Flag-UltraID-expressing hermaphrodites at day 1 adult stage (lower) against 3 age-matched wild-type N2 hermaphrodite worms (upper). Scale bars = 200 µm. ***Note:*** *these animals are mosaic, and hence mCherry expression patterns are not identical across all 3 transgenic worms in the same group.* The images within the first column here are reproduced in **Figure 2A**. See also **Method S1**.

**(B)** Total proteome abundance remains largely the same across the three OS-Ultra-ID strains. Lysates derived from each strain were subjected to label-free quantification (LFQ) MS-based target ID. See **STAR Methods** for details. (**a-c**) The resulting volcano plots show pair-wise comparison across the 3 strains. Only 3 proteins were found to be statistically-significantly different [FDR threshold = 0.05, $s_0 = 1$ for t-test = variance between groups/ (variance within groups + $s_0$), blue curves] and this was observed for comparison between pharynx (myo-2p)- and BW muscles (myo-3p)-specific Ultra-ID strains (middle). Ept-1 expression level was downregulated; and those of klp-15 and D1086.10 were upregulated, in pharynx-expressing strain, with respect to BW muscles-expressing strain (see 3 red dots in the middle plot). See also **Data S4**.

**(C)** Optimizations of Ultra-ID enzymatic activity in newly-generated OS-Ultra-ID strains. Streptavidin blot analyses of lysates originating from indicated strains, following treatment with indicated dosage of biotin over indicated time periods. The worms in these experiments were fed with MG1655:BioBKan (biotin-auxotrophic *E. coli*). The Bristol N2 (wild type) shows the endogenously biotinylated proteome background, to a lower extent than that in transgenic worms expressing active UltraID in the absence of exogenous biotin treatment. Inset on the right: in each case shows quantification of signals below 60 kDa MW marker within the corresponding blot (region marked by a representative dotted blue rectangle), normalized by anti-β-actin loading control. ***Note:*** complete solubility was not achievable at '(25)' mM biotin. Thus, 5 mM was considered the highest dosage in accurately reporting the dose-responsive biotinylation activity data. P-values were calculated by Dunnett's multiple comparisons test, post one-way ANOVA.

**(D)** Validations of OS-Ultra-ID transgenic strains and enrichment workflow for TMT11-plex target-ID of organ-specific high-abundant proteins. Biotinylated proteins were increased in indicated Ultra-ID transgenic (Tg) strains upon treatment with 5 mM of biotin for 4 hours. See **STAR Methods** for details. Worms were lysed and lysates were analyzed by streptavidin blots. The N2 (wild type) strain treated with biotin and the respective Tg strains without exogenous biotin treatment were included as controls. Due to the endogenous biotinylation in these animals (see main text discussions) and the high endogenous activity of UltraID, the signals in Tg strains (without biotin treatment) were higher than wild-type animals, despite the fact that all animals were fed with biotin-auxotrophic *E. coli*. The bands in N2 lanes represent endogenous biotinylated proteins in wild-type *C. elegans*. Quantified data for signals below 60 kDa MW marker (see dotted blue rectangle) of blots were normalized by anti-$\beta$-actin loading control. These observations guided us to choose biotin-treated N2 as more relevant controls for TMT-ratio calculations. See discussions in Main text, and also **Data S3** and **S5**.

**(E)** Workflow illustrating organ-specific proximity mapping (OS-Ultra-ID) in live *C. elegans*, quantitative mapping of organ-specific protein abundance. Animals were cultured from MG1655bioB:Kan (Biotin auxotroph *E. coli* strain). Live animals of selected strain were treated with either external biotin (*top row*, experimental set comprising independent biological triplicates) or buffer alone (*middle row*, 'Control 2', comprising independent biological triplicates). Bristol N2 worms were treated with identical time and concentration of external biotin for control set 1 (*bottom row*, 'Control 1', comprising independent biological triplicates). Following washout, the animals were lysed, and biotinylated proteins were enriched by streptavidin pulldown as described in **STAR Methods.** Subsequent to trypsin digest, the digested peptides were labeled isotopically by TMT11-plex reagents prior to LC-MS/MS analysis. See also **Figure S2D**.

**(F)** Setup and selection criteria for quantitative ranking of organ-specific high-abundant proteins by OS-Ultra-ID. TMT11-plex (only 9 types of TMT tags were used in this experiment due to 9 different groups: see manuscript text discussion for details) was deployed. For each replicate, hits (representing proteins with $\geq$ 1 unique peptide, protein discover FDR = 0.01) were first selected by setting an FDR $\leq$ 0.05. Afterwards, based on the TMT quantitative ratiometric analysis against the data resulting from wild-type (N2) biotin-treated strains as controls, either $\log_2$FC > 0 (all enriched proteins) or $\log_2$FC > 1 (two-fold significantly enriched protein) were further selected. See also **Figure 2A-C** and **Data S3** and **S5**.

**(G)** Unique and common hits mapped by OS-Localis-REX and OS-Ultra-ID across 3 organs. See **Figure S1I** for selection criteria of OS-Localis-REX and **Figure S2F** of OS-Ultra-ID. For OS-Localis-REX (*Top*), briefly, 11, 38, and 10 proteins were regarded as organ-unique and statistically-significant electrophile-responders derived from indicated organ-specific GFP-Halo transgenic strains; and 9, 5, and 2 proteins appear in two strains; and 7 proteins are electrophile-responders commonly responsive in all three strains. See also **Data S1** and **S2**. *Bottom left:* Based on $\log_2$FC > 0 (all enriched proteins) threshold and other criteria as defined in **Figure S2F**, 22, 205, 259 proteins are unique organ-specific high-abundant proteins derived from corresponding OS-Ultra-ID strains; and 14, 136, 27 targets appear in two strains; and 76 proteins are high-abundant proteins common to all three strains. *Bottom right:* Based on $\log_2$FC > 1 (all significantly enriched proteins) threshold and other criteria as defined in **Figure S2F**, 32, 107, 125 proteins are unique organ-specific high-abundant proteins derived from corresponding OS-Ultra-ID strains; and 9, 34, 14 targets appear in two strains; and 27 proteins are high-abundant proteins common to all three strains. See also **Data S5**.

All data present mean±SEM. All sample sizes are listed in **Method S3**.

**Figure S3. Bioinformatic analysis of ranked hits from OS-Localis-REX and OS-Ultra-ID, related to Figure 2**

**(A)** Functional enrichment analysis of ranked hits from OS-Localis-REX and OS-Ultra-ID, using g:Profiler. OS-Localis-REX hits shown in Table S2 were subjected to online software *g:Profiler*, using manufacturer's protocols[1] (https://bio.tools/gprofiler). A similar analysis was performed for OS-Ultra-ID datasets (**Data S5a-c**) by inputting the respective ranked protein list from each organ including all hits with fold-enrichment greater than 2. Insets in each correspond to output showing $p_{adj}$ in the order of table entries. The

resulting Manhattan plots from the enrichment analysis results were set to display: GO subterms (MF, molecular function; CC, cellular component; BP, biological process); and KEGG biological pathways. OS-Localis-REX hits derived from BW-muscle-specific Halo transgenic strain have no functional enrichment. See, URL below for general interpretations of the Manhattan plots, and Main Text for discussions.
https://biit.cs.ut.ee/gprofiler/page/docs.

**(B-E)** Network clustering and functional enrichment of ranked hits from OS-Localis-REX and OS-Ultra-ID, using STRING analysis. OS-Localis-REX hits (in **B**) shown in **Data S2** and OS-Ultra-ID hits (**C-E**, corresponding to **Data S5a-c**: those manifesting fold-enrichment greater than 2), were subjected to STRING analysis followed by MCL clustering, following manufacturer's protocols[2]. Default cut-off thresholds in Cytoscape (v3.10.1) were deployed following https://jensenlab.org/training/stringapp/ (exercise:3). For clarity, GO terms and KEGG pathways outputs are omitted within this figure, but terms identical to those shown in **Figure S3A**, were obtained as anticipated, and no overlapping terms were observed between hits from OS-Localis-REX and those from OS-Ultra-ID. (As in **Figure S3A**, OS-Localis-REX hits derived from BW-muscle-specific Halo transgenic strain have no functional enrichment). See Main Text for discussions.

**Figure S4. Functional relevance study of selected OS-Localis-REX hits, related to Figure 3**

**(A)** Seleted indicated OS-Localis-REX hits. 15 selected hits assayed for functional relevance in this study; their human orthologs and associated functions. Human orthologs in parentheses were identified using BLAST search. Others were established orthologs annotated in Wormbase. See also **Data S2**.

**(B)** RNAi knockdown in *C. elegans* validated by qRT-PCR. RNAi procedure and qRT-PCR analysis were performed as described in Methods. Relative mRNA abundance was normalized to β-actin. Horizontal dotted line indicates knockdown control (Ctrl), the value of which was arbitrarily set to 1.0. See **Method S1** for primer sequences.

**(C-D)** Effect of RNAi knockdown of individual hit proteins on animal survival. The viability assay and RNAi were performed as described in Methods. The resultant data comparing each trace from RNAi of each of the 15 genes, against knockdown-control (Ctrl.) (in **C**), were subjected to log-rank (Mantel-Cox) (**Figure 3A**) and Gehan-Breslow-Wilcoxon (in **D**) tests, respectively, uncovering in 4 and 5 targets, that have an effect on survival, as described in **Figure 3A**. See also illustrative chart in **Figure 2E**.

**(E)** Effect of RNAi knockdown of individual hit proteins on animal motility. The assay assessing the number of body bends made by worms subjected to indicated RNAi targeting 15 individual genes, or knockdown-control (Ctrl.), was performed as described in Methods. See also illustrative chart in **Figure 2E**. Data across all 15 genes (and Ctrl.) were normalized such that the highest body-bending rate corresponds to 100 in each case. **Figure 3B** selectively shows data corresponding to the 7 genes (out of these 15 shown in this supplemental figure) whose RNAi resulted in statistically-significant perturbations in motility, based on Gaussian distribution analysis (see **Figure S4F**).

**(F)** Gaussian distribution analysis of measured non-normalized body-bending rate results in the shown list of outstanding genes with indicated standard deviations, from the mean body-bending rate, against indicated developmental stage. Inset below shows Gaussian plots for non-normalized body-bending rate at, *Left:* day 1 post L4. *Middle*: day 3 post L4, and *Right*: day 5 post L4. Genes exhibiting $>1\sigma$ or $<-1\sigma$ were indicated in each plot.

**(G)** Effect of RNAi knockdown of individual hit proteins in HNE-treated vs. untreated animals on the efficiency of fertile eggs laid. Linear regression analysis of the number of fertile eggs over indicated days. **Figure 3B** selectively shows data corresponding to the 3 genes (out of these 15 shown in this supplemental figure) whose RNAi resulted in statistically-significant effects on egg-laying rate as a function of electrophile treatment.

**(H)** Slope values from linear regression in **Figure S4G**. P values are derived from unpaired two-tailed Students' t-test – error is standard error from linear fitting. See also illustrative chart in **Figure 2E**. The egg-laying assays in worms treated with RNAi of individual 15 genes, in the presence and absence of HNE, were performed as described in **STAR Methods**.

**(I)** Images of the Oil Red O stained L4 *C. elegans* subjected to aex-5 RNAi against RNAi control. 12 fixed age-synchronized L4 worms were Oil Red O stained and manually aligned in each image. Aberrant accumulation of lipids in the anterior intestine was observed only in the intestinal- or ubiquitous-aex-5-RNAi-treated groups, with respect to the corresponding RNAi-control groups. Scale bars: 200 µm.
**(J)** Images of the Oil Red O stained adult *C. elegans* subjected to aex-5 RNAi against RNAi control. 12 fixed age-synchronized adult hermaphrodites were Oil Red O stained and manually aligned in each image. In the intestinal- and ubiquitous-aex-5-RNAi-treated worms, the lipid level was decreased, and the worm sizes were smaller than control worms. Scale bars: 200 µm.
**(K)** The Oil Red O signals were quantified by circling the entire animal except for the pharyngeal region of adult worms (in **Figure S4J**). P-values were calculated by Šídák's multiple comparisons test in two-way ANOVA.
**(L)** Egg-laying rate of *C. elegans* altered in response to specific organ of aex-5 RNAi. Time-dependent rise in the total number of eggs in indicated groups subjected to three types of organ-specific or ubiquitous RNAi, against RNAi control (L4440). ***Inset***: Egg-laying rate of corresponding groups. In ubiquitous-RNAi and intestinal-RNAi groups, worms showed decreased egg-laying rates with respect to the corresponding RNAi-control groups.
All data present mean±SEM. All sample sizes are listed in **Method S3**.

**Figure S5. Cyp-33e1 and its human orthologs CYP2A6 are validated as kinetically-privileged responders of HNE by on-target and *in vitro* investigations, related to Figure 4**
**(A)** Workflow of target-specific electrophile labeling (T-REX) for validating the kinetically-privileged-electrophile-sensing ability of cyp-33e1 and its human ortholog, CYP2A6, in live HEK293T cells. Cultured HEK293T cells ectopically expressing corresponding Halo-fused protein-of-interest (POI) constructs as indicated in (**Figure 4A, S5B-E, S6A,** and **S6B**) were subjected to T-REX precision localized electrophile delivery (against samples not exposed to light, i.e., no electrophile release, as negative controls), followed by cell lysis, TeV-protease treatment (where applicable as indicated in **Figure 4A, S5B-E, S6A,** and **S6B**) that separates Halo and POI, and Click-biotin pulldown (see an overview workflow shown and also **STAR Methods** for details).
**(B-C)** Cyp-33e1 (but not asp-2 and aex-5) is a privileged electrophile sensor in live HEK293T cells. Following cell-based T-REX procedure and Click-biotin pulldown workflow (**Figure S5A**), input and elution samples were analyzed by western blot using anti-Flag antibody. Anti-β-actin antibody was used as loading control of the input samples. Each blot shows data from 3 independent biological replicates. See also **Figure 4A**. Predicted molecular weights of protein of interest (POI) and relevant constructs (MWs): POI-Flag-TeV-Halo-His: 84 kDa (asp-2); 96 kDa (aex-5); and POI-Flag (post TeV-protease cleavage): 49 kDa (asp-2); 61 kDa (aex-5). ***Note:*** Incomplete TeV cleavage results in residual full-length fusion protein in '+ TeV' lanes where applicable. Inadvertent cleavage occurs during the course of pulldown, leading to multiple Flag-positive bands in negative-control samples in Elution blot. Nonetheless, no HNE(alkyne)-tagged enriched species in experimental samples (+ TeV, + light) in Elution blot was observed. These data in combination with corresponding results from cyp-33e1 (**Figure 4A**), CYP2A6 (human ortholog of cyp-33e1, **Figure S5D**), and PCSK1 (**Figure S5E**), and in vitro HNEylation of CYP2A6 (**Figure 4B**), collectively validate kinetically-privileged electrophile-sensing propensity of cyp-33e1/CYP2A6, over other POIs assessed under identical assay conditions. See also **Figure 5A, 5B**, **S6A,** and **S6B; Data S3**.
**(D-E)** CYP2A6 (but not PCSK1), human ortholog of cyp-33e1, is a privileged electrophile sensor in live HEK293T cells. The experimental setup and data interpretations identical to those described above in **Figure S5B** and **S5C** legend were deployed, except that POI corresponds to CYP2A6 or PCSK1. PCSK1 is the human ortholog of aex-5. Predicted molecular weights (MWs): POI-Flag-TeV-Halo-His: 94 kDa (CYP2A6); 121 kDa (PCSK1); and POI-Flag (post TeV-protease cleavage): 59 kDa (CYP2A6); 86 kDa (PCSK1). ***Note:*** Incomplete TeV cleavage results in residual full-length fusion protein in '+ TeV' lanes where applicable. Inadvertent cleavage occurs during the course of pulldown, leading to multiple Flag-positive bands in negative-control samples in Elution blot. Nonetheless, the HNE(alkyne)-tagged enriched species in experimental samples (+ TeV, + light) in Elution blot in (**D**) corresponds to MW of CYP2A6-Flag. In (**E**) with PCSK1, no HNE(alkyne)-tagged enriched species in experimental samples (+ TeV, + light) was observed in Elution blot. These data in combination with corresponding results from cyp-33e1 (**Figure 3A**), asp-2, and aex-5 (**Figure S5B** and **S5C**), and in vitro HNEylation

of CYP2A6 (**Figure 3B**), collectively validate kinetically-privileged electrophile-sensing propensity of cyp-33e1/CYP2A6, over other POIs assessed under identical assay conditions. See also **Figure 5A, 5B**, **S6A,** and **S6B; Data S3**.

**(F)** Determination of $K_m$ of substrate, 3-cyano coumarin, for CYP2A6, human ortholog of cyp-33e1. Derivation of $K_m$ of the substrate, 3-cyano coumarin (see **STAR Methods** for details). _Left_: Progress curves of the CYP2A6-catalyzed fluorescent-product formation, as a function of indicated substrate concentrations. _Right_: the slopes from the progress-curve plot fit to linear regression, to yield enzymatic reaction velocities, that were subsequently plotted against substrate concentrations. Subsequent fit of the data to Michaelis-Menten equation $v = V_{max}*[S]/(K_M + [S])$ indicates: $K_M$ 12 ± 6 µM; $V_{max}$ 34 ± 7 a.u. $s^{-1}$. See also Figure **4C**, **5C**, **5D**, **S6D** and **S6E**.

**(G)** Ribbon representation of human CYP2A6 (PDB: 1Z10) featuring previously-reported canonical enzyme-mediated PTM sites and the two electrophile-sensing cysteines discovered in this study. See also **Figure 4D**. PhosphoSite+ database maps 9 different enzymatic PTMs (albeit noting that these were identified from large-scale proteomics datasets, with limited or no biochemical validations): phosphorylation at Ser4, Thr16, and Ser22 (the 3 residues not visible in crystal structure); Thr163, Thr295, and Ser403; dimethylation at Arg148 and Arg161; and acetylation at Lys436. 3D structural analysis shows Ser403 and Lys436 (shown in the figure) to be spatially proximal to the two newly-identified electrophile-sensing sites, C82 and C439, respectively.

**(H)** Phylogenetic analysis of cyp-33e1 and cysteine sites. Briefly, 38 representative metazoan species were first selected. All possible Cyp-orthologs in each species were analyzed with respect to sequences of cyp-33e1 and CYP2A6, using the NCBI BLASTp tool. The representative isoforms were selected based on the following parameters: E value, Query cover, and positive amino acid, in the BLAST output, to build the maximum likelihood tree[3,4] across different species based on the sequence alignment results. ***Note:*** sequence alignments of representative 38 isoforms, one from each species, were performed using MUSCLE[5]. Numbers reflect the distance between branches. See also **Figure 4D**.

**Figure S6. Cyp-33e1 and CYP2A6 sense HNE at two cysteines but the regulation of intestinal physiology depends on the C439 catalytical cysteine, related to Figures 5 and 6**

**(A)** Cell-based T-REX identifies kinetically-privileged electrophile sensing sites within cyp-33e1. T-REX-assisted evaluation of electrophile sensing ability of protein of interest (POI): cyp-33e1 WT; single; or double ('duo') cysteine-to-alanine mutants as indicated, were assayed in HEK293T cells ectopically expressing the corresponding Halo-Flag-(TeV)-POI constructs. Briefly, HNE(alkyne) is transiently made available in substoichiometric amounts within the proximity of indicated Halo-Flag-(TeV)-POI in live cells following established cell-based T-REX workflow, against indicated controls. Input and elution samples resulting from Click-biotin pulldown of the protein, post lysis of cells subjected to T-REX (versus indicated control), and subsequent TeV-protease-mediated separation of Halo and cyp-33e1 (WT or mutant), analyzed by western blot using anti-Flag and anti-β-actin antibodies. For a generic cell-based T-REX workflow, see **Figure S5A**. Predicted molecular weights (MWs): cyp-33e1-Flag-TeV-Halo-His: 95 kDa and cyp-33e1-Flag (post TeV-protease cleavage): 60 kDa. ***Note:*** Incomplete TeV cleavage resulted in residual full-length fusion protein in all lanes in both Input and Elution blots. See **Figure 5A** for corresponding quantification.

**(B)** Cell-based T-REX identifies kinetically-privileged electrophile sensing sites within CYP2A6. Experiment as in Figure S34 except that CYP2A6 and its indicated mutants replaced cyp-33e1 variants. See **Figure 5B** for corresponding quantification. Predicted molecular weights (MWs): CYP2A6-Flag-TeV-Halo-His: 94 kDa and CYP2A6-Flag (post TeV-protease cleavage): 59 kDa. ***Note:*** Incomplete TeV cleavage resulted in residual full-length fusion protein in all lanes in both Input and Elution blots. The quantified data (see **Figure 5B**) took into account differences in expression levels across WT and mutants.

**(C)** Western blot analysis and quantification of CYP2A6 (WT and indicated mutants) ectopically overexpressed in HEK293+Hycell expression system. ***Note:*** quantified data in **Figure 5C, 5D, S6D,** and **S6E** were adjusted for expression level variations as analyzed by western blot.

**(D)** Oxidoreductase activity tests and progress curve analyses of CYP2A6 (WT and mutants). Product formation (7-hydroxylation of 4-cyanocumarin) catalyzed by partially-purified microsomal CYP2A6 WT and indicated mutants against negative control was measured as described in **STAR Methods**. See also **Figure 5C** and **S6C**.
**(E)** Oxidoreductase activity tests and progress curve analyses of CYP2A6 (WT and mutants) in the presence and absence of HNE. Progress curve analyses of partially-purified human microsomal CYP2A6 (WT and indicated mutants) following treatment or no treatment with HNE (400 µM, 20 min pre-incubation) were undertaken as described in **STAR Methods**. See also **Figure 5D**.
**(F)** Representative images of wild-type N2 hermaphrodites undergoing Intercycle (animal at the stage in-between the DMP cycles) and 3 distinct motor steps of the DMP: pBoc, aBoc, and expulsion (Exp). See also **Movie 1** in **Data S6**. Briefly, following intercycle, defecation begins with posterior body muscle contraction (pBoc), with ensuing tail compression (furrowing, black arrows in pBoc) (0-8 s). After a short (~1 s) relaxation period, the anterior body muscle contraction (aBoc) occurs (10-11 s), with distinct pharynx movement (black arrows in aBoc), which was immediately followed by enteric muscle contraction (Emc) or expulsion (Exp), whereby the animal's internal pressure expels gut contents via anus opening (black arrows in Exp) (12-13 s). ***Note:*** The video was decelerated 0.5x. Scale bars: 200 µm.
**(G)** Cyp-33e1 regulates defecation motor program (DMP) cycles of *C. elegans*. Measured DMP intervals of age-synchronized Control-RNAi (left) or cyp-33e1-RNAi (right) worms at L4 stage (48 hours after seeding on RNAi plates). P values were calculated by the nested t-test. The nested t-test was taken because several defecation intervals were recorded from the same worm. Given this linkage between measurements, an unpaired two-tailed Students' *t*-test is not applicable. See also **Movie 2-3** in **Data S6**.
**(H)** Cyp-33e1's electrophile-sensing activity plays a functional role in regulating defecation motor program (DMP) cycles of *C. elegans*. Measured DMP intervals of age-synchronized Control-RNAi (left) or cyp-33e1-RNAi (right) worms following 1-h exposure to either electrophile (2 mM HNE in DMSO) or vehicle control (equivalent % vol of DMSO) and 16-h recovery (at day 1 adult stage). P-values were calculated by Tukey's multiple comparisons test, post nested one-way ANOVA. See also **Figure 5G**; **Movie 4-7** in **Data S6**.
**(I)** DMP changes are not ascribable uniquely to loss of cyp-33e1 activity. Measured DMP intervals of wild type (*left*), C295A knock-in (*middle*), or C439A knock-in (*right*) age-synchronized live worms at L4 stage (48 hours after seeding to RNAi plates). P-values were calculated by Dunnett's multiple comparisons test, post one-way ANOVA. See also **Movie 8-10** in **Data S6**. See also **Figure 5H**.
**(J)** Both electrophile-sensing cysteines within cyp-33e1 are functional regulators of stress-induced changes in *C. elegans* defecation motor program (DMP) cycles. Measured DMP intervals of wild type (*left*), C295A knock-in (*middle*), or C439A knock-in (*right*) age-synchronized live worms following 1-h exposure to 2 mM HNE(alkyne), or vehicle control (equivalent % vol of DMSO) and subsequent 16-h recovery (at day 1 adult stage). P values were calculated by nested t-test. See also **Movie 11-16** in **Data S6**. See also **Figure 5I**.
**(K)** Lipid-droplet reporter *dhs-3p::dhs-3::gfp* strain manifests a strong GFP signal against background autofluorescence in age-matched wild-type N2 animals. Representative images of 3 age-synchronized *dhs-3p::dhs-3::gfp* hermaphrodites at day 1 adult stage (top) and age-matched wild-type N2 worms (bottom), subsequent to levamisole immobilization. Scale bars: 200 µm. ***Inset***: quantification. The GFP signal was quantified by circling the intestinal region in both groups. P values were calculated by unpaired two-tailed Students' t-test.
**(L)** Images of Oil Red O stained worms subsequent to electrophile HNE treatment. 12 age-synchronized hermaphrodites at day 1 adult stage were fixed and Oil Red O-stained, and subsequently manually aligned for image acquisition. Scale bars: 200 µm. The Oil Red O signals were quantified by circling the entire animal except for the pharyngeal region. See also **Figure 6C** for quantification and analysis. ***Note:*** Images for RNAi-Control group are reproduced in **Figure 6C**. Alkyne-functionalized electrophile was used for consistency with Click-imaging-based assays in other datasets.
**(M)** Images of Oil Red O stained wild type and indicated knock-in worms subsequent to electrophile HNE treatment. 13 age-synchronized hermaphrodites at day 1 adult stage were fixed, Oil Red O-stained, and manually aligned and shown in each image. Scale bars: 200 µm. The Oil Red O signals were quantified by circling

the entire animal except for the pharyngeal region. See also **Figure 6D** and **6E** for quantification. Alkyne-functionalized electrophile was used for consistency with Click-imaging-based assay data elsewhere.
All data present mean±SEM. All sample sizes are listed in **Method S3**.

**Figure S7. Cyp-33e1 mediated metabolite (HNA) promotes lipid depletion in both external and internal HNE stress model animals, related to Figure 7**

**(A)** Images of *C. elegans* posterior intestine, following either DMSO, HNE, HNA, and DHN treatment in the presence of cyp-33e1 RNAi or RNAi control. 4 age-synchronized, adult *dhs-3p::dhs-3::gfp* hermaphrodites were aligned manually after levamisole immobilization. In every image, the top 2 worms and the bottom 2 worms were treated under indicated conditions. In L4440 (control) groups, HNA-treated worms, and HNE-treated worms, both showed decreased lipid-droplet signals, relative to the DMSO- and DHN-treated worms. In cyp-33e1-RNAi groups, only HNA-treated worms showed decreased lipid-droplet signals relative to DMSO-, DHN, and HNE-treated worms. Scale bars: 100 µm. See also **Figure 7A** for quantification. Alkyne-functionalized versions of the small-molecules were used for consistency with Click-imaging-based assay data elsewhere.

**(B)** Images of Oil Red O stained worms with either HNE or HNA treatment in the presence of cyp-33e1 RNAi or RNAi control. Representative images of Oil Red O stained age-synchronized hermaphrodites at day 1 adult stage, validating decreased lipid levels in HNA-treated worms in both RNAi control and cyp-33e1-RNAi groups. Consistent with other data elsewhere, HNE-treatment decreased Oil Red O signals in RNAi-control groups but not in cyp-33e1-RNAi groups. 12 fixed worms were manually aligned and shown in each image. Scale bars: 200 µm.

**(C)** Quantification of data in **Figure S7B**, presented in 2 different formats. Quantification of Oil Red O signals was performed by circling the entire animal except for the pharyngeal region to compare the effects of treatments. P-values were calculated by Dunnett's multiple comparisons test post two-way ANOVA. Alkyne-functionalized versions of the small-molecules were used for consistency with Click-imaging-based assay data elsewhere.

**(D)** Images of the *C. elegans* posterior intestine under organ-specific cyp-33e1 RNAi. 4 age-synchronized adult *dhs-3p::dhs-3::gfp* hermaphrodites at day 1 adult stage were aligned manually after levamisole immobilization. In every image, the top 2 worms and the bottom 2 worms were treated under indicated conditions. In L4440 (control) groups, all three strains (housing differential RNAi-machinery, see **Method S2D-G**) showed decreased lipid droplet levels following HNE treatment. By contrast, only intestinal-specific and ubiquitous-cyp-33e1-RNAi worms showed blockage of HNE-induced lipid depletion, whereas BW-muscle-specific cyp-33e1-RNAi worms failed to block HNE-induced lipid depletion. Scale bars: 100 µm. See also **Figure 7C** for the quantification results.

**(E)** Western blot analyses of HNEylated proteomes in *C. elegans* Huntington's diseased (D) strains and age-matched non-diseased (ND) animals subjected to cyp-33e1 RNAi or control RNAi. The higher degree of endogenous HNEylation was validated in *dhs-3p::dhs-3::gfp* x *unc-54p::htt513(Q128)::yfp* worms (diseased strain, **D**) using 3 independent biological replicates of anti-HNE western blots. See **Figure 7F** for quantification. The data from worms subjected to cyp-33e1 RNAi further reflect the higher levels of endogenous HNE selectively associated with L4440-fed (control, no RNAi) worms, but not with cyp-33e1-RNAi worms. P-values were calculated by Šidak's multiple comparisons test, post two-way ANOVA.

**(F)** Oxyblot™ analyses of HNEylated proteomes in *C. elegans* Huntington's diseased (D) strains and age-matched non-diseased (ND) animals subjected to cyp-33e1 RNAi or control RNAi. Contrasting the data in **Figure S7E** and **Figure 7D**, the levels of carbonylated proteomes are indifferent across *dhs-3p::dhs-3::gfp* x *unc-54p::htt513(Q128)::yfp* worms (diseased strain, **D**) and age-matched *dhs-3p::dhs-3::gfp* x *unc-54p::htt513(Q15)::yfp* worms (non-diseased strain, **ND**) either in the absence or present of cyp-33e1 RNAi. 3 independent biological replicates of oxyblots are shown. *Inset on the right*: quantification. P-values were calculated by Šidak's multiple comparisons test, post two-way ANOVA.

**(G)** Fluorescent images of doubly-Tg *C. elegans* Huntington's disease model strains (generated and validated as shown in **Method S2H**), in the presence of cyp-33e1 RNAi or RNAi control. Age-synchronized hermaphrodites of indicated strains were subjected to control RNAi (empty vector L4440) or RNAi(cyp-33e1). 4 adults encoding: *dhs-3p::dhs-3::gfp* x *unc-54p::htt513(Q15)::yfp* (upper 4 worms, non-disease strain, ND) and *dhs-3p::dhs-3::gfp* x *unc-54p::htt513(Q128)::yfp* worms (lower 4 worms, disease strain, D) were aligned manually after levamisole immobilization. White arrows illustrate the known aggregation of Huntington (Htt) protein selectively observed in **D** strain, but not in age-matched **ND** controls. See also **Figure 7D** and **S7H**. ***Note:*** YFP signal (fused to Htt protein) is selectively expressed in body-wall muscles, and thus it does not interfere with the lipid-droplet reporter GFP-signal in the intesting. This result was independently validated using Oil Red O staining (**Figure 7E**, **S7I**). Scale bars: 200 µm.

**(H)** Fluorescent images of doubly-Tg *C. elegans* Huntington's disease model strains (generated and validated as shown in **Method S2H**), in the presence of cyp-33e1 RNAi or RNAi control, zoomed into posterior intestinal regions. Experimental setup as in **Figure S7G**. 2 adults encoding: *dhs-3p::dhs-3::gfp* x *unc-54p::htt513(Q15)::yfp* (upper 2 worms, **ND** strain) and *dhs-3p::dhs-3::gfp* x *unc-54p::htt513(Q128)::yfp* worms (lower 2 worms, **D** strain) were aligned manually after levamisole immobilization. Animals were age-synchronized hermaphrodites at day 1 adult stage. Scale bars: 100 µm. See also **Figure 7D, 7E**, **S7G**, and **S7I**.

**(I)** Images of Oil Red O stained doubly-Tg *C. elegans* Huntington's disease model strains. 13 fixed age-synchronized Day-1 adult hermaphrodites encoding either *dhs-3p::dhs-3::gfp* x *unc-54p::htt513(Q15)::yfp* worms (non-disease strain, **ND**) or age-matched *dhs-3p::dhs-3::gfp* x *unc-54p::htt513(Q128)::yfp* worms (disease strain, **D**) that had been Oil Red O-stained were manually aligned and shown in each image. Scale bars: 200 µm. The Oil Red O signals were quantified by circling the entire animal except for the pharyngeal region. P-values were calculated by Šidak's multiple comparisons test, post two-way ANOVA. See also **Figure 7D**, **7E**, **S7G**, and **S7H**.

All data present mean±SEM. All sample sizes are listed in **Method S3**.
